# Subcytoplasmic location of translation controls protein output

**DOI:** 10.1101/2022.11.04.515216

**Authors:** Ellen L. Horste, Mervin M. Fansler, Ting Cai, Xiuzhen Chen, Sibylle Mitschka, Gang Zhen, Flora C. Y. Lee, Jernej Ule, Christine Mayr

## Abstract

The cytoplasm is highly compartmentalized, but the extent and consequences of subcytopIasmic mRNA localization in non-polarized cells are largely unknown. We determined mRNA enrichment in TIS granules (TGs) and the rough endopIasmic reticuIum (ER) through particle sorting and isolated cytosolic mRNAs by digitonin extraction. When focusing on non-membrane protein-encoding mRNAs, we observed that 52% have a biased transcript distribution across these compartments. Compartment enrichment is determined by a combinatorial code based on mRNA length, exon length, and 3′UTR-bound RNA-binding proteins. Compartment-biased mRNAs differ in the functional classes of their encoded proteins: TG-enriched mRNAs encode low-abundance proteins with strong enrichment of transcription factors, whereas ER-enriched mRNAs encode large and highly expressed proteins. Compartment localization is an important determinant of mRNA and protein abundance, which is supported by reporter experiments showing that redirecting cytosolic mRNAs to the ER increases their protein expression. In summary, the cytoplasm is functionally compartmentaIized by local translation environments.

## Introduction

In polarized cells such as neurons, intestinal epithelial cells, or cells of the early fly embryo, the majority of mRNAs have a distinct spatial localization pattern. ^1–5^ mRNA localization enables the local control of protein production and activity. ^6–8^ In non-polarized cells, mRNA localization has primarily been studied for membrane proteins, which mostly localize to the endoplasmic reticulum (ER) or to mitochondria. ^9–12^ Whereas the rough ER is established as major site of local protein synthesis for membrane and secretory proteins, ^9,10,13^ the cytoplasm is compartmentalized by additional membrane-bound and membraneless organelles. ^14–17^ Some of these compartments may enable the generation of unique biochemical translation environments, which have been suggested to be crucial for protein interaction partner selection during protein synthesis, ^16,18–20^ but it is currently largely unknown if the location of protein synthesis also matters for protein output.

TIS granules (TGs) represent one such unique translation compartment, which promotes the co-translational formation of protein complexes. ^16,19^ TGs are formed by the RNA-binding protein (RBP) TIS11B together with its bound mRNAs. ^16,21^ *TIS11B* mRNA is ubiquitously expressed, ^22^ suggesting that TGs are widespread. TGs are present under steady-state cultivation conditions and form a network-like structure that is intertwined with the rough ER. ^16,21^ To investigate the broader biological significance of TGs, we set out to determine the mRNAs enriched in TGs, the neighboring rough ER, and the surrounding cytosol.

As TIS11B protein is present in cells in two states (Fig. 1A), (i) as soluble protein in the cytosol, and (ii) as phase-separated TG network, ^16,21^ we decided to use fluorescent particle sorting ^23^ to identify mRNAs enriched in TGs. We also applied fluorescent particle sorting to isolate ER-enriched mRNAs and extracted cytosolic mRNAs using digitonin. We focused our analysis on mRNAs that encode non-membrane proteins and found more than 3600 mRNAs to be consistently enriched in one of the three compartments. mRNAs enriched in each compartment share similar mRNA architectures which differ substantially between compartments. Compartment-enriched mRNAs also differed significantly in production and degradation rates as well as in the functional classes and expression levels of their encoded proteins. TIS11B knockout (KO) and reporter experiments support a model by which a combinatorial code consisting of mRNA architecture features together with 3′UTR-bound RBPs, including TIS11B, TIA1/L1, and LARP4B, largely determines the compartment-biased mRNA localization pattern. Intriguingly, we observed that redirecting cytosolic mRNAs to the ER controls protein expression, which indicates that protein abundance regulation is spatially regulated in the cytoplasm.

**Figure 1.**
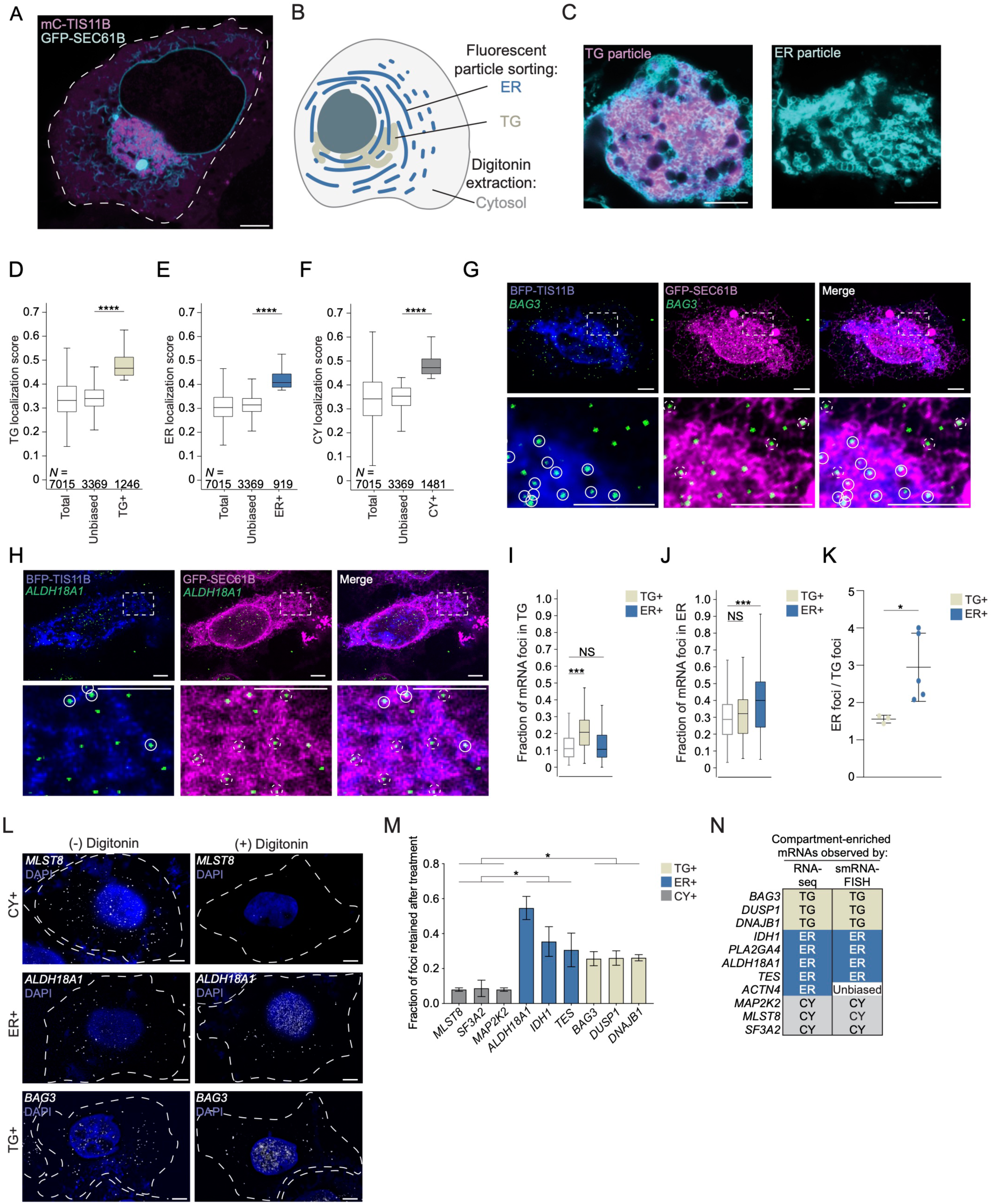
Strategy to identify compartment-enriched mRNAs. 1A. Confocal live cell imaging of HeLa cells after transfection of mCherry (mC)-TIS11B and GFP-SEC61B to visualize TGs and the rough ER. Scale bar, 5 µm. 1B. Schematic of a cell with three cytoplasmic compartments. 1C. As in (A) but showing fluorescent TG (left) and ER (right) particles. Scale bar, 5 µm. 1D. The transcript localization scores for the TG samples are shown for all mRNAs that encode non-membrane proteins (*N* = 7015), for the mRNAs defined as TG+ (*N* = 1246), and for mRNAs considered to have an unbiased transcript distribution (*N* = 3369) to illustrate that the relative transcript distribution between TG+ and unbiased mRNAs is substantially different. Mann Whitney test, *P =* 0. 1E. As in (D), but the transcript localization scores for the ER samples are shown for all mRNAs that encode non-membrane proteins (*N* = 7015), for the mRNAs defined as ER+ (*N* = 919), and for mRNAs considered to have an unbiased transcript distribution (*N* = 3369). Mann Whitney test, *P =* 1 × 10^−123^. 1F. As in (D), but the transcript localization scores for the cytosolic fractions are shown for all mRNAs that encode non-membrane proteins (*N* = 7015), for the mRNAs defined as CY+ (*N* = 1481), and for mRNAs considered to have an unbiased transcript distribution (*N* = 3369). Mann Whitney test, *P =* 0. 1G. SmRNA-FISH of endogenous TG+ mRNA *BAG3* (green) in HeLa cells. TIS granules (BFP-TIS11B, blue) and the ER (GFP-SEC-61B, magenta) were simultaneously visualized. Bottom panel shows a 5x zoom-in of the area indicated by dashed white box. White circles indicate colocalization of mRNA puncta with TGs, whereas dashed white circles indicate colocalization with the ER. Representative images are shown. Scale bar, 5 µm. 1H. As in (G), but smRNA-FISH of the endogenous ER+ mRNA *ALDH18A1* is shown. 1I. Quantification of TG-localizing smRNA-FISH foci of three TG+ and five ER+ endogenous mRNAs. The white box plot indicates the expected fraction of mRNA transcripts based on the TG compartment size, obtained from of 186 cells. mRNAs defined as TG+ are enriched in TGs (Mann Whitney test, ***, (TG), *P* = 5 × 10^−11^), whereas mRNAs defined as ER+ are not enriched. Additional images are shown in Fig. S2A-F and values for the individual mRNAs are shown in Fig. S2H. 1J. As in (I) but quantification of ER-localizing smRNA-FISH foci of three TG+ and five ER+ endogenous mRNAs. The white box plot indicates the expected fraction of mRNA transcripts based on the ER compartment size, obtained from of 186 cells. mRNAs defined as ER+ are enriched on the ER (Mann Whitney test, ***, (TG), *P* = 1 × 10^−6^), whereas mRNAs defined as TG+ are not enriched. Values for the individual mRNAs are shown in Fig. S2I. 1K. The ratio of smRNA-FISH foci colocalizing with the ER compared to the foci colocalizing with TGs is shown for the TG+ and ER+ endogenous mRNAs from (I) and (J). T-test for independent samples, *, *P* = 0.044. 1L. smRNA-FISH foci of endogenous mRNAs in HeLa cells before (−) and after (+) digitonin extraction. Cell boundaries are indicated by the dotted lines. Representative images are shown. Scale bar, 5 µm. 1M. Quantification of (L). Shown is the fraction of digitonin-resistant smRNA-FISH foci of endogenous mRNAs as mean ± std of three independent experiments. Number of cells analyzed, *MLST8*, *N* = 70; *SF3A2*, *N* = 67; *MAP2K2*, *N* = 48; *ALDH18A1*, *N* = 63; *TES, N* = 81; *IDH*, *N* = 127; *BAG3*, *N* = 187; *DUSP1, N* = 162; *DNAJB1, N* = 138. Additional images are shown in Fig. S3A-C. T-test for independent samples, *, *P* < 0.041. 1N. Summary of the smRNA-FISH validation for mRNAs defined as compartment enriched.

## Results

### Approach to determine subcytoplasmic mRNA localization

We set out to identify mRNAs that are localized in non-polarized human HEK293T cells under steady-state cultivation conditions. We focused on three major unenclosed cytoplasmic compartments—TGs, a condensate network formed by the RBP TIS11B, the cytosolic surface of the ER, and the soluble part of the cytoplasm known as the cytosol (Fig. 1B). For simplicity, we consider here the sum of the three compartments as the universe of cytoplasmic mRNAs.

To identify TG-enriched (TG+) and ER-enriched (ER+) mRNAs, we performed fluorescent particle sorting followed by RNA-seq. After co-transfecting cells with mCherry-TIS11B and GFP-SEC61B to label TGs and rough ER, respectively, we used mechanical lysis and differential centrifugation to isolate the cytoplasmic membrane fraction, followed by flow cytometry-based sorting of fluorescent particles (Fig. S1A, S1B). DAPI staining allowed us to identify and discard TG and ER particles that were still associated with nuclei. We used confocal microscopy to assess the purity of the particles. While ER particles generally did not contain mCherry-TIS11B, most of the TG particles contained GFP-SEC61B, consistent with the intimate association of TG and ER inside cells (Fig. 1C). Using western blot analysis, we observed that both particles contain similar amounts of ER proteins such as Calnexin and GFP-SEC61B, but TG particles contain 13-fold more mCherry-TIS11B than ER particles (Fig. S1C-E). As TG are defined by the presence of TIS11B, ^16^ we reasoned that the strong overrepresentation of TIS11B in TG particles would allow us to identify relative enrichments of mRNAs between the compartments. For subsequent RNA-seq experiments, we sorted TG particles from the TIS11B+SEC61B+ population and we sorted ER particles from the TIS11B−SEC61B+ population (Fig. S1B).

To isolate cytosolic mRNAs, we used digitonin extraction. ^24^ The extracted cytosol was not contaminated by nuclei or the ER, but it contained cytosolic proteins, including GAPDH which was used as a positive control (Fig. S1C). It also contained TIS11B, which was expected since soluble TIS11B is known to be present in the cytosol. We performed RNA-seq to determine the mRNA composition in the three fractions and focused our analysis on protein-coding mRNAs. The biological replicate samples of each compartment correlated well (Fig. S1F).

### mRNAs that encode membrane or secretory proteins largely localize to the ER membrane

We set out to investigate if the relative mRNA transcript distribution differs across the three compartments. For each gene, we determined a compartment-specific localization score. This score is calculated using the RPKM value obtained in each of the three compartments respectively and dividing it by the sum of the RPKM values in all three compartments. Thus, each gene is assigned three localization scores that correspond to the fraction of its transcripts localizing to each of the three compartments: TGs, the ER, and the cytosol.

It is accepted that most mRNAs that encode membrane or secretory proteins are translated on the ER. ^10,11,13^ To account for the difference in distribution, we plotted the localization scores of mRNAs that encode membrane/secretory proteins separately from the rest of the mRNAs that encode non-membrane proteins (Fig. S1G). In line with previous analyses, we find preferential partitioning of mRNAs encoding membrane/secretory proteins in the ER samples (Fig. S1G). ^10,11,13^ To examine if our compartment isolation method is valid, we compared it with datasets derived from three alternative isolation methods. ^9,11,13^ In our analysis, we consider 69% (*N* = 1,476) of membrane/secretory proteins to be enriched on the ER (Fig. S1H, Table S1). When comparing our results with mRNAs identified by biochemical fractionation, MERFISH or APEX-seq, we detected between 80-90% overlap and our data showed a quantitative relationship with data obtained by APEX-seq (Fig. S1I, S1J). ^9,11,13^ These results strongly support the validity of our purification strategy for mRNAs that encode membrane/secretory proteins.

### Half of mRNAs that encode non-membrane proteins have a biased transcript distribution in the cytoplasm

It was unclear whether non-membrane protein encoding mRNAs are biased in their localization. Not surprisingly, we found that these mRNAs have more evenly distributed localization scores across the three compartments (Fig. S1G). To faithfully compare absolute differences in mRNA distribution across the three compartments, the relative size of each compartment needs to be considered. However, this parameter is currently unknown. Therefore, instead, we calculated the relative enrichment of mRNAs within each compartment. We considered an mRNA compartment-enriched, if its mean localization score across biological replicates was at least 1.25-fold higher than the median localization score of the compartment samples (Fig. 1D-F). Based on this criterion, we identified 1246 TG+ mRNAs, 919 ER+ mRNAs, and 1481 mRNAs enriched in the cytosol (CY+), which were non-overlapping (Fig. 1D-F, Table S1). The remaining 3369 mRNAs were not enriched in a single compartment and were considered to have an unbiased localization pattern (Fig. 1D-F, Fig. S1K). Fig. 1D-F illustrate that the distribution of TG+, ER+, or CY+ localization scores are significantly different from the localization scores of mRNAs with unbiased localization patterns. Since localization scores across the three compartments sum to 1, an mRNA enriched in one compartment is relatively de-enriched in the other two (Fig. S1L). Based on this strategy, 52% of mRNAs that encode non-membrane proteins are significantly enriched in one of the three subcytoplasmic compartments in steady-state conditions.

As a recent study also analyzed the relative distribution of mRNA transcripts across subcellular compartments, we compared our compartment enrichment data with their results. ^25^ Although their dataset was generated by density gradient centrifugation in a different cell line, the two compartment enrichment datasets strongly agree in a qualitative and quantitative manner (Fig. S1M), suggesting that our isolation method as well as our strategy to define compartment-enriched mRNAs are valid. As non-membrane protein encoding mRNAs with biased transcript distributions in the cytoplasm have not been systematically characterized, we focused all subsequent analyses on mRNAs that encode non-membrane proteins.

### Validation of compartment-enriched mRNAs by single-molecule RNA-FISH

We further validated the mRNAs designated as compartment-enriched by performing single-molecule (sm) RNA-FISH on endogenous mRNAs (Table S2). ^26^ To distinguish between TG+ and ER+ mRNAs, we performed smRNA-FISH together with co-transfection of BFP-TIS11B and GFP-SEC61B to simultaneously visualize mRNA puncta, TGs, and the rough ER (Fig. 1G, 1H, Fig. S2A-G). Here, we considered an mRNA to have an unbiased localization pattern if its transcript distribution correlated with compartment size. As proxy for relative compartment size, we used the areas of the maximum projection of the fluorescent signals for each compartment and compared them to the whole cell area. Based on the compartment size distribution across 186 cells, for unbiased mRNAs, we expect that 11% of transcripts localize to TGs and 29% of transcripts localize to the ER (Fig. 1I, 1J).

For 3/3 TG+ mRNAs, we observed a significant enrichment of mRNA puncta in TGs, but not on the ER (Fig. 1G, 1I, 1J, Fig. S2A, S2B, S2H, S2I). For the five ER+ mRNAs tested, the mRNA puncta of 4/5 mRNAs were significantly enriched on the ER and for all five, we observed a 2-4-fold higher fraction of mRNA puncta that co-localized with the ER compared to TGs (Fig. 1H-K, Fig. S2C-I).

Cytosolic mRNAs were isolated through digitonin extraction. This means that CY+ mRNAs localize to the soluble part of the cytoplasm and are not attached to cytoplasmic structures, including membranes or the cytoskeleton. As steady-state smRNA-FISH can only inform on co-localization and not attachment, we validated CY+ mRNAs by performing smRNA-FISH before and after digitonin extraction and calculated the fraction of retained mRNAs. We observed significantly greater retention of both TG+ and ER+ mRNAs compared to CY+ mRNAs, which were depleted by about 90% following digitonin treatment (Fig. 1L, 1M, Fig. S3A-C). This confirms that CY+ mRNAs predominantly localize to the soluble part of the cytoplasm. Taken together, as we successfully validated 10/11 mRNAs that were designated to be TG+ or ER+ or CY+ (Fig. 1N), we conclude that about half (52%) of mRNAs encoding non-membrane proteins are enriched in distinct subcytoplasmic compartments.

### mRNA and protein levels strongly correlate with the location of translation

Next, we characterized the features of compartment-enriched mRNAs and found substantial differences in their steady-state mRNA and protein levels (Fig. 2A, 2B, Fig. S4A, S4B). We observed that TG+ mRNAs have the lowest steady-state expression levels and encode proteins with the lowest expression levels (Fig. 2A, 2B). To examine if the low mRNA levels are caused by high mRNA degradation rates, we estimated mRNA half-lives by analyzing Precision Run-On sequencing (Pro-seq) and RNA-seq data (Fig. 2C, 2D, Fig. S4C-E). ^27,28^ Pro-seq values can be treated as transcription rates and RNA-seq data can be viewed as a measure of RNA concentration to estimate RNA decay rates required for a steady-state equilibrium. ^28^ For TG+ mRNAs, we observed that the low steady-state mRNA levels were not primarily caused by a low mRNA stability. Instead, these mRNAs had the lowest transcription rates, suggesting that these mRNAs are either produced at a low rate or have high cotranscriptional degradation rates (Fig. 2C, 2D, Fig. S4D, S4E). ^29^ CY+ mRNAs had the highest degree of mRNA turnover with both high production and degradation rates (Fig. 2C, 2D). ER+ mRNAs encode proteins with the highest expression levels, particularly when normalizing to their intermediate steady-state mRNA levels (Fig. 2A, 2B).

**Figure 2.**
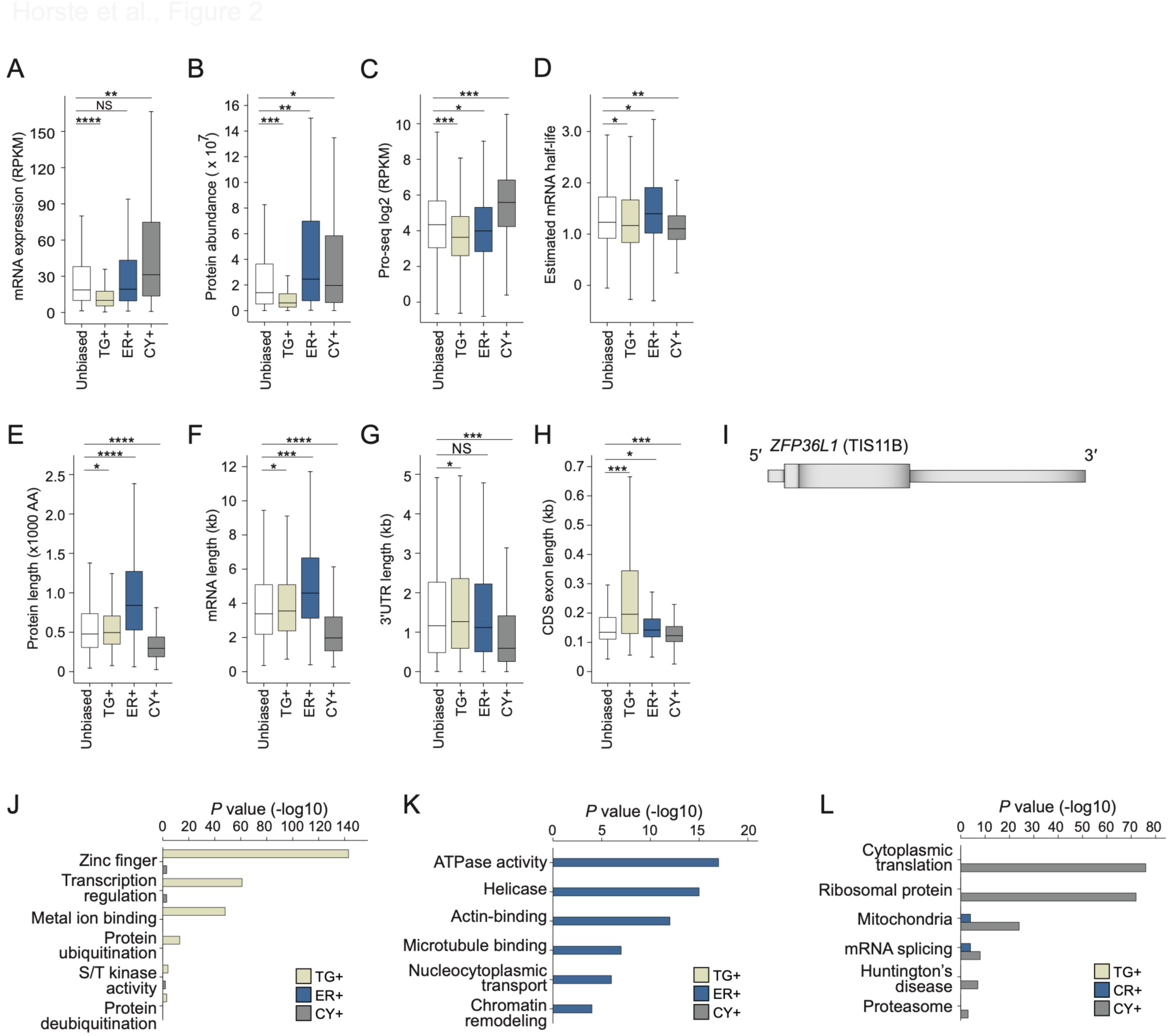
Characteristics of compartment-enriched mRNAs. 2A. Steady-state mRNA abundance levels obtained from whole cell lysates. TG+, *N* = 1246; ER+, *N* = 919, CY+, *N* = 1481; unbiased, *N* = 3369. Mann Whitney test: *, 0.05 >*P>* 1 × 10^−9^; **, 1 × 10^−10^ >*P>* 1 × 10^−20^; ***, 1 × 10^−21^ >*P>* 1 × 10^−80^; ****, 1 × 10^−81^ >*P>* 0. Exact *P* values are listed in Table S3. RPKM, reads per kilobase of transcript per million reads mapped. 2B. As in (A), but steady-state protein levels obtained from whole cell lysates are shown. TG+, *N* = 469; ER+, *N* = 638; CY+, *N* = 833; unbiased, *N* = 2001. 2C. As in (B), but Pro-seq levels are shown, which indicate transcription rates. TG+, *N* = 1222; ER+, *N* = 896; CY+, *N* = 1425; unbiased, *N* = 3268. 2D. As in (C), but estimated mRNA half-lives are shown. 2E. As in (A), but protein size distributions are shown. AA, amino acid 2F. As in (A), but mRNA length distributions are shown. 2G. As in (A), but 3′UTR length distributions are shown. 2H. As in (A), but average CDS exon length distributions are shown. 2I. Gene model of *ZFP36L1* (TIS11B) showing its unusual CDS exon length distribution. Tall boxes indicate CDS exons and the narrow boxes indicate the 5′ and 3′UTRs. 2J. Gene ontology analysis for TG+ mRNAs. Shown are the top six functional gene classes and their Benjamini-Hochberg adjusted *P* values for categories that are significantly and uniquely enriched in TG+ mRNAs. The Benjamini-Hochberg adjusted *P* values for the same categories for ER+ and CY+ mRNAs are shown for comparison. 2K. As in (J), but for ER+ mRNAs. 2L. As in (J), but for CY+ mRNAs.

In addition, we observed that the compartment-enriched mRNAs differed substantially in their gene architectures (Fig. 2E-H, Fig. S4F-K). ER+ mRNAs encode the largest proteins with a median size of 840 amino acids, nearly three-times larger than proteins encoded by CY+ mRNAs (Fig. 2E). The difference in protein size was reflected in the large differences in exon number and mRNA length between ER+ and CY+ mRNAs (Fig. 2F, Fig. S4J, S4K). The median length of ER+ mRNAs is 4600 nucleotides (nt), whereas the median length of CY+ mRNAs is 2000 nt. Not surprising, CY+ mRNAs also have the shortest 3′UTRs (Fig. 2G). TG+ mRNAs are uniquely characterized by large coding sequence (CDS) exons with a median size of 200 nt, compared to a median exon size of 133 nt for the remaining mRNAs (Fig. 2H). Further analysis revealed that the majority of TG+ mRNAs have a gene architecture that is similar to the gene architecture of *ZFP36L1* (encoding TIS11B), which is characterized by a short first exon and a long last exon that contains ∼95% of its coding sequence (Fig. 2I).

Moreover, compartment-enriched mRNAs encode substantially different functional gene classes. ^30^ Consistent with the low steady-state protein expression levels, TG+ mRNAs were strongly enriched in proteins containing zinc fingers and in transcription factors, which are known to have low expression (Fig. 2J). ^31^ In contrast, ER+ mRNAs encode large and highly abundant proteins, and they were enriched in helicases, cytoskeleton-binding proteins, and chromatin regulators (Fig. 2K). CY+ mRNAs often encode smaller proteins involved in the regulation of translation or splicing (Fig. 2L).

### TGs support active translation

TIS granules may constitute a specialized translation environment for nuclear proteins that require low expression levels, such as transcription factors (Fig. 2A, 2B, 2J). ^31^ In order to provide evidence for active translation in TIS granules, we applied the SunTag system to simultaneously visualize mRNAs and their nascent proteins in TGs and in the cytosol (Fig. S3D, S3E). ^32^ We confirmed that TGs represent a translation environment for mRNAs. ^16,19^ We observed that the number of mRNA foci in TGs was five-fold lower compared to the cytosol. However, the proportion of mRNA translated was similar in TGs and the cytosol (Fig. S3F, S3G). Taken together, our data show that TGs are sites of active translation and that the low expression level of TG-translated proteins is predominantly a result of their low nuclear gene expression (Fig. 2A, 2C).

### Differential 3′UTR binding of several RBPs correlates with compartment enrichment of mRNAs

Our next goal was to identify the RBPs responsible for mRNA enrichment in the three compartments (Fig. 1D-F). As TIS11B is the scaffold protein of TGs, ^16^ we performed iCLIP of TIS11B in HEK293T cells (Fig. S5A, S5B). We confirmed that the top binding motif of TIS11B in 3′UTRs of mRNAs is the canonical AU-rich element (UAUUUA) (Fig. S5C). To perform a comprehensive analysis on localization regulators, we included additional CLIP datasets. ^33,34^ Altogether, we analyzed CLIP data from 170 RBPs and found that 24 of them showed an enrichment of binding sites in 3′UTRs of mRNAs belonging to transcripts that preferentially localize to one of the three compartments (Table S4). We applied logistic regression and identified seven RBPs whose binding contributed most significantly to mRNA enrichment in the three compartments. They include TIS11B, HuR, PUM2, HNRNPC, TIA1/L1, LARP4B and METAP2 (Fig. 3A). As a previous CLIP anaIysis showed that peaks for TIA1 and TIAL1 cannot be distinguished, ^35^ we used the sum of peaks from TIA1 and TIAL1 to obtain the values for TIA1/L1. According to this regression analysis, the presence of TIS11B, HuR, PUM2, and HNRNPC on mRNAs correIates with TG enrichment, the presence of TIA1/L1 correIates with ER enrichment, and the presence of LARP4B or METAP2 correlates with cytosoI enrichment (Fig. 3A).

**Figure 3.**
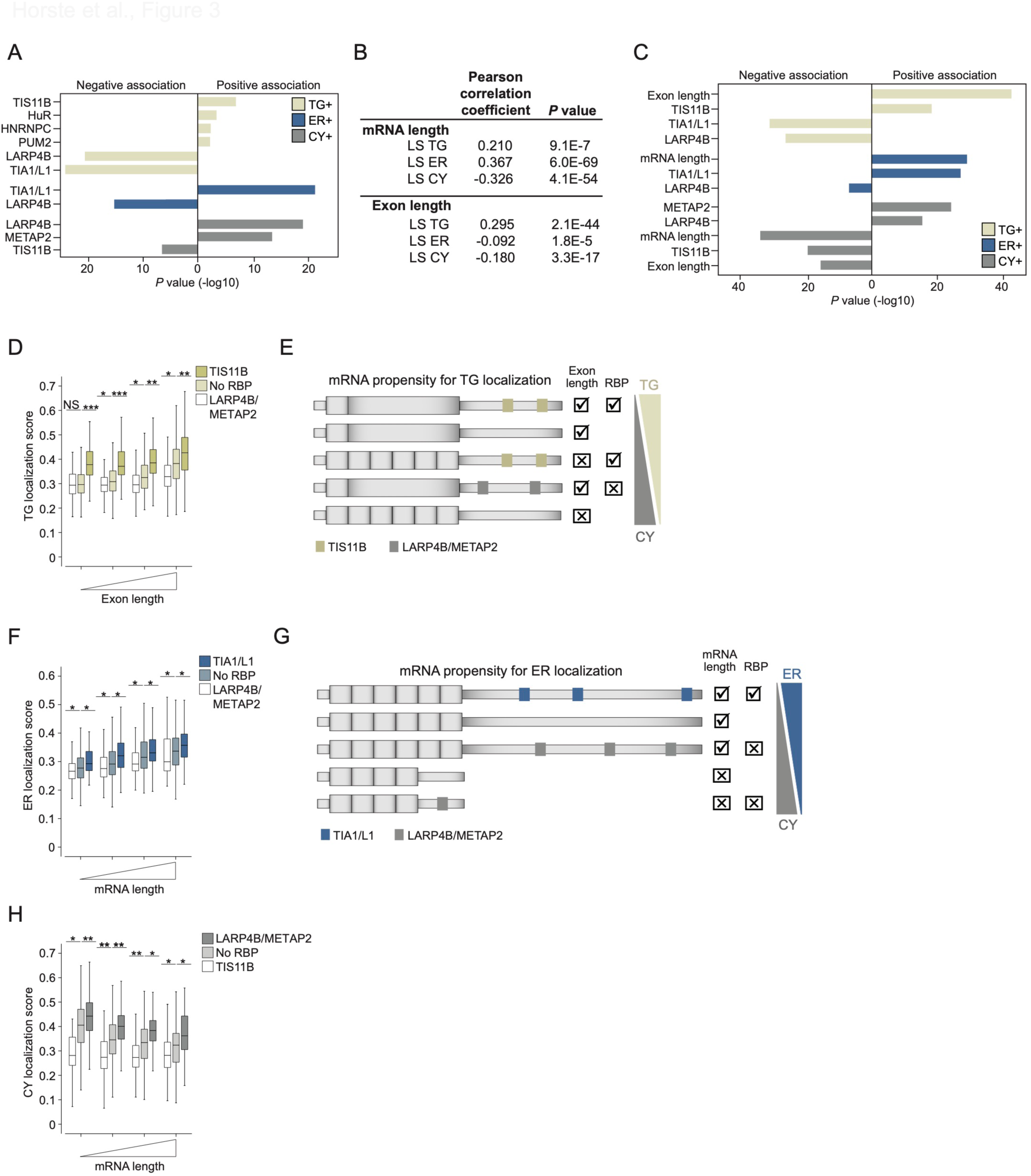
mRNA architecture features together with RBPs determine the subcytoplasmic transcript distribution. 3A. 3′UTR-bound RBPs that are positively or negatively associated with compartment-enriched mRNAs. Shown are the -log10 transformed *P* values obtained from logistic regression (see Table S4). 3B. Pearson’s correlation coefficients of mRNA and exon length with compartment localization scores (LS). 3C. As in (A) but integrating 3′UTR-bound RBPs from (A) and the mRNA architecture features ‘mRNA length’ and ‘CDS exon length’. 3D. The propensity to localize to TGs is shown for mRNAs stratified by exon length and bound RBPs. No RBP (*N* = 1498), bound only by LARP4B or METAP2 (*N* = 717) or only by TIS11B (*N* = 834). Mann Whitney test was performed. *P* value categories as in Fig. 2A. 3E. Model showing the additive effects of exon length and RBPs on the mRNA localization propensity to TGs or the cytosol. RBPs can have positive (check) or negative (x) effects. Shown as in Fig. 2I. 3F. As in (D) but shown is the propensity of mRNAs to localize to the ER, stratified by mRNA length and the bound RBPs. Bound only by TIA1/L1 (*N* = 634). 3G. As in (E) but showing the additive effects of mRNA length and RBPs on the mRNA localization propensity to the ER or the cytosol. RBPs can have positive or negative effects. 3H. As in (F) but shown is the propensity of mRNAs to localize to the cytosol. mRNAs bound only by LARP4B or METAP2 (*N* = 717) or only by TIS11B (*N* = 834).

### mRNA architecture features together with RBPs generate a combinatorial code for subcytoplasmic transcript distribution

As our analysis revealed that 2154 mRNAs (30.7%) that encode non-membrane proteins were not bound by any of the seven RBPs (Fig. S5D), we considered additionaI regulatory factors for mRNA localization. Among these mRNAs, we observed that mRNA Iength correIated strongIy with the ER and CY IocaIization scores, but in opposite directions, suggesting that long mRNAs are associated with ER localization (Fig. 3B). Similarly, average CDS exon Iength correIated strongIy and in a positive manner with the TG IocaIization score, but negatively with the CY IocaIization score (Fig. 3B).

When including mRNA and exon length in the logistic regression, we observed that mRNA architecture features contribute strongly to mRNA enrichment in each of the three compartments (Fig. 3C, Table S4). Our data indicate that the propensity of an mRNA for compartment localization depends on both architectural features and the presence of 3′UTR-bound RBPs. To better understand the rules for mRNA localization to the compartments, we plotted the propensity for TG enrichment and integrated RBPs that were positively or negatively associated with TG enrichment together with exon length (Fig. 3D). Regardless of exon length, binding of LARP4B/METAP2 decreased the TG localization score (Fig. 3D, 3E). For mRNAs with average exon length, TIS11B binding strongly increased their localization propensity to TGs. A similar TG localization propensity was observed for mRNAs with long exons that were not bound by TIS11B. The strongest TG enrichment was found for mRNAs with long exons that were bound by TIS11B, indicating that TIS11B and exon length have additive effects (Fig. 3D, 3E).

The two features that best promote mRNA localization to the ER are mRNA length and 3′UTR-bound TIA1/L1 (Fig. 3F, 3G). Short mRNAs increase their ER IocaIization propensity by the presence of TIA1/L1. Long mRNAs that are not bound by TIA1/L1 have a similar propensity for ER IocaIization, which can be further increased by the presence of TIA1/L1 (Fig. 3F, 3G). In contrast, shorter mRNAs not bound by any RBP or that are bound by LARP4B/METAP2 have a high propensity to localize to the cytosolic fraction (Fig. 3G, 3H). Taken together, our data suggest the existence of a combinatoriaI code, that integrates mRNA and exon length with the presence of RBPs, to determine subcytoplasmic mRNA localization (Fig. 3E, 3G).

### mRNA transcript distribution changes upon TIS11B deletion

Next, we set out to experimentally test the proposed mRNA localization code by investigating the influence of TIS11B and TIA1/L1 on subcytoplasmic mRNA transcript distribution. We generated HEK293T cells with an inducible knockout (KO) of TIS11B, isolated ER particles and extracted the cytosol (Fig. S5E, S5F, Table S5). To examine where mRNAs designated as TG+ localize in the absence of TGs, we identified the top 20% of mRNA localization changes to the ER and the cytosol and intersected them with mRNAs designated as TG+ (Fig. 4A). As only two compartments were isolated, increased mRNA localization to the ER means decreased cytosolic localization and vice versa (Fig. 4A).

**Figure 4.**
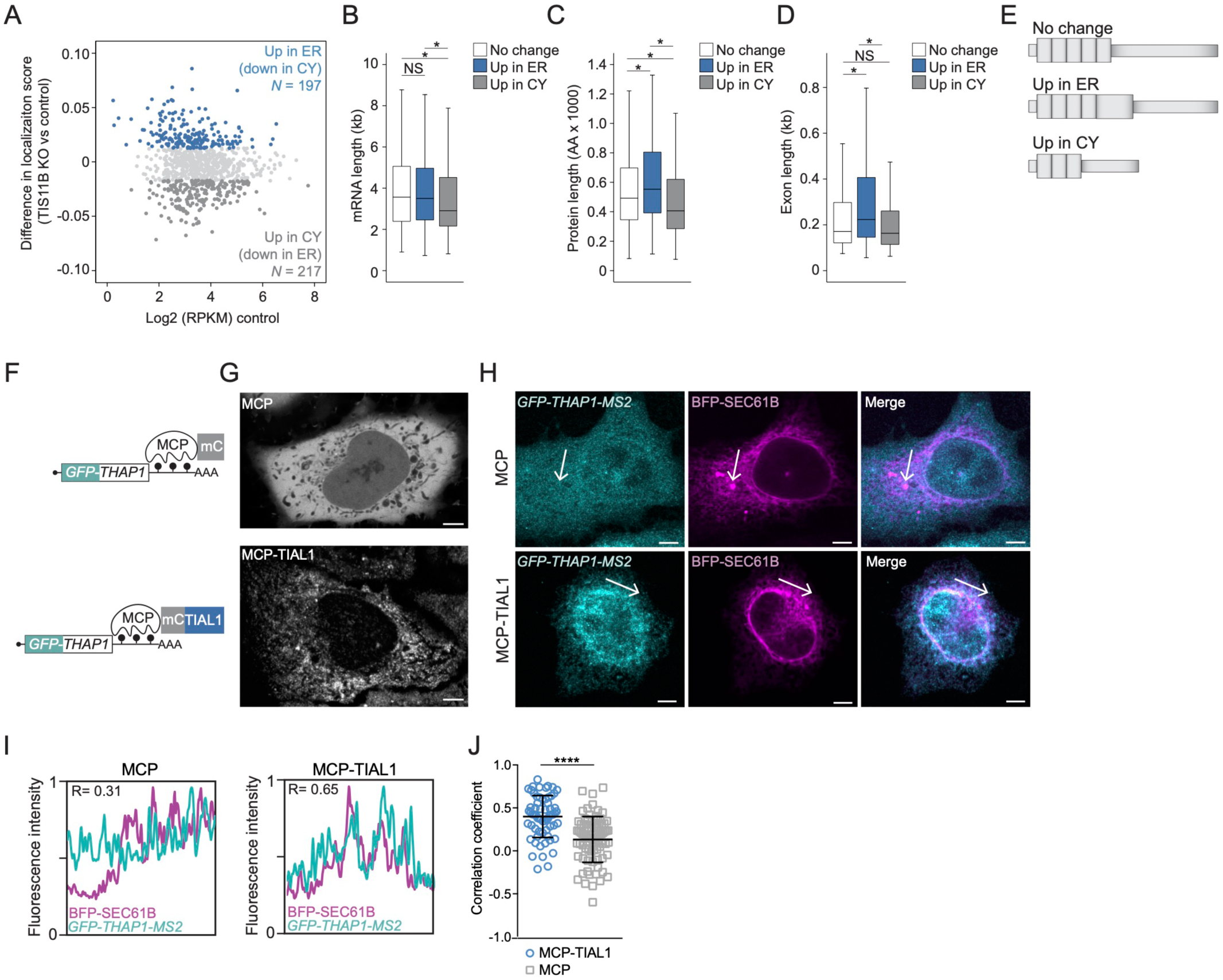
Experimental validation of the regulators of subcytoplasmic mRNA transcript distribution. 4A. TG+ mRNAs are shown and are color-coded based on their change in compartment localization in TIS11B KO cells compared with control cells. No change (*N* = 508); up in the ER (*N* = 197); up in CY (*N* = 217). 4B. Length distribution of mRNAs from (A). Mann Whitney test was performed. *P* value categories as in Fig. 2A. Exact *P* values are listed in Table S3. 4C. As in (B) but shown is protein size distribution. 4D. As in (B) but shown is CDS exon length distribution. 4E. As in Fig. 3E but shown are the mRNA features of TG+ mRNAs that change their localization upon TIS11B KO. 4F. Schematic of the mRNA reporter used to validate the effect of a single 3′UTR-bound RBP on mRNA localization. The *GFP-THAP1* reporter mRNA contains MS2 hairpins as 3′UTR, which allow binding of the cotransfected MS2 coat protein (mCherry-tagged MCP). Fusion of TIAL1 to MCP tethers it to the 3′UTR of the reporter mRNA. mC, mCherry. 4G. Confocal live cell imaging of HeLa cells expressing the indicated constructs. Scale bar, 5 µm. 4H. RNA-FISH (teal) of the GFP reporter mRNA from (F) in HeLa cells coexpressing the indicated MCP-fusion construct together with BFP-SEC61B to visualize the rough ER (magenta). Representative confocal images are shown. Scale bar, 5 µm. 4I. Line profiles of the fluorescence intensities obtained from the arrows shown in (H) together with the obtained Pearson’s correlation coefficients. 4J. Quantification of the Pearson’s correlation coefficients between the *GFP-THAP1* reporter mRNA and the rough ER in the experiment shown in (H). Two line profiles were generated for each cell. For MCP, *N* = 26 cells and for MCP-TIAL1 *N* = 21 were analyzed. The horizontal line denotes the median and the error bars denote the 25^th^ and 75^th^ percentiles. Mann-Whitney test, ****, *P* < 0.0001.

We did not find specific RBPs associated with the localization-changing mRNAs, because TG+ mRNAs are mostly bound by TIS11B and only a small minority of them (13% and 15%) are LARP4B or TIA1/L1 targets (Table S5). However, the TG+ mRNAs that increased their cytosolic localization upon TIS11B KO were the shortest, encoded the smallest proteins, and had the shortest exon length (Fig. 4B-E). In contrast, TG+ mRNAs that increased their ER localization upon deletion of TIS11B were significantly longer, encoded the largest proteins, and had longer exons (Fig. 4B-E).

These results support a model in which mRNA architecture features set up a ‘default’ steady-state mRNA transcript distribution pattern in the cytoplasm which can be overcome or reinforced through the binding of RBPs. Short mRNAs with average exon length localize to TGs when bound by TIS11B but in the absence of TIS11B they revert to the transcript distribution established by mRNA architecture and the remaining bound RBPs, in this case the cytosol (Fig. 3E). Similarly, longer TG+ mRNAs that encode the largest proteins localize to the ER upon loss of TIS11B (Fig. 3G). Currently, the ‘readers’ of the mRNA architecture features are unknown.

### 3′UTR-bound TIAL1 promotes localization of non-membrane protein-encoding mRNAs to the ER

We observed that ER+ mRNAs were enriched in 3′UTR-bound TIA1/L1, which had not been previously reported (Fig. 3A, 3C, 3F). To test the influence of TIA1/L1 on mRNA localization, we set out to investigate mRNA localization changes to the three compartments in TIA1/L1 double KO cells. ^36^ However, as previously reported, these cells showed a high rate of cell death, which prevented us from obtaining high-quality particles.

To validate TIA1/L1-dependent mRNA localization to the ER, we used the MS2 tethering system to mimic 3′UTR-binding of TIA1/L1 (Fig. 4F). We generated a *GFP-THAP1* reporter mRNA that contained MS2-binding sites as 3′UTR. ^37–39^ Coexpression of mCherry-tagged MS2 coat protein (MCP) fused to TIAL1 tethers TIAL1 to the 3′UTR of the reporter mRNA (Fig. 4F). As a control, mCherry-tagged MCP was tethered to the *GFP-THAP1* reporter mRNA.

Coexpression of the reporter mRNA and MCP resulted in evenly distributed cytosolic expression of both MCP protein and reporter mRNA (Fig. 4F-H). In contrast, coexpression of the reporter mRNA and MCP-TIAL1 resulted in perinuclear, reticulated expression of MCP-TIAL1 with the mRNA reporter predominantly localizing to the rough ER (Fig. 4F-H). Colocalization was assessed by RNA-FISH of the GFP-tagged reporter mRNA and simultaneous visualization of the rough ER through fluorescently tagged SEC61B. Using line diagrams of the fluorescence intensities, we quantified the overlap between the reporter mRNAs and the ER (Fig. 4I). In the presence of MCP-TIAL1, we observed higher correlation coefficients between the reporter mRNA and the ER (Fig. 4J). This result indicated that 3′UTR-bound TIAL1 was sufficient to induce localization of non-membrane protein encoding mRNAs to the rough ER surface.

### 3′UTR-bound TIAL1 increases protein expression

For endogenous mRNAs, we observed that ER+ mRNAs encode the highest expressed proteins (Fig. 2B). Moreover, mRNAs predominantly bound by TIA1/L1 encode proteins with higher expression levels compared with other mRNAs (Fig. 5A). Using our mRNA reporter (Fig. 4F), we investigated the contribution of TIAL1 to steady-state protein expression. We used FACS to measure GFP protein expression of the mRNA reporter with or without tethering of TIAL1 to the 3′UTR (Fig. S6A-C). We observed a 3.5-fold increase in protein expression upon 3′UTR-tethering of TIAL1 compared to tethering of MCP alone (Fig. 5B, 5C). Increased GFP protein expression was not due to an increase in mRNA abundance (Fig. 5D). We confirmed the TIA1/L1-dependent increase in protein expression using a second GFP reporter (Fig. S6D-F). As TIAL1 promotes translation of mRNAs on the ER membrane, it was unclear if increased protein expression was caused by TIAL1 or by a potentially unique translation environment provided by the rough ER membrane. For example, it was reported that mRNAs that encode non-membrane proteins contain 1.4-fold more ribosomes when translated on the ER membrane than when translated in the cytosol. ^40^

**Figure 5.**
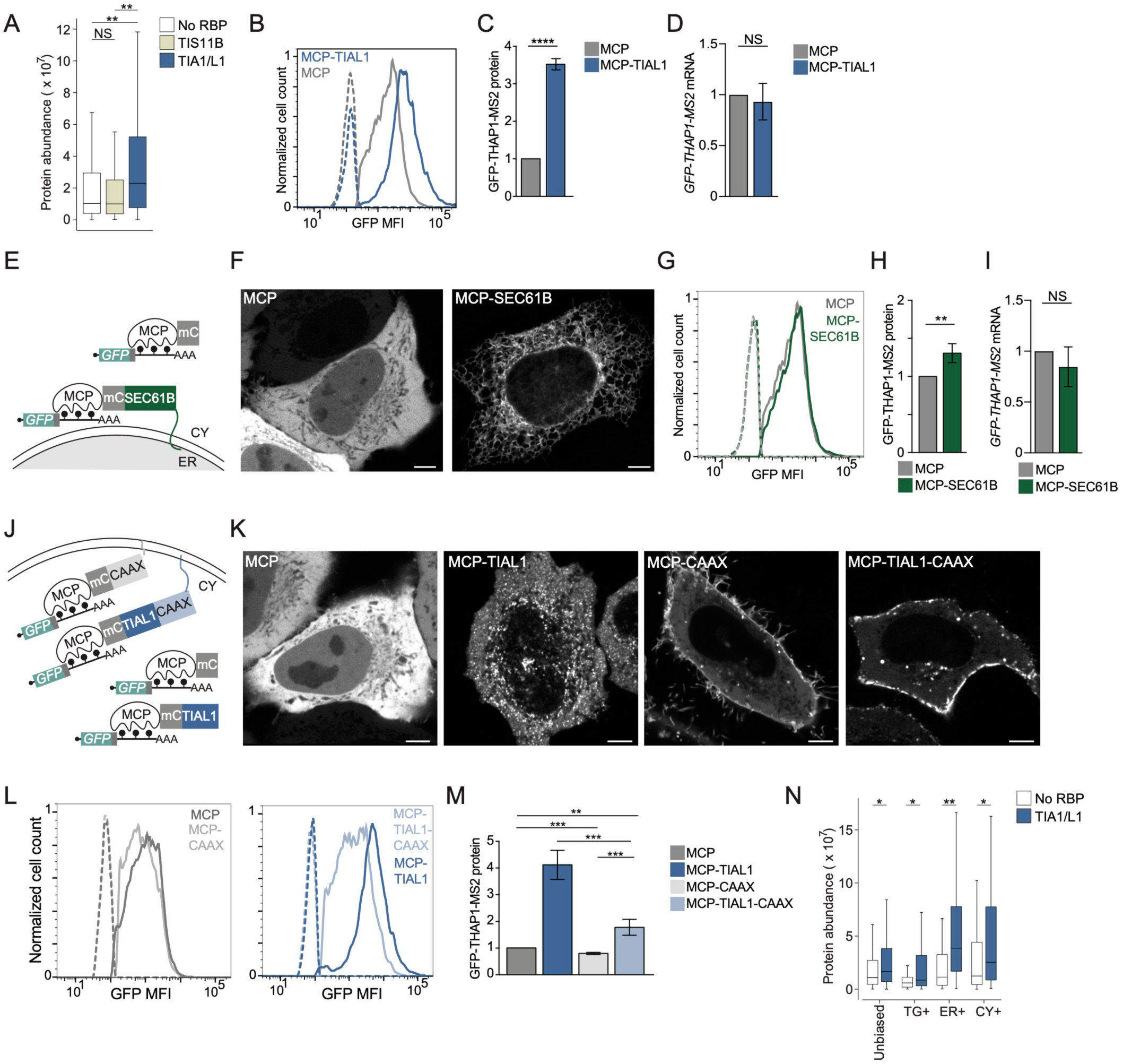
3′UTR-bound TIAL1 cooperates with the rough ER membrane environment to increase protein expression. 5A. Protein abundance of mRNAs stratified by RBP binding. No RBP (*N* = 126), bound only by TIS11B (*N* = 267), bound only by TIA1/L1 (*N* = 232). Mann Whitney test was performed. *P* value categories as in Fig. 2A. 5B. GFP protein expression measured by FACS in HeLa cells using the *GFP-THAP1* reporter mRNA with and without TIAL1 tethering. Representative histograms are shown. GFP-negative cell populations are shown as dotted lines. 5C. Quantification of the experiment shown in (B). Shown is the mean ± std of five independent experiments. T-test for independent samples, ****, *P* = 0.0003. 5D. Quantification of mRNA level of the experiment from (B). Shown is the mean ± std of three independent experiments. T-test for independent samples, NS, not significant. 5E. Schematic of a *GFP-THAP1* mRNA reporter that investigates the influence of subcellular mRNA localization on protein expression. Fusion of MCP to SEC61B localizes the GFP reporter mRNA (shown as in Fig. 4F) to the rough ER membrane, whereas MCP alone localizes it to the cytosol. 5F. Confocal live cell imaging of HeLa cells expressing the indicated constructs. Scale bar, 5 µm. 5G. As in (B) but the *GFP-THAP1* reporter mRNA was used with and without SEC61B tethering. 5H. Quantification of the experiment from (G). Shown is the mean ± std of four independent experiments. T-test for independent samples, **, *P* =0.0026. 5I. Quantification of mRNA level in the experiment from (G). Shown is the mean ± std of three independent experiments. T-test for independent samples, NS, not significant. 5J. As in Fig. 4F, but addition of a prenylation signal (CAAX) localizes the TIAL1-bound *GFP-THAP1* reporter mRNA to the plasma membrane. In the absence of CAAX, the TIAL1-bound reporter mRNA localizes to the rough ER. 5K. Confocal live cell imaging of HeLa cells expressing the indicated constructs. Scale bar, 5 µm. 5L. As in (B) but the *GFP-THAP1* reporter mRNA was tethered with the indicated constructs. 5M. Quantification of the experiment from (L). Shown is the mean ± std of four independent experiments. T-test for independent samples, ****, *P* < 0.0006, **, *P* = 0.002. 5N. Endogenous mRNAs bound by TIA1/L1 encode higher expressed proteins than mRNAs not bound by any RBP. The largest TIA1/L1-associated increase was observed for ER+ mRNAs. Mann Whitney test was performed. *P* value categories as in Fig. 2A.

### TIAL1 cooperates with the rough ER environment to promote protein expression

To disentangle the effects of TIAL1 and the ER membrane on protein expression, we tethered the reporter mRNA directly to the ER surface by fusing MCP to SEC61B, a subunit of the translocon complex in the rough ER (Fig. 5E). MCP-SEC61B perfectly colocalized with the ER and recruited reporter mRNAs to the ER (Fig. 5F, S6G-I). However, reporter protein expression only increased by 1.25-fold compared to the tethering of MCP alone and did not increase mRNA abundance of the reporter (Fig. 5G-I). We used a second ER localization reporter by fusing MCP to TRAPα, which represents a different subunit of the translocon complex and obtained a similar result. We observed an increase in protein expression by 1.5-fold when the reporter mRNA was tethered to TRAPα (Fig. S6J-M). These results suggested that the ER membrane environment has a significant but small stimulatory effect on translation.

Next, we investigated if the TIAL1-dependent increase in protein expression is intrinsic to TIAL1 or if it depends on its localization to the ER membrane. We added a CAAX motif to TIAL1 to localize the TIAL1-bound mRNA reporter to the plasma membrane instead of the ER membrane (Fig. 5J). The CAAX signal is a prenylation motif that efficiently localized MCP and MCP-TIAL1 to the plasma membrane (Fig. 5K). ^32^ Translation of the TIAL1-bound mRNA reporter at the plasma membrane increased protein expression by 1.8-fold (Fig. 5L, 5M). As translation of the TIAL1-bound reporter at the ER membrane resulted in two-fold higher protein expression than its translation at the plasma membrane (Fig. 5M), our result suggested that TIAL1 cooperated with the environment on the rough ER membrane to promote protein expression.

As the RBPs bound to the reporter mRNA were identical in these experiments, our results demonstrate that the subcytoplasmic location of translation controls steady-state protein expression levels by two-fold when comparing plasma and ER membranes. This relationship was also observed for endogenous mRNAs, where TIA/L1 bound mRNAs were associated with high protein output in every compartment, but with the highest protein yields being observed in the ER compartment (Fig. 5N).

### The repressive effect of cytosolic TIS11B on protein expression is overcome by its localization to rough ER membrane

Next, we examined if the environment on the rough ER membrane also promotes protein expression of mRNAs bound by other RBPs, including TIS11B (Fig. 6A, 6B). In cells expressing mCherry-TIS11B fusion constructs, about 30% form TGs at steady state (Fig. S7A, S7B). ^16^ However, we noticed that addition of MCP to TIS11B fusion constructs resulted in limited TG formation and predominant expression of TIS11B in the cytosol (Fig. S7A, S7B). In the cytosolic state, binding of MCP-TIS11B to the reporter mRNA repressed reporter protein expression by two-fold, compared to tethering of MCP alone (Fig. 6C, 6D). This decrease in protein expression was partially caused by a TIS11B-dependent decrease in mRNA level (Fig. 6E), consistent with previous reports that suggested that cytosolic TIS11B represses the expression of certain cytokine and cell cycle mRNAs. ^41–43^ In contrast, fusing TIS11B to MCP-SEC61B localizes TIS11B and the bound reporter mRNA to the rough ER (Fig. 6A, 6B), which overcomes the repressive effect of cytosolic TIS11B and increased protein expression two-fold (Fig. 6A-E). The two-fold increase in protein expression was recapitulated with a second reporter and indicates that the repressive effect on protein expression mediated by cytosolic TIS11B is overcome by translation of the TIS11B-bound mRNA on the ER (Fig. 6D, Fig. S7C-E).

**Figure 6.**
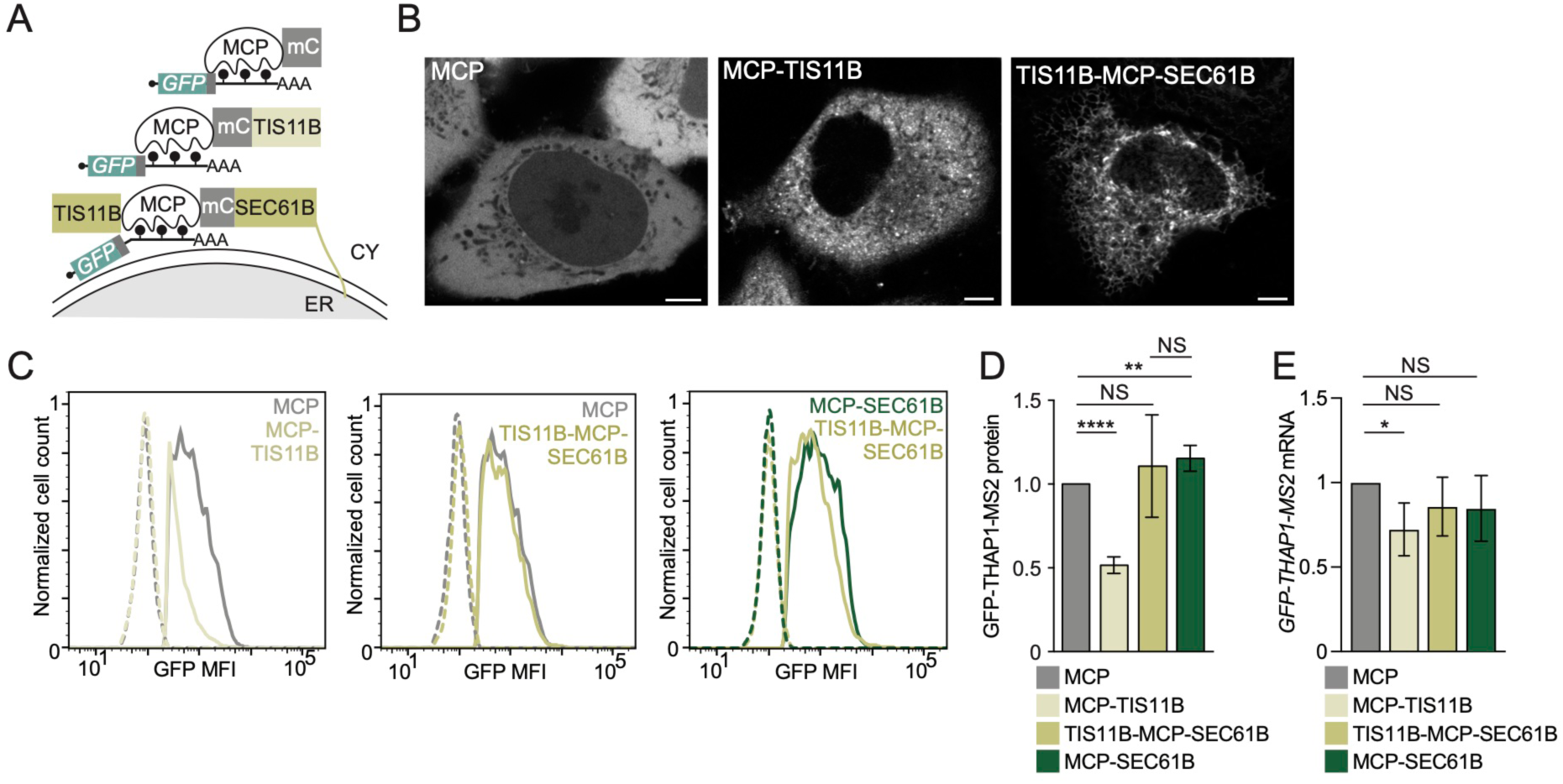
Localization of cytosolic mRNAs to the rough ER membrane increases their protein expression. 6A. Schematic of a *GFP-THAP1* reporter mRNA bound by TIS11B that allows investigation of localization-dependent GFP protein expression. Fusion of MCP and TIS11B localizes the mRNA reporter to the cytosol, whereas the TIS11B-MCP-SEC61B fusion localizes the mRNA to the rough ER membrane. 6B. Confocal live cell imaging of HeLa cells expressing the constructs from (A). Scale bar, 5 µm. 6C. As in Fig. 5B, but the *GFP-THAP1* reporter mRNA was tethered with the indicated MCP-fusion constructs. 6D. Quantification of the experiment from (C). Shown is the mean ± std of four independent experiments. T-test for independent samples, ****, *P* < 0.0001, **, *P* = 0.003, NS, not significant. 6E. Quantification of mRNA level in the experiment from (C). Shown is the mean ± std of three independent experiments. T-test for independent samples, *, *P* = 0.037; NS, not significant.

### Model

Taken together, we observed that mRNAs that are uniquely enriched in one of three cytoplasmic compartments differ substantially in their architectural features, in the RBPs bound to them, and in the expression levels and functional classes of their encoded proteins (Fig. 7). TG+ mRNAs are characterized by the longest CDS exons and TIS11B binding to the 3′UTR. These mRNAs encode the lowest abundance proteins with a strong enrichment of transcription factors. In contrast, although TGs are intertwined with the rough ER, ER+ mRNAs are the longest, are predominantly bound by TIA1/L1, and encode highly abundant large proteins. CY+ mRNAs are the shortest and encode small and highly abundant proteins. They are bound by LARP4B/METAP2 and have high production and degradation rates (Fig. 7). Moreover, by using mRNA reporters, we showed that relocation of protein synthesis from the cytosol to the ER increases protein expression, indicating that the location of translation influences protein output (Fig. 7).

**Figure 7.**
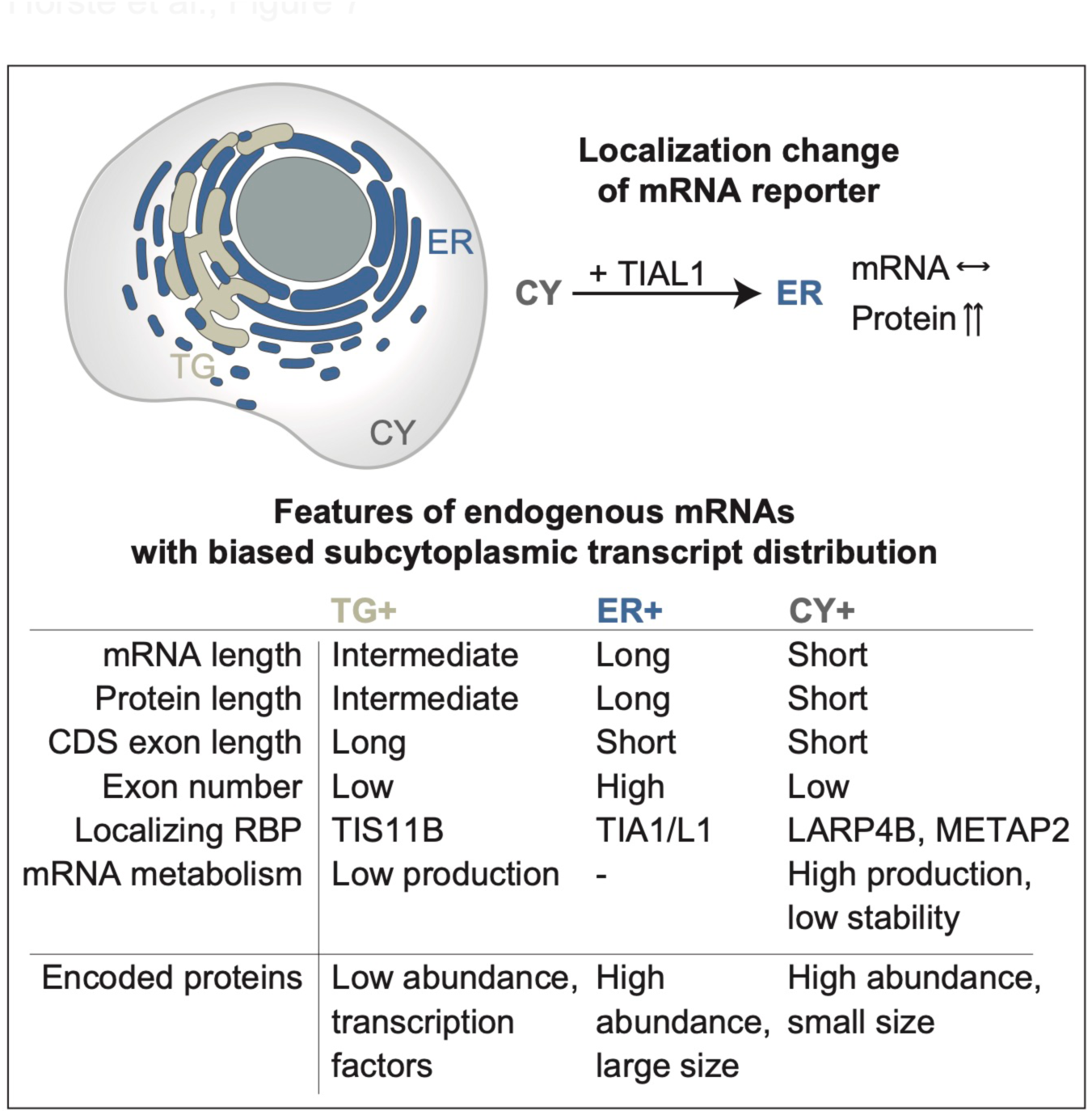
Model. Model showing features of endogenous mRNAs with biased subcytoplasmic transcript distribution. Summarized are also the changes in protein abundance upon relocalization of the reporter from the cytosol to the ER (see text for details). The horizontal arrow indicates no change.

## Discussion

We determined the distribution of endogenous mRNA transcripts across three cytoplasmic compartments, including TGs, the rough ER, and the cytosol under steady-state conditions. Our RNA-seq results, which were validated by smRNA-FISH, suggest that approximately half of all mRNAs are uniquely enriched in one of these three cytoplasmic compartments.

### Functionally related genes are translated in unique compartments

One of our most striking findings was that within each investigated compartment a different group of functionally related mRNAs is translated (Fig. 2). Moreover, the compartment-enriched mRNAs have vastly different gene architectures and are characterized by substantially different production and degradation rates as well as the expression levels of their encoded proteins (Fig. 2). These features are consistent with the compartment-enriched gene groups, indicating that the cytoplasm is strongly partitioned into different functional and regulatory compartments that are not enclosed by membranes.

Surprisingly, we observed that transcription factors are substantially enriched among TG+ mRNAs (Fig. 2J). This unexpected result can be explained by the previous observation that transcription factors are often present in low abundance ^31^ and we found that TG+ mRNAs encode the proteins with the lowest expression levels (Fig. 2B). Moreover, many transcription factors have an unusual gene architecture with longer than average coding exons. Together with TIS11B binding, this was the strongest feature for mRNA enrichment in TGs (Fig. 3C-E). Interestingly, both characteristics correlate with features associated with low mRNA abundance levels, but whereas TIS11B-binding correlates negatively with pre-mRNA production rates (Spearman’s correlation coefficient R = −0.26), exon length negatively correlates with mRNA half-life (Spearman’s correlation coefficient R = −0.34). ^44^ The unique gene architecture together with predominant binding of TIS11B provides an explanation for why TGs enrich for low-abundance mRNAs.

In contrast, ER+ mRNAs encode the largest proteins with the highest expression levels. These include helicases, cytoskeleton-bound proteins, and chromatin regulators (Fig. 2). It is possible that anchoring of ribosomes on the ER membrane may facilitate the protein synthesis of very large proteins. Moreover, it is notable that despite the intertwinement of TGs and the rough ER, the compartment-enriched mRNAs encode proteins that differ substantially in their expression levels, which are the lowest for TG+ mRNAs and the highest for ER+ mRNAs.

It was previously shown that localization to the ER membrane of certain non-membrane protein encoding mRNAs increases their translation, ^9,40^ and we confirmed this result. In addition, we describe a new role for TIAL1 in the regulation of translation, as TIAL1 binding substantially increased mRNA translation (Fig. 5C). So far, TIA1 and TIAL1 have mostly been described as regulators of pre-mRNA splicing and as translational repressors in the context of cellular stress, where they assemble into stress granules. ^45,46^ However, in the absence of stress, TIA1/L1 has been reported to promote polysome association which supports our findings. ^36,47^ For both reporter mRNAs and endogenous mRNAs, we observed that the presence of TIAL1 increased protein expression in all compartments, but only in the context of the ER did we observe a cooperative effect on translation (Fig. 5M, 5N). The factor that cooperates with TIAL1 on the ER to upregulate translation is currently unknown. Importantly, our reporter results demonstrate that a change in the location of protein synthesis within the cytoplasm strongly influences protein output, indicating that a change in mRNA localization can alter protein abundance.

### A combinatorial code of mRNA architecture features and 3′UTR-bound RBPs controls subcytoplasmic mRNA transcript distribution

RBPs play an established role in mRNA localization. ^1,7^ Additionally, we observed a strong association of mRNA architecture features with mRNA transcript localization to the three cytoplasmic compartments (Fig. 3C). It is most likely that mRNA length, CDS length, and CDS exon length are not direct regulators of mRNA localization but that specific factors, including currently unknown RBPs, read-out this information. In addition, we speculate that mRNA architecture influences mRNP size, conformation, and packaging ^48,49^ and that these biophysical features may act as additional determinants of subcytoplasmic mRNA localization. This idea is supported by previous insights into *oskar* mRNA localization, where the deposition of the exon junction complex, which is involved in mRNP packaging, ^48,49^ was found to be required for proper mRNA localization in the cytoplasm. ^50^

We present a model for the regulation of subcytoplasmic transcript distribution that is based on a combinatorial code generated by mRNA architecture features together with the bound RBPs, where individual components act in an additive manner (Fig. 3E, 3G). This model was tested experimentally by analyzing the localization propensity of TG+ mRNAs upon deletion of TIS11B. The obtained results confirmed the contribution of mRNA architectural features to mRNA localization and suggest that the binding of RBPs can overcome the default localization pattern established by the mRNA architecture features (Fig. 4A-E).

### Is it biologically relevant if only 20% of transcripts localize to TGs?

Based on the estimated size of TGs (Fig. 1I), we expect that 11% of mRNA transcripts localize to TGs by chance. Using smRNA-FISH on three individual TG+ mRNAs we observed a two-fold enrichment in TGs, meaning that on average 22% of these transcripts localize to TGs. This raises the important question: whether it matters biologically that a minority population of transcripts for a given mRNA localizes to a certain compartment.

This question was addressed in a follow-up project, where we investigated the biological consequences of *MYC* mRNA, which is a TG+ mRNA, when it was translated in TGs or the cytosol. ^19^ We observed that several MYC protein complexes were only formed when *MYC* mRNA was translated in TGs and not when it was translated in the cytosol. The TG-dependent protein complexes formed co-translationally and had functional consequences for MYC target gene expression in the nucleus. TG-translated MYC induced different target genes than cytosol-translated MYC. ^19^ Our results indicate biological relevance, even when only a fraction of transcripts are translated in TGs.

In summary, our study revealed a surprisingly high degree of cytoplasmic compartmentalization. This is the basis for the translation of functionally related proteins in defined environments that strongly affect mRNA and protein expression. Our results highlight the contribution of spatial regulation whose consequences go beyond the effects mediated by the mRNA-bound proteins. In the future, our findings may provide the basis for biotechnology applications that make use of engineered 3′UTR sequences to boost protein expression in experimental settings or to increase protein production of mRNA vaccines.

### Limitations of our study

The exact compartment sizes of TGs, the rough ER, and the cytosol are currently unknown and can only be estimated. However, using two different methods for the identification of compartment-enriched mRNAs yielded highly similar enrichment values.

To obtain sufficient material for TG and ER particle sorting, we used transfected, fluorescently labeled proteins instead of endogenous proteins. In cells that highly express TIS11B, in the future, TG particle sorting may be possible using endogenous, fluorescently tagged TIS11B.

The use of spike-ins to isolated compartments obtained from defined cell numbers may have enabled us to perform absolute enrichments versus the relative enrichment analyses that we report here. Moreover, all analyses were performed at the gene level. Alternative 3′UTR isoforms are known to differentially localize and therefore, we would expect to obtain a higher resolution for compartment enrichment of transcripts if instead of genes alternative 3′UTR isoforms are analyzed. ^38,51^ However, with our purification strategy we did not obtain sufficient mRNA quantities to perform the analysis at the level of alternative 3′UTR isoforms.

## Acknowledgements

This work was funded by the NIH Director′s Pioneer Award (DP1-GM123454), the R35GM144046 NIH grant, a grant from the Pershing Square Sohn Cancer Research Alliance, a grant from the G. Harold and Leila Y. Mathers Foundation, and the MSK Core Grant (P30 CA008748) to CM. J.U. received funding from the ERC under the European Union Horizon 2020 Research and Innovation Program (835300-RNPdynamics) and The Francis Crick Institute receives its core funding from Cancer Research UK (CC0102), the UK Medical Research Council (CC0102), and the Wellcome Trust (CC0102). An NIH F31 (CA254335-01) fellowship was awarded to E.L.H.

We thank Yevgeniy Romin and Eric Chan from the Molecular Cytology Core Facility for help with microscopy and image analysis. We thank Kathleen Daniels from the Flow Cytometry Core Facility (now Sana Biotechnologies) for help with particle sorting and Liana Boraas at Yale University School of Medicine for help with endogenous RNA-FISH. We thank Weirui Ma (Zhejiang University) for experimental guidance and mentorship of E.L.M. and Alban Ordureau for analysis of the TMT mass spectrometry data. We thank Yang Luo (MSKCC) and Ben Kleaveland (WCMC) for helpful discussions and critical reading of the manuscript.

## Author contributions

E.L.H. performed all experiments, except the mass spectrometry analysis whose samples were prepared by X.C. and the TIS11B KO cells were generated by S.M. F.C.Y.L. and J.U. performed and analyzed the TIS11B iCLIP experiment. M.M.F performed the logistic regression and provided the gene architecture features. T.C. analyzed the RNA-seq data. G.Z. analyzed the CLIP data with input from C.M. E.L.H. and C.M. conceived the project, designed the experiments, and wrote the manuscript with input from all authors.

## Methods

### STAR methods table

#### Cell lines

##### Generation of a doxycycline inducible *TIS11B* knockout cell line (*TIS11B* KO)

Doxycycline inducible Cas9 (iCas9) HEK293T cells were generated by infecting cells with lentivirus containing a Cas9-P2A-GFP expression cassette under a doxycycline inducible promoter as described previously. ^52^ During consecutive rounds of fluorescence-activated cell sorting, we selected a cell pool exhibiting robust induction of Cas9/GFP expression after doxycycline treatment (100 ng/ml for 24 hours), and low levels of leaky transgene expression in the absence of the drug.

Next, we transduced iCas9 cell lines with a lentiviral construct harboring a pair of guide RNAs either targeting *TIS11B* or gRNAs that target an intergenic region. To generate these constructs, we adapted the plentiGuide-puro vector. ^53^ to incorporate a second guide RNA expression cassette as described previously. ^54^ For this purpose, the plasmid was digested with BsmBI (FastDigest Esp3I, Thermo Fisher Scientific) and a synthetic 391 bp double-stranded DNA fragment encoding 5′-(1st gRNA/scaffold/H1 promoter/2nd gRNA)-3′ was inserted using the NEBuilder HiFi assembly system (NEB). Synthetic DNA fragments were ordered from Genewiz and sequences are listed in Table S6. The assembled vector DNA was used to transform chemically competent Stbl3 bacteria cells (Invitrogen), and correct vector clones were identified by Sanger sequencing.

Lentivirus was generated in HEK293T cells using standard methods and 200 μl of viral supernatant was used to transduce iCas9 cells in a 6-well dish together with 8 μg/ml polybrene. Transduced cells were subjected to puromycin selection (1 μg/ml) for five days and resistant cells were aliquoted and frozen for all further experiments. Finally, for induction of gene knockouts, *TIS11B* KO and corresponding control cells (with gRNAs targeting an intergenic region) were treated with doxycycline (100 ng/ml) for five days, after which TIS11B protein expression was evaluated by western blotting, ER particle sorting and digitonin extraction was performed.

### Detailed methods

#### Constructs

##### Fluorescently-tagged TIS11B and SEC61B constructs

The eGFP/mCherry/BFP fusion constructs for TIS11B and SEC61B expression were described previously. ^16^ They were generated in the pcDNA3.1-puro expression vector. The TIS11B and SEC61B coding regions were PCR amplified from HeLa cDNA and inserted downstream of eGFP/mCherry/BFP using BsrGI/EcoRI or BsrGI/HindIII restriction sites, respectively.

##### Constructs to generate the mRNA localization reporter

To investigate the influence of RBPs on mRNA localization of a GFP mRNA reporter, RBPs were fused to MCP and tethered to a GFP mRNA reporter containing MS2 binding sites as 3′UTR. ^37,38^ To investigate mRNA localization-dependent protein expression of the GFP mRNA reporter, a CAAX sequence was fused to MCP or to MCP-RBP fusions.

##### GFP mRNA reporter

To generate the GFP mRNA reporter, the GFP-BIRC3-MS2-SU ^39^ vector was used the BIRC3 coding region was replaced with the THAP1 coding region. It was PCR amplified from the GFP-THAP1 vector using THAP1-MS2 F and THAP1-MS2 R primers and inserted between the BsrGI and AgeI sites. The SU fragment was removed with HindIII and XhoI and blunt end ligated, resulting in GFP-THAP1-MS2.

##### MCP-mCherry RBP fusion constructs

To generate MCP-mCherry, the MCP coding sequence was PCR amplified from UBC NLS-HA-2XMCP-tagRFPt vector (Addgene 64541) using MCP F and MCP R primers and inserted in-frame, upstream of mCherry (mCherry lacking a start codon) between BmtI and BamHI sites in pcDNA3.1-puro-mCherry vector. ^16^ To generate MCP-mCherry-TIS11B and MCP-mCherry-TIAL1, their coding sequences were inserted in-frame, downstream of mCherry between the BsrGI and XbaI sites. The TIS11B coding sequence was amplified from pcDNA3.1-puro-GFP-TIS11B using TIS11B MCP F and TIS11B MCP R primers and the TIAL1 coding sequence was PCR amplified from pFRT_TO_FlagHA_TIAL1 (Addgene 106090) using TIAL1 MCP F and TIAL1 MCP R primers.

##### MCP-mCherry fusion constructs with subcellular localization signals

To generate pcDNA3.1-puro-MCP-mCherry-SEC61B, the MCP-mCherry coding sequence was cut from MCP-mCherry vector using BmtI and BsrGI and pasted in-frame, upstream of SEC61B in pcDNA3.1-mCherry-SEC61B (replacing mCherry). To generate the TIS11B-MCP-mCherry-SEC61B vector, TIS11B coding sequence was PCR amplified from pcDNA3.1-puro-GFP-TIS11B using TIS-SEC F and TIS-SEC R primers and pasted in-frame, upstream of MCP into the BmtI site in the MCP-mCherry-SEC61B vector. To generate TRAPα-MCP-mCherry, the TRAPα coding sequence (encoded by the *SSR1* gene) was PCR amplified from HeLa cDNA using TRAPα MCP F and TRAPa MCP R and inserted in-frame, upstream of MCP in the pcDNA3.1-puro-MCP-mCherry vector.

For plasma membrane localization, the CAAX prenylation signal was added to the C-terminus of MCP-mCherry or MCP-mCherry-TIAL1. The CAAX coding sequence was purchased as a gene fragment from Azenta as described ^32^ and PCR amplified using TIAL1 CAAX F and CAAX R primers. It was inserted in-frame using the BsrGI and ApaI sites, located downstream of mCherry to generate pcDNA3.1-puro-MCP-mCherry-CAAX. It was inserted in-frame using EcoNI and ApaI sites to generate MCP-mCherry-TIAL1-CAAX.

##### SunTag constructs

were described previously. ^32^

#### Isolation of subcytoplasmic compartments

##### Transfection

HEK293T cells were seeded in six 10 cm dishes (particle sorting) or one well from a 6-well plate (cytosol extraction) at 80% confluency in antibiotic free media. After 24 hours, cells were transfected by calcium phosphate with either 3 µg mCherry-TIS11B or 1 µg GFP-SEC61B per dish (particle sorting), or 500ng mCherry-TIS11B (cytosol extraction).

##### Particle purification

20 hours after transfection, cells were rinsed once with ice-cold PBS, scraped in 10 ml ice-cold PBS, and pelleted at 300 x g. Pellets from two plates were resuspended in 1 ml ice-cold hypotonic isolation buffer (225 mM mannitol, 75 mM sucrose, 20 mM Tris-HCl pH 7.4, 0.1 mM EDTA). Cells were lysed with 50 strokes in a 1 ml dounce-homogenizer with pestle on ice in order to shear the nuclei from the ER. Nuclei were pelleted with a two-minute spin at 600 x g. The supernatant contains the cytoplasmic membrane fraction, which was pelleted with a 15-minute spin at 7000 x g and resuspended in ice-cold PBS for fluorescent particle sorting.

##### Fluorescent particle sorting

Particles were sorted on a BD FACSAria III cell sorter equipped with a 70 µm nozzle. The forward-scatter threshold was decreased from 5,000 to 800 in order to visualize subcellular particles. Particles were first detected by fluorescence using the 594 nm and 488 nm excitation lasers, for mCherry-TIS11B and GFP-SEC61B respectively, and 405 nm excitation laser for DAPI. A sorting gate was drawn on particles that were either mCherry-positive or GFP-positive, but DAPI-negative, to exclude any remaining nuclei. Sorting was performed in purity mode with an average speed of 150 particles/second. Particles were sorted directly into 1 ml of TRIzol solution in Eppendorf tubes, holding 180,000 particles per tube. RNA extraction was performed for each tube separately and total RNA for each sample was combined for library preparation. Two biological replicates for each particle prep were sequenced. For each replicate, about 1.5 million TIS11B granule particles and 2.0 million ER particles were collected.

##### Cytosol extraction

The cytosol was extracted as previously described. ^24^ HEK293T cells transfected were plated in a six-well plate at 80% confluency. After 24 hours, cells were rinsed once in the dish with ice-cold PBS. After aspirating PBS, 300 µl ice-cold digitonin solution (40 µg/ml digitonin, 150 mM NaCl, 20 mM HEPES pH 7.4, 0.2 mM EDTA, 2 mM DTT, 2 mM MgCl2) was added and incubated on a shaker at 4°C for ten minutes. After incubation, the digitonin-derived cytosolic extract was pipetted from the plate and spun at 20,000 x g for one minute to pellet any floating cells. 200 µl of cytosolic extract was added to 1 ml TRIzol solution for RNA extraction.

#### RNA-seq library preparation

RiboGreen RNA Reagent (ThermoFisher) was used for RNA quantification and quality control was performed by Agilent BioAnalyzer. 50-500 ng of total RNA underwent polyA selection and TruSeq library preparation according to instructions provided by Illumina (TruSeq Stranded mRNA LT Kit, catalog # RS-122-2102), with eight cycles of PCR. Samples were barcoded and run on a HiSeq 4000 in a PE50 run, using the HiSeq 3000/4000 SBS Kit (Illumina). An average of 27 million paired reads was generated per sample.

#### Western Blotting

For whole cell lysate preparation, cells were trypsinized and washed twice with PBS and lysed in 2x Laemmli Sample buffer (Alfa Aesar, J61337). For cytosolic lysate, cytosol was extracted with digitonin as described above and one volume of 2x Laemmli Sample buffer was added. Laemmli lysates were boiled for 10 min at 95°C. Samples were subjected to SDS-PAGE on NuPAGE 4%–12% Bis-Tris gradient protein gel (Invitrogen). Imaging was captured on the Odyssey DLx imaging system (Li-Cor). Quantification was performed using ImageJ. The antibodies used are listed in the Key Resources Table.

#### TIS11B iCLIP

##### Transfection

HEK293T cells were seeded in 10 cm dishes at 80% confluency in antibiotic free media. After 24 hours, cells were transfected by calcium phosphate with either 3 µg GFP-TIS11B or 1.5 µg GFP-only per dish.

##### Sample preparation

20 hours after transfection, cells were rinsed once with ice-cold PBS and 6 ml of fresh PBS was added to each plate before crosslinking. Cells were irradiated once with 150 mJ/cm^2^ in a Spectroline UV Crosslinker at 254 nm. Irradiated cells were scraped into Eppendorf tubes, spun at 500 x g for one minute, and snap-frozen. Crosslinked cell pellets were lysed in iCLIP lysis buffer (50 mM Tris-HCl pH 7.4, 100 mM NaCl, 1% Igepal CA-630 (Sigma I8896), 0.1% SDS, 0.5% sodium deoxycholate), sonicated with the Bioruptor Pico for 10 cycles 30 seconds ON/30 seconds OFF, and supplemented with 0.5 U of RNase I per 1 mg/ml lysate for RNA fragmentation. Lysates were pre-cleared by centrifugation at 20,000 x g at 4°C. A mix of Protein A/G Dynabeads (50 µl of each per sample, Life Technologies) were coupled to 10 µg of rabbit anti-GFP antibody (Abcam ab290). TIS11B protein-RNA complexes were immunoprecipitated from 1 ml of crosslinked lysate and washed with high salt and PNK buffer (NEB). RNA was repaired by 3′ dephosphorylation and ligated to L3-IR adaptor on beads. ^55^ Excess adaptor was removed by incubation with 5′ deadenylase and the exonuclease RecJf (NEB). TIS11B protein-RNA complexes were eluted from the beads by heating at 70°C for one minute. The complexes were then visualized via the infrared-labeled adaptor, purified with SDS-PAGE, and transferred to nitrocellulose membrane. cDNA was synthesized with Superscript IV Reverse Transcriptase (Life Technologies) and circularized by CircLigase II. Circularized cDNA was purified with AmPURE bead-based purification (A63880, Beckman Coulter), amplified by PCR and sequenced by Novaseq.

#### RNA-FISH

##### Single molecule RNA-FISH for endogenous mRNAs. Probe design

Primary probes were designed using the ProbeDealer package in MATLAB. ^56^ Each primary probe contains 30 transcript-targeting nucleotides preceded by 20 common nucleotides that are complementary to the secondary probe. At least 30 probes were designed for each transcript, purchased in a pool from IDT. The secondary probes are 5′ conjugated to AlexaFluor 633 and were purchased from IDT.

##### Transfection

Prior to cell seeding, 35 mm glass cover slips were sterilized with ethanol then incubated in 1 µg/ml fibronectin in PBS at room temperature for one hour. Cover slips were rinsed in PBS and HeLa cells were seeded at 100,000 per coverslip. 24 hours after seeding, cells were co-transfected with 250 ng BFP-TIS11B and 100ng of GFP-SEC61B using Lipofectamine 3000 (Invitrogen).

##### Sample preparation

20 hours after transfection, cells were rinsed once with PBS then fixed in 4% paraformaldehyde for 10 minutes at room temperature. All steps were performed at room temperature if not otherwise noted. Cells were rinsed twice with PBS and permeabilized with 0.5% Triton-X solution for 10 minutes. Cells were rinsed twice with PBS and incubated for five minutes in pre-hybridization buffer (2xSSC, 50% formamide). Cells were incubated in primary probe hybridization solution (40 µM primary probe, 2xSSC, 50% formamide, 10% dextran sulfate (Sigma), 200 µg/ml yeast tRNA (Sigma), 1:100 Murine RNase Inhibitor (NEB)), for at least 15 hours at 37°C. To remove excess or unbound primary probes, cells were then rinsed twice in 2xSSC + 0.1% Tween for 15 minutes at 60°C then once more for 15 minutes at room temperature. Cells were incubated in secondary probe solution (4 nM secondary probe, 2xSSC, 50% ethylene carbonate, 1:100 Murine RNase Inhibitor) for 30 minutes in the dark. Secondary probes were rinsed twice in 50% ethylene carbonate, 2xSSC solution for five minutes then mounted with Prolong Diamond mounting solution (Invitrogen).

##### Cytosol extraction

To visualize and validate CY+ versus TG+ or ER+ endogenous mRNAs, HeLa cells were seeded as described above, then incubated in 2 ml digitonin solution described above (40 µg/ml digitonin, 150 mM NaCl, 20 mM HEPES pH 7.4, 0.2 mM EDTA, 2 mM DTT, 2 mM MgCl2) for 10 min at 4°C. Digitonin solution was removed, coverslips were rinsed with 2 ml PBS, and RNA-FISH was performed as described above. Mounting media with DAPI was used to visualize nuclei (Invitrogen P36931).

##### Validation of TG+ and ER+ mRNAs using smRNA-FISH

We performed smRNA-FISH on endogenous mRNAs (Table S2) while simultaneously visualizing TGs and the ER. We considered an mRNA to have an unbiased localization pattern if its transcript distribution correlated with the cytoplasmic compartment sizes. As a proxy for the relative compartment sizes, we used the area occupied by TGs or the ER compared to the whole cell area, obtained from the maximum projection of the fluorescent signals in 186 cells. We used FIJI to delineate the whole cell border with the fluorescent signal from RNA-FISH. For TGs, the fluorescent signal from BFP-TIS11B and for the ER the fluorescent signal from GFP-SEC61B both obtained from the maximum intensity Z-projections was used to delineate each compartment. Where there was overlap between the TG mask and the ER mask, the ER was subtracted, and the region was defined as TG. In this way the compartments are mutually exclusive. The mask area of each compartment was quantified and read out as a proportion of the total cell area. Across all cells, the median size of TGs was estimated to be 11% of the cell size, whereas the median ER size was estimated to be 29% of the cell size (Fig 1I, 1J). Therefore, for mRNAs with an unbiased transcript distribution, we expect that typically 11% of transcripts colocalize with TGs and 29% colocalize with the ER.

To determine mRNA transcripts enriched in TG or ER, smRNA-FISH foci were counted using the maxima function and the total number of foci per cell are quantified. Next, all foci are overlaid with the TG mask and the ER mask to identify mRNAs that colocalize with each compartment. To determine if an mRNA is compartment enriched, we tested if its observed compartment distribution differs from the expected distribution based on compartment size using a Mann Whitney test. The code for the image analysis is available (see below).

Of note, this analysis does not distinguish between nuclear and cytoplasmic mRNA localization. For 7/8 mRNAs this does not influence the outcome because the mRNA signal in the nucleus is negligible or non-existent. However, smRNA-FISH probes for endogenous *TES* produce high nuclear background signal. In this case, the prominence value, used to define local maxima to call foci, is increased such that nuclear noise does not substantially influence foci quantification (Fig. S2G).

##### Validation of CY+ mRNAs by smRNA-FISH after digitonin extraction

To distinguish CY+ mRNAs from TG+ or ER+ mRNAs, we performed smRNA-FISH on endogenous mRNAs in untreated and digitonin treated cells, as previously reported. ^57^ The total number of mRNA foci per cell is calculated using the maxima function in FIJI. Next, thresholding is applied to DAPI fluorescence to generate a nuclear mask. Total mRNA foci are overlaid with the DAPI mask and nuclear foci are subtracted from the total, yielding cytoplasmic foci. Cytoplasmic foci are quantified for at least 10 cells per condition per experiment. For each experiment, the mean fraction of transcripts retained is calculated as the average cytoplasmic foci per digitonin-treated cell divided by the average cytoplasmic foci per untreated cell. At least three separate experiments per mRNA were performed.

##### RNA-FISH after transfection of constructs

RNA-FISH experiments probing for GFP-fusion constructs were performed as described previously. ^16^ Stellaris FISH probes for eGFP with Quasar 670 Dye were used.

##### Line profile analysis

To quantify colocalization of ER (GFP-SEC61B) and mRNA (AF633) fluorescence signals, line profiles were generated with FIJI (ImageJ). For each cell, 2-4 straight lines were drawn to cross the ER in different directions, indicated by the white arrows shown in the figures. Fluorescence signal along the straight line of the ER and the mRNA reporter was calculated for each channel using the plot profile tool in FIJI. The values of the Pearson’s correlation coefficient r were calculated using Excel. Perfect correlation of protein-mRNA is indicated by r = 1, perfect exclusion is indicated by r = −1, and random distribution is indicated by r = 0.

#### Confocal microscopy

Confocal imaging was performed using ZEISS LSM 880 with Airyscan super-resolution mode or Nikon CSU-W1 with SoRa super-resolution mode. A Plan-Apochromat 63x/1.4 (Zeiss) or 60x/1.49 (Nikon) Oil objective was used. For live cell imaging, cells were incubated with a LiveCell imaging chamber (Zeiss, Nikon) at 37°C and 5% CO2 and imaged in cell culture media. Excitations were performed sequentially using 405, 488, 594 or 633 nm laser wavelength and imaging conditions were experimentally optimized to minimize bleed-through. Z-stack images were captured with the interval size of 0.2 µm. Images were prepared with FIJI (ImageJ) software.

#### TMT mass spectrometry

To obtain protein expression levels, TMT mass spectrometry analysis was performed on HEK293T cells cultivated in steady-state conditions. Cells were trypsinized and washed three times with ice-cold PBS. Pelleted cells were snap-frozen in liquid nitrogen. Cell pellets were lysed with 200 μl buffer containing 8 M urea and 200 mM EPPS (pH at 8.5) with protease inhibitor (Roche) and phosphatase inhibitor cocktails 2 and 3 (Sigma). Benzonase (Millipore) was added to a concentration of 50 μg/ml and incubated at room temperature for 15 min followed by water bath sonication. Samples were centrifuged at 14,000 g at 4°C for 10 min, and supernatant extracted. The Pierce bicinchoninic acid (BCA) protein concentration assay was used to determine protein concentration. Protein disulfide bonds were reduced with 5 mM tris (2-carboxyethyl) phosphine at room temperature for 15 min, and alkylated with 10 mM iodoacetamide at room temperature for 30 min in the dark. The reaction was quenched with 10 mM dithiothreitol at room temperature for 15 min. Aliquots of 100 μg were taken for each sample and diluted to 100 μl with lysis buffer. Samples were subject to chloroform/methanol precipitation as previously described. ^58^ Pellets were reconstituted in 200 mM EPPS buffer and digested with Lys-C (1:50 enzyme-to-protein ratio) and trypsin (1:50 enzyme-to-protein ratio), and digested at 37°C overnight.

Peptides were TMT-labeled as described. ^58^ Briefly, peptides were TMT-tagged by the addition of anhydrous ACN and TMTPro reagents (16plex) for each respective sample and incubated for 1 hour at room temperature. A ratio check was performed by taking a 1 μl aliquot from each sample and desalted by StageTip method ^59^. TMT tags were then quenched with hydroxylamine to a final concentration of 0.3% for 15 min at room temperature. Samples were pooled 1:1 based on the ratio check and vacuum-centrifuged to dryness. Dried peptides were reconstituted in 1 ml of 3% ACN/1% TFA, desalted using a 100 mg tC18 SepPak (Waters), and vacuum-centrifuged overnight.

Peptides were centrifuged to dryness and reconstituted in 1 ml of 1% ACN/25mM ABC. Peptides were fractionated into 48 fractions. Briefly, an Ultimate 3000 HPLC (Dionex) coupled to an Ultimate 3000 Fraction Collector using a Waters XBridge BEH130 C18 column (3.5 um 4.6 x 250 mm) was operated at 1 ml/min. Buffer A, B, and C consisted of 100% water, 100% ACN, and 25mM ABC, respectively. The fractionation gradient operated as follows: 1% B to 5% B in 1 min, 5% B to 35% B in 61 min, 35% B to 60% B in 5 min, 60% B to 70% B in 3 min, 70% B to 1% B in 10 min, with 10% C the entire gradient to maintain pH. The 48 fractions were then concatenated to 12 fractions, (i.e. fractions 1, 13, 25, 37 were pooled, followed by fractions 2, 14, 26, 38, etc.) so that every 12^th^ fraction was used to pool. Pooled fractions were vacuum-centrifuged and then reconstituted in 1% ACN/0.1% FA for LC-MS/MS.

Fractions were analyzed by LC-MS/MS using a NanoAcquity (Waters) with a 50 cm (inner diameter 75 µm) EASY-Spray Column (PepMap RSLC, C18, 2 µm, 100 Å) heated to 60°C coupled to an Orbitrap Eclipse Tribrid Mass Spectrometer (Thermo Fisher Scientific). Peptides were separated by direct injection at a flow rate of 300 nl/min using a gradient of 5 to 30% acetonitrile (0.1% FA) in water (0.1% FA) over 3 hours and then to 50% ACN in 30 min and analyzed by SPS-MS3. MS1 scans were acquired over a range of m/z 375-1500, 120K resolution, AGC target (standard), and maximum IT of 50 ms. MS2 scans were acquired on MS1 scans of charge 2-7 using isolation of 0.5 m/z, collision-induced dissociation with activation of 32%, turbo scan, and max IT of 120 ms. MS3 scans were acquired using specific precursor selection (SPS) of 10 isolation notches, m/z range 110-1000, 50K resolution, AGC target (custom, 200%), HCD activation of 65%, max IT of 150 ms, and dynamic exclusion of 60 s.

#### Visualization of translation in TGs

The SunTag system was used to visualize mRNA translation in the cytosol and the TGER domain. Stable expression of td-PP7-3xmCherry (Addgene 74926) and scFv-GCN4-sfGFP (Addgene 60907) was achieved by generating virus in HEK293T cells and transducing HeLa cells. Cells were seeded on 3.5 cm glass bottom dishes (Cellvis, D35-20-1-N). 20 hours later, cells were transfected with either the SunTag vector expressing KIF18B (Addgene 74928) or SunTag-FOS-UTR. At 15 hours post transfection, cells were treated with 100 ng/ml doxycycline for one hour to induce SunTag expression. Confocal imaging was performed as described above. Colocalization of foci was quantified using FIJI.

#### mRNA localization-dependent GFP protein expression

##### Transfection

HeLa cells were seeded in 12-well plates at 80% confluency and transfected with 250 ng GFP-THAP1-MS2 and 250 ng of the MCP-mCherry fusion constructs indicated in the figure (Lipofectamine 3000, Invitrogen). When indicated, GFP-THAP1 or GFP-BIRC3-MS2-SU was used instead of GFP-THAP1-MS2. At 13-15 hours post transfection, cells were analyzed by FACS. For RNA-FISH experiments, cells were seeded at 80% confluency in 4-well slide chambers (Millipore Sigma) and cotransfected with 75 ng GFP-THAP1-MS2, 100 ng BFP-SEC61B, and 75 ng of the indicated MCP-mCherry fusion constructs.

##### FACS analysis to measure GFP protein expression

Cells were trypsinized, washed once in complete media, then resuspended in FACS buffer (PBS plus 1% FCS). At least 5,000 cells were measured on a BD LSR-Fortessa Cell Analyzer and FACS data were analyzed using FlowJo software. GFP protein expression corresponds to GFP mean fluorescence intensity (MFI). To determine the effect of MCP-tethered RBPs on protein output of the GFP reporter mRNA, only cells that were successfully cotransfected with both the MCP-mCherry fusion and the GFP reporter constructs were analyzed. To do so, the double-positive cells (mCherry+/GFP+) were gated, and all single positive and unstained cells were excluded from the analysis. The reported GFP MFI was calculated from the double-positive cells. Untransfected cells were used to draw the gates for mCherry+ or GFP+ cells.

##### qPCR analysis to measure *GFP* mRNA abundance

Cells were trypsinized, washed once in complete media, then resuspended in FACS buffer (PBS plus 1% FCS). To determine the effect of MCP-tethered RBPs on GFP reporter mRNA stability, cells were sorted based on expression of both the MCP-mCherry fusion and the GFP reporter constructs. The BD FACSAria III cell sorter was used to collect 50,000 cells from each co-transfected population. Cells were sorted directly into 1 ml of TRIzol solution in Eppendorf tubes for total RNA was extraction. cDNA synthesis was performed on 200 ng of RNA per sample using the SuperScript IV VILO ezDNase Master Mix (Invitrogen). ezDNase enzyme was included to eliminate plasmid DNA contamination. To measure the relative expression levels of reporter mRNA by qRT-PCR, FastStart Universal SYBR Green Master Mix (ROX) from Roche was used together with GFP-qPCR F/R primers. GAPDH was used as a housekeeping gene.

### Data analysis

#### RNA-seq of subcytoplasmic fractions from HEK293T cells

##### RNA-seq

Alignment was generated in Dragen v3.10 (Illumina) against the hg38-alt-masked-v2 reference acquired from GENCODE v43 with default parameters. Gene expression analysis was performed using HOMER v4.11 software. ^60^ The mean RPKM values of all biological replicates were calculated and used for downstream analyses. Only protein-coding genes were analyzed. A gene was considered expressed if the RPKM value is 3 or greater.

##### Classification of membrane/secretory proteins versus non-membrane proteins

Information on the presence of transmembrane domains or a signal sequence was obtained from uniprot. All expressed genes were separated into mRNAs that encode membrane/secretory proteins or non-membrane proteins. If a protein contains a signal sequence but not a transmembrane domain, it is considered as secretory protein. All proteins with transmembrane domains are considered membrane proteins and all remaining proteins are classified as non-membrane proteins. Among the 9155 mRNAs expressed in HEK293T cells, 2140 were classified as membrane/secretory proteins, whereas 7015 were classified as non-membrane proteins (Table S1).

##### Compartment-specific localization scores

The sum of RPKM values obtained from TG particles, ER particles, and the cytosol was considered as total cytoplasmic mRNA expression. For each gene, the mean compartment-specific RPKM value was divided by the total cytoplasmic mRNA expression. As a result, each gene is assigned three localization scores that correspond to the fraction of its transcripts that localize to each of the three compartments: TGs, the ER, and the cytosol.

##### Compartment-specific enrichment of mRNAs that encode membrane/secretory proteins

We considered an mRNA to be ER-enriched if the ratio of localization scores (ER/TG) was greater than 1.25 and classified it as TG-enriched if it was smaller than 0.8. The median localization score of membrane/secretory mRNAs in the cytosol was 0.09. If the cytosolic localization score of an mRNA was greater than 0.36, it was considered enriched in the cytosol. If the ER and TG-specific localization scores were similar and the cytosolic partition coefficient was smaller than 0.18, the mRNA was assigned to the ER, whereas it was considered not localized if the cytosolic localization score was smaller than 0.18 (Fig. S1H).

##### Compartment-specific enrichment of mRNAs that encode non-membrane proteins

To faithfully compare differences in mRNA distribution across the three compartments, it is necessary to know the relative size distribution of the three compartments. However, this parameter is currently unknown. Therefore, instead of comparing the localization scores across samples, we determined the most enriched mRNAs within each compartment. We considered an mRNA compartment-enriched, if its average localization score (from biological replicates) was at least 1.25-fold higher than the median localization score of its corresponding compartment samples. For TG particles, the median localization score was 0.32, for ER particles, it was 0.30, and for the cytosol, the median localization score was 0.34. If the enrichment was observed in two compartments, the mRNA was assigned to the compartment with the higher value. With this strategy, we identified 1246 TG+ mRNAs, 919 non-overlapping ER+ mRNAs, and 1481 CY+ mRNAs. The remaining 3369 mRNAs (48%) do not have a compartment-biased mRNA localization pattern and were called (unbiased).

Justification of the cut-off used to determine compartment-enriched mRNAs. A minimum cut-off of 1.25-fold higher than the median localization score corresponds to approximately one standard deviation. The compartment-enriched mRNAs differed substantially in their functional and architectural features (Fig. 2). We generated subgroups among the compartment-enriched mRNAs that represent the top, middle, and bottom-enriched subgroups (Fig. S4). Even when focusing on the bottom-enriched groups (which are close to the cut-off used), the differences in functional and architectural features across the compartment-enriched groups were still highly significant (Fig. S4). The cut-off is further justified as we were able to validate 10/11 mRNAs considered to be compartment enriched with an independent method. Moreover, we demonstrate that TG-translated MYC has biological effects, despite *MYC* mRNA being found in the bottom enriched TG+ group. ^19^

#### mRNA transcript distribution in HEK293T TIS11B KO cells

We focused on the analysis of mRNAs that encode non-membrane proteins (Table S5). The mean RPKM values of the biological replicates of digitonin-extracted samples and the ER particles were calculated for TIS11B KO cells and their corresponding control HEK293T cells. A gene was considered expressed if the average RPKM value in the ER and in the cytosol samples was greater than 3 RPKM (*N* = 6229). The compartment-specific localization scores were calculated and the difference in localization scores between TIS11B KO and control samples were calculated for ER and cytosol. The top 20% of genes with a localization change towards ER or the cytosol were intersected with genes considered as TG+ (*N* = 1246) and further analyzed with respect to their bound RBPs and architectural features.

#### mRNA and protein features of the localized mRNAs

RPKM values of mRNAs were obtained from RNA-seq data of unfractionated HEK293T cells and were determined for the compartment-biased mRNAs. Pro-seq and RNA-seq from HEK293 cells were obtained from GEO (GSE140365: PRO-seq; GSE142895: RNA-seq). ^27^ Raw reads were processed by trimmomatic (version: 0.39) to trim low-quality ends (average quality per base < 15, 4 bp window) and adapters. ^61^ Trimmed reads were mapped to the human genome (hg19) using hisat2 (version: 2.1.0). ^62^ Reads mapped to each gene were counted by featureCounts (version: 1.6.4). ^63^ To estimate mRNA stability rates, log2-normalized counts of Pro-seq data were divided by the log2-normalized RNA-seq data, as described previously ^28^. 3′UTR length of each mRNA was obtained from Ref-seq. The longest 3′UTR isoform of each gene is reported. mRNA length, CDS length, average CDS exon length, and total exon number of genes were determined using transcripts from the Matched Annotation from the <u>NCBI</u> and <u>EMBL-EBI</u> (MANE) ^64^ human version 1.2. For each gene, the transcript with longest mRNA length was selected. Protein length was calculated by dividing CDS length by three.

#### Proteomics protein expression analysis

Protein expression was obtained from TMT-based quantitative mass spectrometry analysis of HEK293T cells. Precursor protein abundance was calculated for each protein and scaled to the TMT abundance for each channel. Relative abundance was then calculated by averaging the condition-specific biological replicates. In brief, mass spectra were processed using Protein Discoverer 2.5 (ThermoFisher) using the Minora algorithm (set to default parameters) for precursor quantification and using a TMTpro workflow for TMT-based quantification. Database searching included all canonical entries from the human Reference Proteome UniProt database (SwissProt – 2022-03), as well as an in-house curated list of contaminants. The identification of proteins was performed using the SEQUEST-HT engine against the database using the following parameters: a tolerance level of 10 ppm for MS^1^ and 0.6 Da for MS^2^ post-recalibration and the false discovery rate of the Percolator decoy database search was set to 1%. Trypsin was used as the digestion enzyme, two missed cleavages were allowed, and the minimal peptide length was set to 7 amino acids. Carbamidomethylation of cysteine residues (+57.021 Da) was set as static modifications, while oxidation of methionine residues (+15.995 Da) was set as a variable modification. The final protein-level FDR was set to 1%. Precursor abundance quantification was determined based on intensity, and the minimum replicate feature parameter was set at 50%. Proteins were quantified based on unique and razor peptides and proteins with less than two different peptides were excluded. For TMT-based quantification, similar search parameters were used, with the addition of TMTpro tags on lysine residues and peptide N termini (+304.207 Da) set as static modifications. For TMTpro-based reporter ion quantitation, the summed signal-to-noise (S:N) ratio for each TMT channel was extracted, and the closest matching centroid to the expected mass of the TMT reporter ion was found (integration tolerance of 0.003 Da). PSMs with poor quality, MS^3^ spectra with TMT reporter ion channels missing, or isolation specificity less than 0.7, or with less than 70% of SPS masses matching to the identified peptides, or with an average TMT reporter summed signal-to-noise ratio that was less than 10 or had no MS^3^ spectra were excluded from quantification. We exported the results of protein identification and quantification to Excel, including the TMT-based reporter ion quantitation. Additionally, we extracted the MS^1^ precursor abundance for each protein (Minora algorithm), which indicates its relative abundance in the tryptic sample. Each MS^1^-based abundance measured should be a representation of the sum of all the respective TMT-labeled peptides combined. Therefore, for a rudimentary metric of protein abundance across samples, we divided the total MS^1^-abundance for individual proteins by their respective TMT summed signal-to-noise ratio to each TMT channel.

#### CLIP data analysis

##### iCLIP analysis of TIS11B in HEK293T cells

Raw fastq files were demultiplexed using the iCount python package (https://icount.readthedocs.io). 5′ and 3′ adapters were trimmed by Cutadapt. ^65^ Trimmed reads were mapped to human genome using STAR and reads mapping to tRNA/rRNA were discarded. ^66^ Crosslink sites were called from bam files using the “xlsites” function of iCount. CLIP-seq analysis was carried out on the iMaps platform (https://imaps.genialis.com/iclip), where peak calling was performed by analysing cDNA counts at crosslink sites using Paraclu. ^67^ Motif analysis was carried out using HOMER software. Enrichment was calculated within the genomic coordinates of a total of 57,714 TIS11B CLIP peaks found in 3′UTRs. Total peaks: 190,920; peaks in 3′UTRs: 57,714.

##### POSTAR3 CLIP data

CLIP data on 168 RBPs were downloaded from Postar3 ^34^ and peak counts that overlapped with annotated 3′UTRs from Ref-seq in all mRNAs that encode non-membrane proteins were recorded. For each RBP, the median number of 3′UTRs CLIP peaks was calculated and all 3′UTRs with peaks counts greater than the median were considered as targets. Based on the fraction of mRNAs that are considered compartment-specific (TG: 17.8%; ER 13.1%; CY: 21.1%; unbiased: 48.0%), we determined the expected number of target genes for each compartment. If the observed number of targets divided by the expected number of targets in a compartment was greater than 1.5, the RBP was added to our short-list (Table S4). As TIS11B and TIA1/L1 are known to bind to AU-rich sequences, we added the processed PAR-CLIP data of the LARP4B RBP as it was reported to bind to AU-rich elements. ^33^

##### Logistic regression

The R package ‘nnet’ (v7.3-17) was used to fit logistic regression models to predict the subcytoplasmic mRNA localization of non-membrane proteins. An initial model used CLIP peak counts from the RBPs on the short list (*N* = 24). A second model used the top seven RBPs from the first model fit and added mRNA length and average CDS exon length. Covariates with missing values were imputed as zeros. All covariates were first ‘sqrt’ transformed and then standardized. The ‘unbiased’ category was used as the base level. The R package ‘broom’ (v0.8.0) was used to compute t-test statistics for the model coefficients. The code is available on github (github.com/Mayrlab/tiger-seq).

##### Confirmation of the logistic regression

To validate the contribution of each individual RBP, we used more stringent criteria to determine their targets. Among all mRNAs that encode non-membrane proteins with at least one CLIP peak in the 3′UTR, we considered the top third of mRNAs as targets of each RBP (TIS11B: 1781 targets; TIA1/L1: 1313 targets; LARP4B: 1621 targets; METAP2: 256 targets; HuR: 1124 targets; PUM2: 427 targets; HNRNPC: 232 targets). mRNAs only bound by LARP4B or METAP2 are LARP4B/METAP2 targets and not bound by another RBP (from the seven RBPs investigated), *N* = 717. mRNAs predominantly bound by TIS11B are TIS11B targets exclusively bound by TIS11B or co-bound by TIA1/L1, with TIS11B/TIA1/L1 ≥ 2 (*N* = 834). mRNAs predominantly bound by TIA1/L1 are TIA1/L1 targets exclusively bound by TIA1/L1 or co-bound by TIS11B but TIS11B/TIA1/L1 < 2 (*N* = 634).

#### Intersection of membrane/secretory mRNAs with previous datasets

<u>APEX-seq.</u> The mRNAs that are coexpressed in our RNA-seq dataset (*N* = 9155 mRNAs) and the ER membrane-localized mRNAs from the APEX-seq dataset (*N* = 1045) were determined. ^11^ The overlapping 845 mRNAs were intersected with the mRNAs that encode membrane/secretory proteins found to be ER+ in our analysis (*N* = 1476). We detected 673 mRNAs which correspond to 80% of all APEX-seq mRNAs that are considered to be ER membrane (ERM)-enriched. The universe used to test for enrichment were all mRNAs that encode non-membrane proteins (*N* = 2140). <u>Biochemical fractionation.</u> A similar analysis was performed for the fractionation dataset from Reid (2012). ^9^ Among the 385 coexpressed mRNAs that are enriched on the ER according to Reid, we detected 308 in our ER+ fraction when focusing on membrane/secretory protein encoding mRNAs. This group represents 80% of all ER-enriched mRNAs detected by Reid. <u>MERFISH.</u> In the MERFISH dataset, which was generated in U2OS cells, 1037 mRNAs are considered ER-enriched. Among them, *N* = 571 are co-expressed in our dataset and considered mRNAs encoding membrane/secretory proteins. Among the 571 co-expressed mRNAs we consider 511 as ER+, which corresponds to 89%. Among the ER-de-enriched mRNAs (Log2FC nonER vs ER = −0.34), only 69 mRNAs encode membrane/secretory proteins. Among the 69 mRNAs, we consider 8 as ER+, which corresponds to 11.6%. ^13^

#### Intersection of mRNAs that encode non-membrane proteins with a previous dataset

The relative distribution of mRNA transcripts across subcellular compartments, including the membrane fraction, phase-separated granules, and the cytosol was determined using density gradient centrifugation in U2OS cells. ^25^ The number of co-expressed mRNAs that encode non-membrane proteins was *N* = 6557, which corresponds to 93% of our dataset. This dataset determines the proportion of transcripts that localize to the different fractions. For co-expressed TG+ mRNAs (*N* = 1153), ER+ mRNAs (*N* = 839) and CY+ mRNAs (*N* = 1400), we plotted the proportion of mRNAs that localize to phase-separated granules, to the membrane fraction, and to the cytosol in the LoRNA dataset in U2OS cells.

#### Gene ontology analysis

Gene ontology (GO) analysis was performed using DAVID. ^30^

#### Further statistical analysis

Statistical parameters are reported in the figures and figure legends, including the definitions and exact values of *N* and experimental measures (mean ± std or boxplots depicting median, 25^th^ and 75^th^ percentile (boxes) and 5% and 95% confidence intervals (error bars). Pair-wise transcriptomic feature comparisons were performed using a two-sided Mann-Whitney test. For more than two samples, a Kruskal-Wallis test was performed. For transcriptomic analyses, statistical significance is indicated by asterisks *, 0.05 > *P >* 1 × 10^−9^; **, 1 × 10^−10^ > *P >* 1 × 10^−20^; ***, 1 × 10^−21^ > *P >* 1 × 10^−80^; ****, 1 × 10^−81^ > *P >* 0. Exact *P* values are listed in Table S3. Enrichment was determined using a Χ^2^ test. The *P* value was calculated using a two-sided Fisher’s exact test. When indicated, a two-sided t-test with assumption of equal variance was applied. Statistical significance for experimental data is indicated by asterisks *, *P* < 0.05, **, *P* < 0.01, ***, *P* < 0.001, ****, *P* < 0.0001.

#### Data and code availability

The data of the proteomics experiment were deposited in the MassIVE repository (dataset identifier MSV000092176). The RNA-seq samples obtained from the subcytoplasmic fractionation and the TIS11B iCLIP data obtained from HEK293T cells are available at GEO (Accession number: GSE215770). The code for logistic regression is available on github (github.com/Mayrlab/tiger-seq). Raw western blot data, raw imaging data and scripts for analysis are deposited at Mendeley (https://data.mendeley.com/datasets/nmt7ppsp8r/1).

## Supplementary Figures

**Figure S1. Strategy to determine subcytoplasmic mRNA localization.**

S1A. Cell fractionation strategy to obtain the cytoplasmic membrane fraction. Transfected HEK293T cells were lysed in hypotonic buffer, followed by douncing and differential centrifugation at 600 g to pellet nuclei. The supernatant was subsequently spun at 7000 g. The pellet contains the cytoplasmic membrane fraction that was used for subsequent fluorescent particle sorting.

S1B. FACS plot showing the gating strategy to obtain mCherry-TIS11B+/GFP-SEC61B+ TG particles and mCherry-TIS11B−/GFP-SEC61B+ ER particles. The particles were costained with DAPI to segregate TG and ER particles from nuclear contamination (DAPI high). The DAPI low particles were sorted.

S1C. Immunoblot showing markers used to evaluate the quality of the three compartment fractions. WCL is the unfractionated whole cell lysate. CY is the digitonin-extracted cytosol. Pre-sort is the membrane-enriched cytoplasmic lysate from which TG and ER particles are sorted, which is the final step in (A). 50K TG indicates 50,000 sorted TG particles and 50K ER indicates 50,000 sorted ER particles. H3 antibody was used as a marker for nuclear components, GAPDH was used as marker for cytosolic proteins, and Calnexin and GFP-SEC61B were used as ER markers. *, signal from mCherry-TIS11B.

S1D. Quantification of endogenous TIS11B and mCherry-TIS11B from (C) together with additional biological replicates. Shown is the ratio of protein abundance in 50K TG over 50K ER as mean ± std of three independent experiments.

S1E. As in (D), but quantification of endogenous Calnexin and GFP-SEC61B. Shown is the ratio of protein abundance in 50K ER over 50K CY as mean ± std of three independent experiments.

S1F. Correlation of log2-transformed RPKM values obtained by RNA-seq for biological replicates of subcytoplasmic compartments. The Pearson correlation coefficients are shown.

S1G. Baseline distribution of localization scores across the three investigated cytoplasmic compartments is shown separately for mRNAs that encode membrane/secretory proteins and mRNAs that encode non-membrane proteins.

S1H. Distribution of localization scores in each fractionation sample for compartment-enriched mRNAs that encode membrane/secretory proteins.

S1I. Overlap of ER+ mRNAs that encode membrane/secretory proteins (*N* = 1476) defined by us (particle sorting) with previous datasets that used alternative isolation methods. In the APEX-seq dataset 78% of ER-localized mRNAs overlap with our ER+ mRNAs. In the fractionation dataset, 80.5% of ER-localized mRNAs overlap with our ER+ mRNAs. Among the MERFISH ER-localized mRNAs 89% overlap with our ER+ mRNAs and only 11.6% of our ER+ mRNAs overlap with the ER-de-enriched mRNAs obtained by MERFISH.

S1J. ER+ mRNAs that encode membrane/secretory proteins (*N* = 1476) defined by us are significantly enriched on the ER membrane (ERM) according to APEX-seq, whereas CY+ mRNAs that encode membrane/secretory proteins show a significantly lower ERM enrichment. Similarly, CY+ mRNAs that encode membrane/secretory proteins have significantly higher APEX2 enrichment scores in the cytosol than mRNAs considered ER+ by us. Mann Whitney test: *P* < 2×10^−7^.

S1K. Ternary plot showing compartment-enriched mRNAs. Each dot represents an mRNA that is color-coded as in Fig. 1D-F. mRNAs in the center (light grey) are considered to have an unbiased transcript distribution.

S1L. Distribution of localization scores in each fractionation sample for compartment-enriched mRNAs that encode non-membrane proteins.

S1M. Validation of mRNAs that encode non-membrane proteins and are defined as compartment-enriched by us in comparison with the LoRNA dataset. Our TG+ mRNAs are highest enriched in the mRNAs that LoRNA identifies in the phase-separated granule fraction. Our ER+ mRNAs are highest enriched in the mRNAs that LoRNA identifies in the membrane fraction. Our CY+ mRNAs are highest enriched in the mRNAs that LoRNA identifies in the cytosol. Mann Whitney test was performed. Exact *P* values are listed in Table S3.

**Figure S2. Validation of endogenous TG+ and ER+ mRNAs by smRNA-FISH.**

S2A. SmRNA-FISH of endogenous TG+ mRNA *DUSP1* (green) in HeLa cells. TGs (BFP-TIS11B, blue) and the ER (GFP-SEC-61B, magenta) were simultaneously visualized. The maximum projection of the fluorescent signals is shown. Bottom panel shows a 5x zoom-in of the area indicated by the white dashed box. White circles indicate colocalization of mRNA puncta with TGs, whereas dashed circles indicate colocalization with the ER. Representative images are shown. Scale bar, 5 µm.

S2B. As in (A), but smRNA-FISH of endogenous TG+ mRNA *DNAJB1* is shown. S2C. As in (A), but smRNA-FISH of endogenous ER+ mRNA *PLA2GA4* is shown. S2D. As in (A), but smRNA-FISH of endogenous ER+ mRNA *IDH1* is shown.

S2E. As in (A), but smRNA-FISH of endogenous ER+ mRNA *TES* is shown. S2F. As in (A), but smRNA-FISH of endogenous ER+ mRNA *ACTN4* is shown.

S2G. As in (E) but shown is a 2x magnification of the area indicated by the white dashed box. The image illustrates the high background signal in the nucleus observed when probing for endogenous *TES*. For quantification of this mRNA, the noise tolerance value used in the puncta-calling function was increased to limit nuclear background noise. The white circles indicate nuclear foci that are included in the quantification at this tolerance level. Scale bar, 5 µm.

S2H. Colocalization of TGs and smRNA-FISH foci of three TG+ (beige) and five ER+ (blue) endogenous mRNAs. The beige box plot indicates the expected fraction of mRNA transcripts based on the TG compartment size distribution, obtained from 186 cells. Number of cells analyzed for RNA-FISH: *BAG3* (*N* = 17), *DUSP1* (*N* = 30), *DNAJB1* (*N* = 25), *PLA2GA4* (*N* = 17), *IDH1* (*N* = 18), *ALDH18A1* (*N* = 22), *TES* (*N* = 34), *ACTN4* (*N* = 23). Mann Whitney test shows that the mRNA transcript distribution of 3/3 TG+ mRNAs (beige) is significantly higher than what would be expected for unbiased mRNAs. *P* value categories as in Figure 2A, exact *P* values listed in Table S3.

S2I. As in (H), but colocalization of the ER and smRNA-FISH foci of three TG+ (beige) and five ER+ (blue) endogenous mRNAs. The blue box plot indicates the expected fraction of mRNA transcripts based on the ER compartment size distribution, obtained from of 186 cells. Mann Whitney test shows that the mRNA transcript distribution of 4/5 ER+ mRNAs (blue) is significantly higher than what would be expected for unbiased mRNAs.

**Figure S3. Validation of CY+ mRNAs by smRNA-FISH after digitonin extraction.**

S3A. Shown are smRNA-FISH foci of endogenous *SF3A2* mRNA and *MAP2K2* mRNA in HeLa cells before (−) and after (+) digitonin extraction. *SF3A2* and *MAP2K2* are considered CY+ mRNAs. Cell boundaries are indicated by the dotted lines. Representative images are shown. Scale bar, 5 µm.

S3B. As in (A), but smRNA-FISH foci of endogenous *TES* mRNA and *IDH1* mRNA are shown, which are considered ER+ mRNAs.

S3C. As in (A), but smRNA-FISH foci of endogenous *DUSP1* mRNA and *DNAJB1* mRNA are shown, which are considered TG+ mRNAs.

S3D. Schematic of the reporter mRNA used with the SunTag system to measure nascent protein synthesis. CDS, coding sequence. The KIF18B construct was used previously. ^32^

S3E. Confocal imaging of HeLa cells stably expressing SunTag reporter proteins svFc-GFP and mCherry-tagged PP7 protein (mC-PP7) co-transfected with two constructs (i) BFP-TIS11B to visualize TGs and (ii) SunTag-labeled mRNA encoding KIF18B with PP7-binding sites in the 3′UTR. The *KIF18B* mRNA is visualized by mC-PP7 binding (teal) whereas the KIF18B protein is visualized by svFc-GFP binding (magenta). Foci with co-localization of mRNA and protein represent nascent protein synthesis and are indicative of active translation. A representative example is shown. The white box indicated area is shown at 6x magnification in the lower panel. White arrows indicate actively translating mRNA, yellow arrow indicates non-translating mRNA. Scale bar, 5 µm (top panel), 1 µm (bottom panel).

S3F. Quantification of the experiment from (E). Shown are the number of mRNA foci in TGs or the cytosol (CY) using the Suntag reporter from (D). Each dot represents a cell*. N* = 24 cells were analyzed.

S3G. As in (F), but shown are the mRNAs that are actively translated in each compartment, which were identified by counting the teal and magenta-double positive foci.

**Figure S4. Characteristics of compartment-enriched mRNAs are shown for subgroups.**

S4A. To justify the cut-off used to determine compartment-enriched mRNAs (Fig. 1D-F), we show the data from Figure 2 in more detail. Steady-state mRNA abundance levels obtained from whole cell lysates is shown for unbiased mRNAs (*N* = 3369), three subgroups of TG+ mRNAs (top, middle, bottom, *N* = 415 each); three subgroups of ER+ mRNAs (top, middle, bottom, *N* = 306 each); three subgroups of CY+ mRNAs (top, middle, bottom, *N* = 493 each). Even when focusing on the bottom-enriched groups (which are close to the cut-off used), the differences across the compartment-enriched groups are still highly significant. Mann Whitney tests were performed. *P* value categories as in Fig. 2A. Exact *P* values are shown in Table S3. RPKM, reads per kilobase of transcript per million reads mapped.

S4B. As in (A), but steady-state protein levels obtained from whole cell lysates are shown.

S4C. As in Fig. 2C, but steady-state mRNA abundance levels of compartment-enriched mRNAs obtained by RNA-seq from whole cell lysates of HEK293 cells are shown. This sample was used together with the Pro-seq sample to estimate mRNA half-lives.

S4D. As in (A), but Pro-seq levels are shown.

S4E. As in (A), but estimated mRNA half-lives are shown. S4F. As in (A), but protein size distributions are shown.

S4G. As in (A), but mRNA length distributions are shown. S4H. As in (A), but 3′UTR length distributions are shown.

S4I. As in (A), but average CDS exon length distributions are shown. S4J. As in Fig. 2A, but the number of exons per mRNA is shown.

S4K. As in (A), but the number of exons per mRNA is shown.

**Figure S5. Analyses of TIS11B CLIP data and TIS11B KO samples.**

S5A. Gel showing samples used for iCLIP of GFP-tagged TIS11B. The region outlined in red was used for iCLIP sample preparation.

S5B. TIS11B iCLIP tag distribution obtained from HEK293T cells.

S5C. The top five motifs that were enriched within TIS11B peaks in 3′UTRs compared to all nucleotides in 3′UTRs. Shown are *P* values obtained by HOMER.

S5D. The fraction of mRNAs bound by at least one RBP (from Fig. 3A) for the different groups of compartment-enriched mRNAs is shown.

S5E. Immunoblot of TIS11B in control cells and TIS11B KO HEK293T cells. H3 was used as loading control.

S5F. Correlation of log2-transformed RPKM values obtained by RNA-seq for biological replicates of sorted ER particles or digitonin-extracted cytosol samples for control and TIS11B KO cells. The Pearson correlation coefficients are shown.

**Figure S6. mRNA localization-dependent protein expression of the GFP reporter.**

S6A. Gating strategy to assess GFP protein expression of the reporter mRNA by FACS. Left panel shows the ungated population of HeLa cells coexpressing MCP-mCherry and the *GFP-THAP1-MS2* reporter, separated by size (forward scatter) and granularity (side scatter). The black circle indicates the live cells that were used for subsequent analysis. Middle panel, the GFP-and mCherry-double positive population was gated to obtain the GFP mean fluorescence values (MFI, right panel) which corresponds to the reported GFP protein expression values.

S6B. As in (A), but HeLa cells coexpressing MCP-mCherry-TIAL1 and the *GFP-THAP1-MS2* reporter.

S6C. As in Fig. 4F and 5C, but the MS2 sites in the GFP reporter were omitted. Top: coexpression of MCP-mCherry-TIAL1 does not result in the binding of MCP to the reporter mRNA without MS2 sites. This experiment serves as control for the effect of TIA1L1 overexpression on reporter mRNA expression. Bottom: Quantification of the experiment with the GFP mRNA reporter lacking the MS2 binding sites. Shown is the mean ± std of three independent experiments. T-test for independent samples, NS, not significant.

S6D. Schematic of a second mRNA reporter used to validate the effect of a single 3′UTR-bound RBP on protein expression. The *GFP-BIRC3* reporter mRNA contains the BIRC3 coding region and MS2 hairpins as 3′UTR, which allow binding of the co-transfected MS2 coat protein (mCherry-tagged MCP). Fusion of TIAL1 to MCP tethers TIAL1 to the 3′UTR of the reporter mRNA. mC, mCherry.

S6E. GFP protein expression of the reporter mRNA from (D) in HeLa cells, coexpressing the indicated MCP-fusion constructs, measured by FACS. Representative histograms are shown. GFP-negative cell populations are shown as dotted lines.

S6F. Quantification of the experiment shown in (E). Shown is the mean ± std of five independent experiments. T-test for independent samples, **, *P* = 0.005.

S6G. RNA-FISH of the GFP reporter mRNA (teal) from Fig. 5E in HeLa cells coexpressing MCP-mCherry-SEC61B (magenta) to visualize colocalization between the mRNA and the rough ER membrane. Representative confocal images are shown. Scale bar, 5 µm.

S6H. Line profiles of the fluorescence intensities of the arrows from (G).

S6I. Quantification of the experiment from (G). Two line profiles were generated for each cell. The Pearson’s correlation coefficients between the reporter mRNA and the ER were determined. For MCP, *N* = 26 cells were analyzed, for MCP-SEC61B, *N* = 26 cells were analyzed. The horizontal line denotes the median and the error bars denote the 25^th^ and 75^th^ percentiles. Mann-Whitney test, ****, *P* < 0.0001.

S6J. Schematic of a second GFP-tagged mRNA reporter that investigates the influence of subcellular mRNA localization on protein expression. Fusion of MCP to TRAPα localizes the GFP reporter mRNA to the ER membrane, whereas MCP alone localizes it to the cytosol.

S6K. Confocal live cell imaging of HeLa cells expressing mC-tagged TRAPα-MCP. Scale bar, 5 µm.

S6L. GFP protein expression of the reporter mRNAs from (J) coexpressing the indicated MCP-fusion constructs in HeLa cells measured by FACS. Representative histograms are shown. The histograms on the left indicate GFP-negative cell populations.

S6M. Quantification of the experiment from (L). Shown is the mean ± std of four independent experiments. T-test for independent samples, ****, *P* < 0.0001.

**Figure S7. Redirecting mRNA localization from the cytosol to the rough ER overcomes the repressive effect of a bound RBP.**

S7A. Confocal live cell imaging of HeLa cells expressing the indicated constructs. Shown are representative images with TGs or cytosolic TIS11B. mC, mCherry. Scale bar, 5 µm.

S7B. Quantification from (A). The fraction of HeLa cells with TGs is shown after transfection of the indicated TIS11B fusion constructs. *N* = 165 cells were analyzed for mCherry-TIS11B and *N* = 198 cells were analyzed for MCP-mCherry-TIS11B. MCP-mCherry-TIS11B largely prevents TG formation.

S7C. Schematic of a second TIS11B-bound mRNA reporter that allows investigation of mRNA localization-dependent GFP expression. The coding region of the reporter is provided by BIRC3, followed by MS2 binding sites. Tethering of MCP or TIS11B localizes the mRNA reporter to the cytosol, whereas the MCP-TIS11B-SEC61B fusion localizes the mRNA reporter to the rough ER.

S7D. GFP protein expression of the reporter mRNA from (C) in HeLa cells, coexpressing the indicated MCP-fusion constructs, measured by FACS. Representative histograms are shown. The histograms on the left indicate GFP-negative cell populations.

S7E. Quantification of the experiment from (D). Shown is the mean ± std of four independent experiments. T-test for independent samples, ****, *P* < 0.0001, MCP vs TIS11B-SEC61B: *P* = 0.058; NS).

