## Supplemental Figures for "Subcytoplasmic location of translation controls protein output"

Horste et al., Figure S1

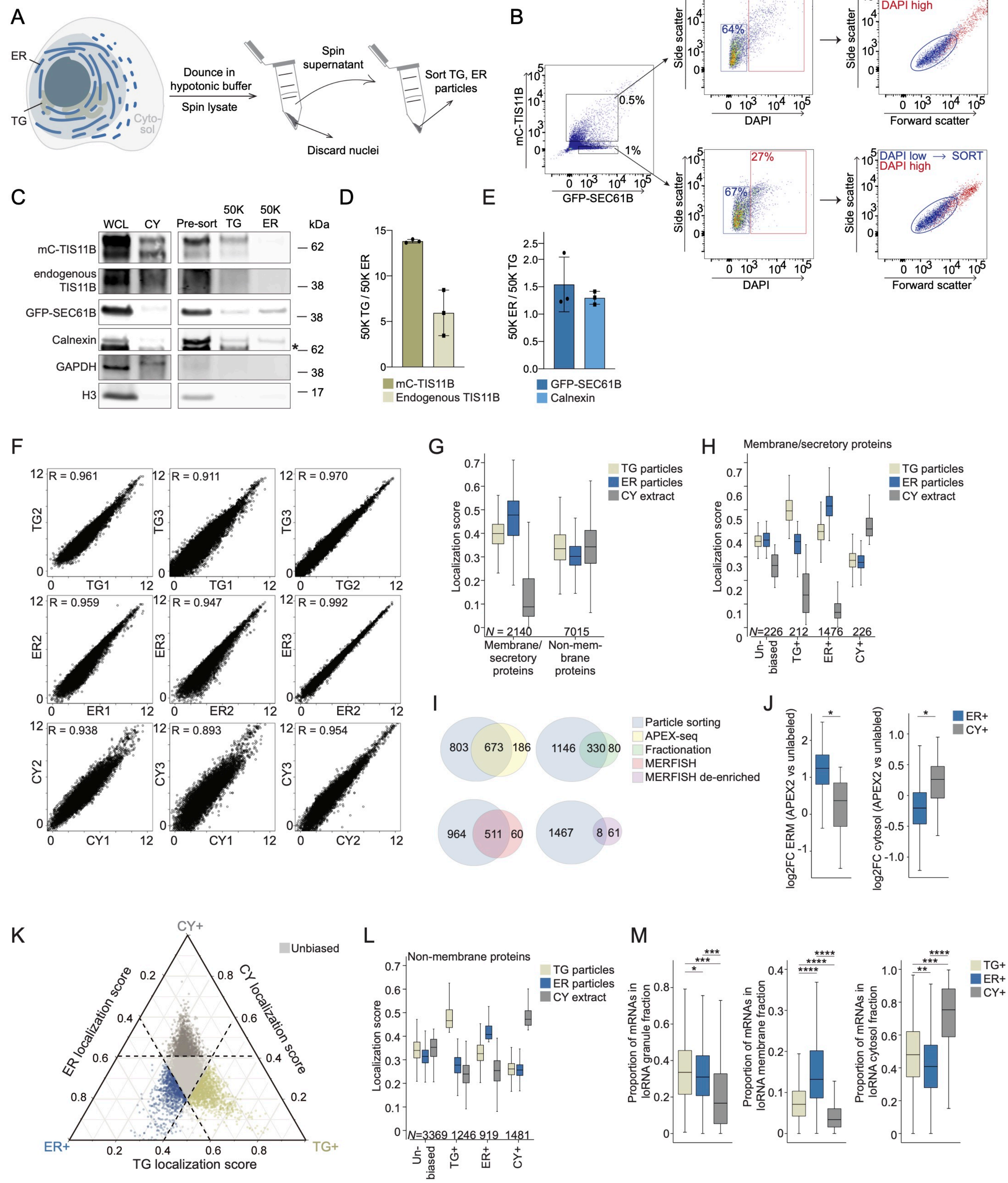

Horste et al., Figure S2

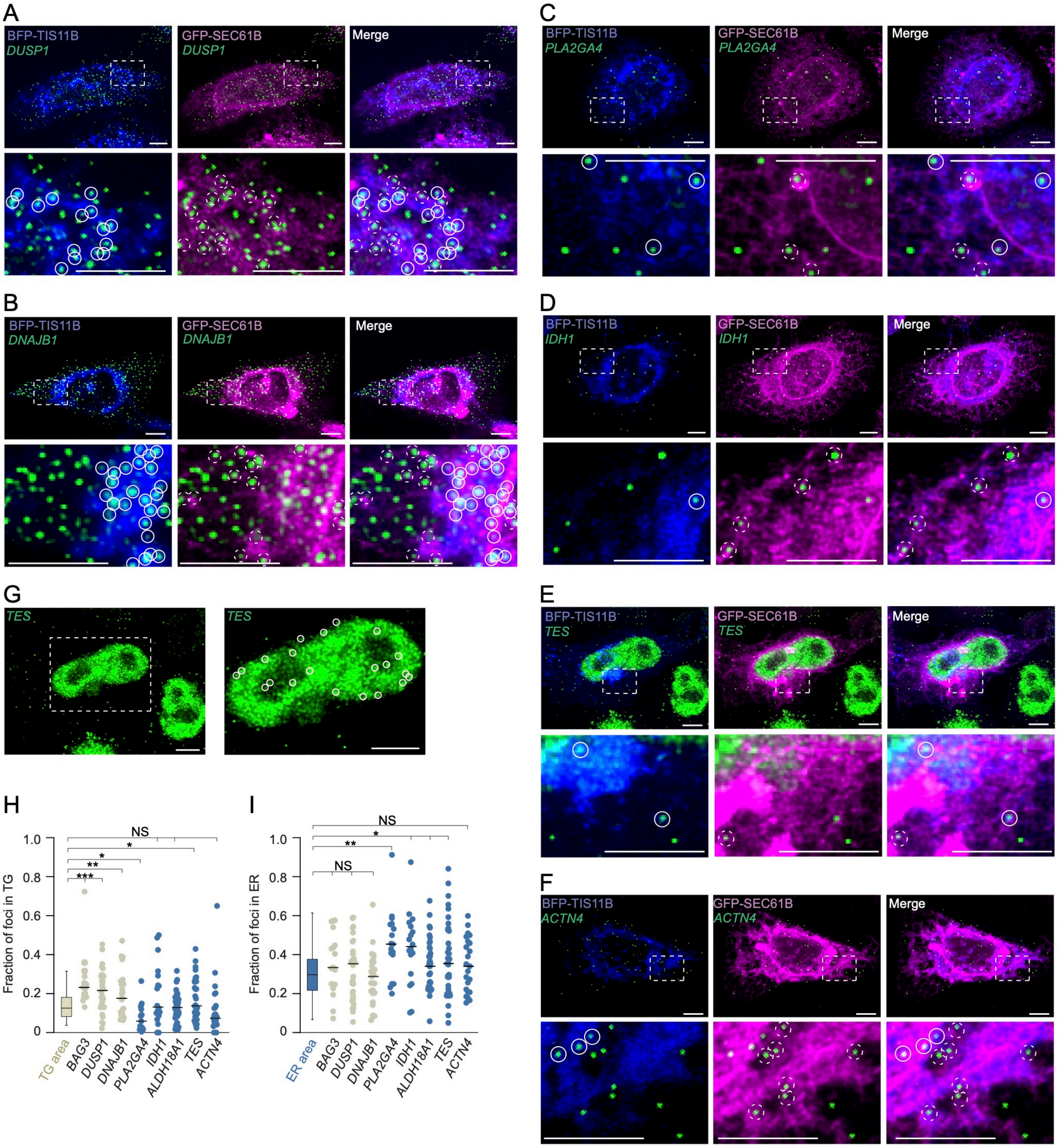

Horste et al., Figure S3

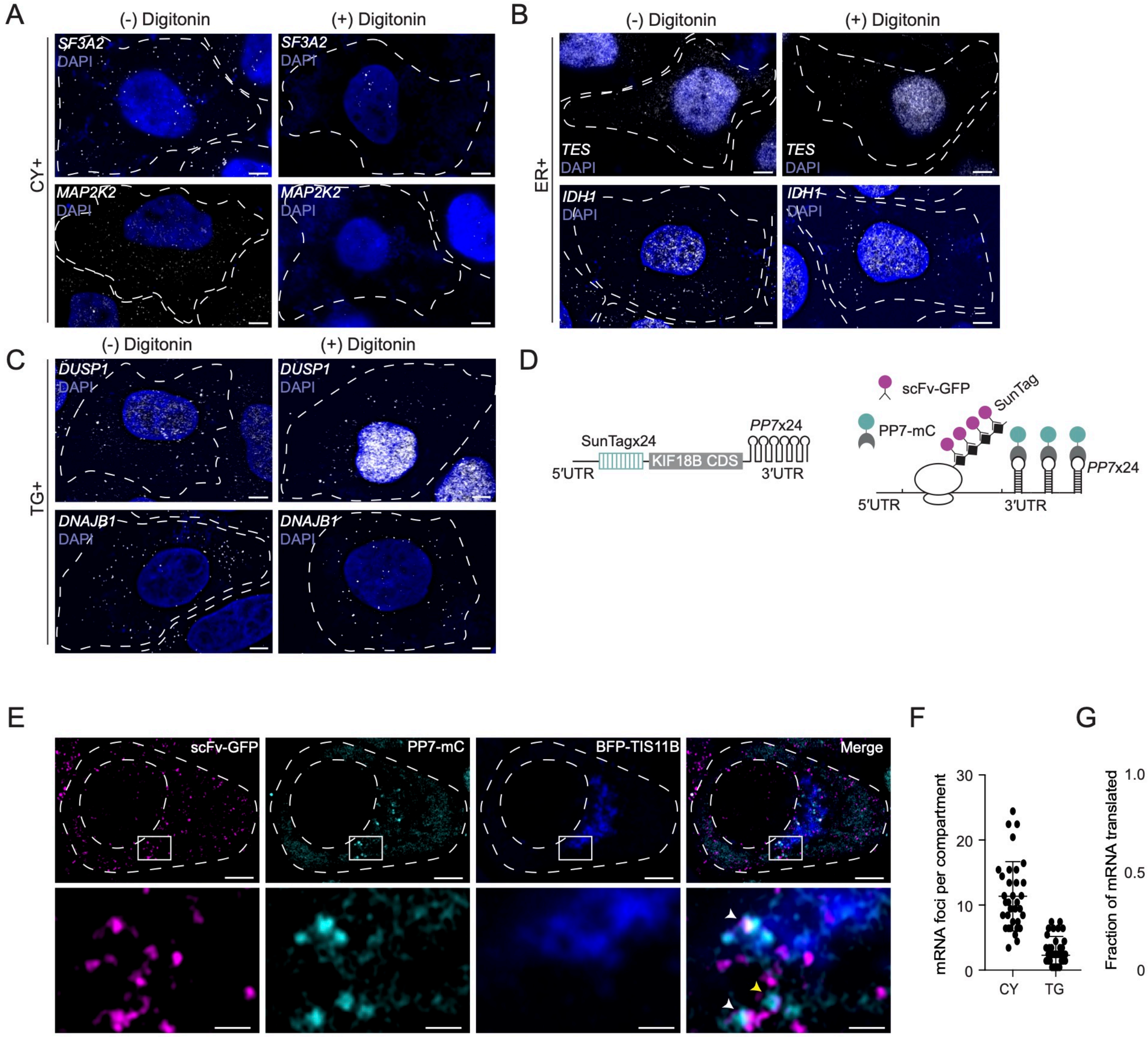

Horste et al., Figure S4

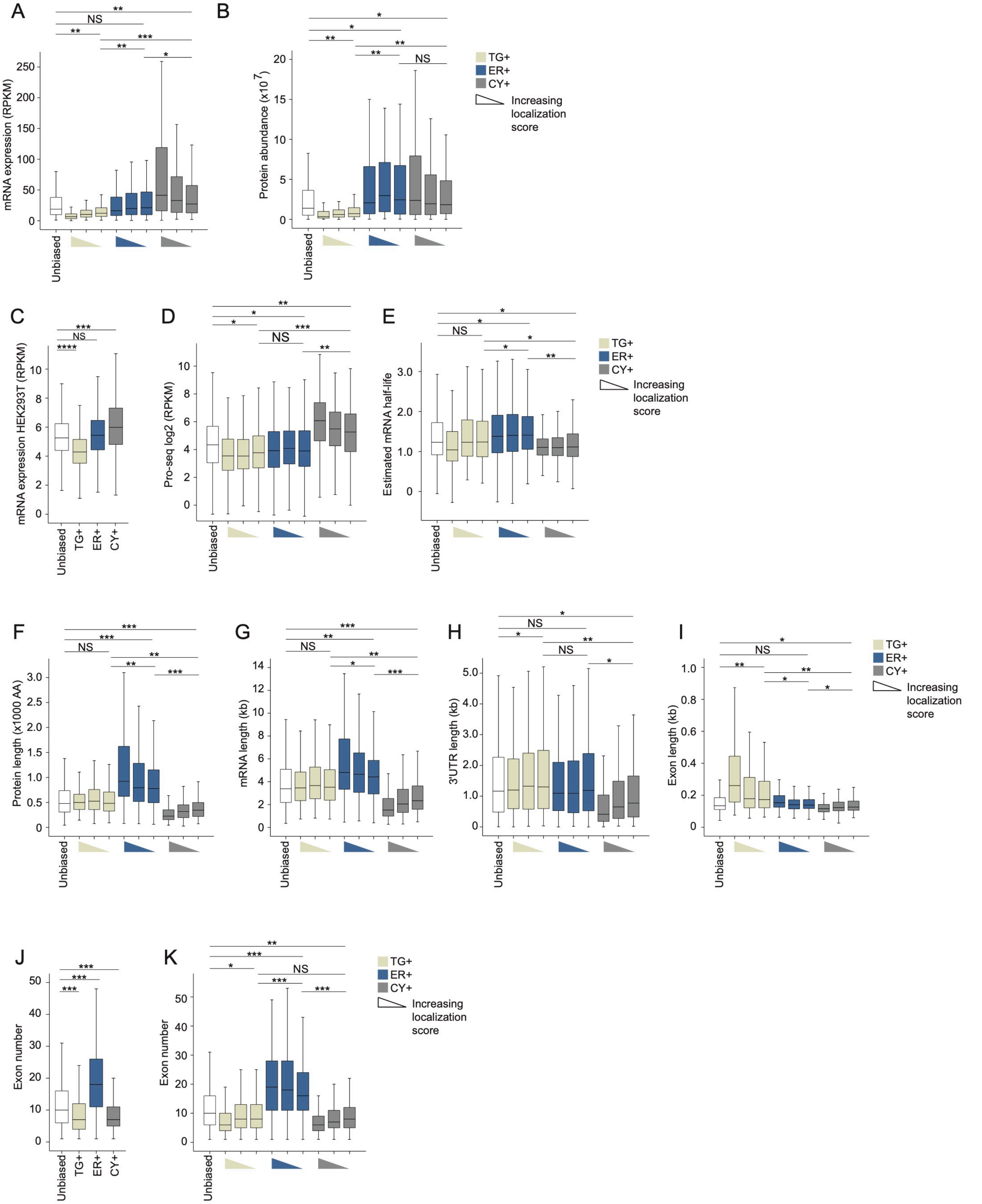

Horste et al., Figure S5

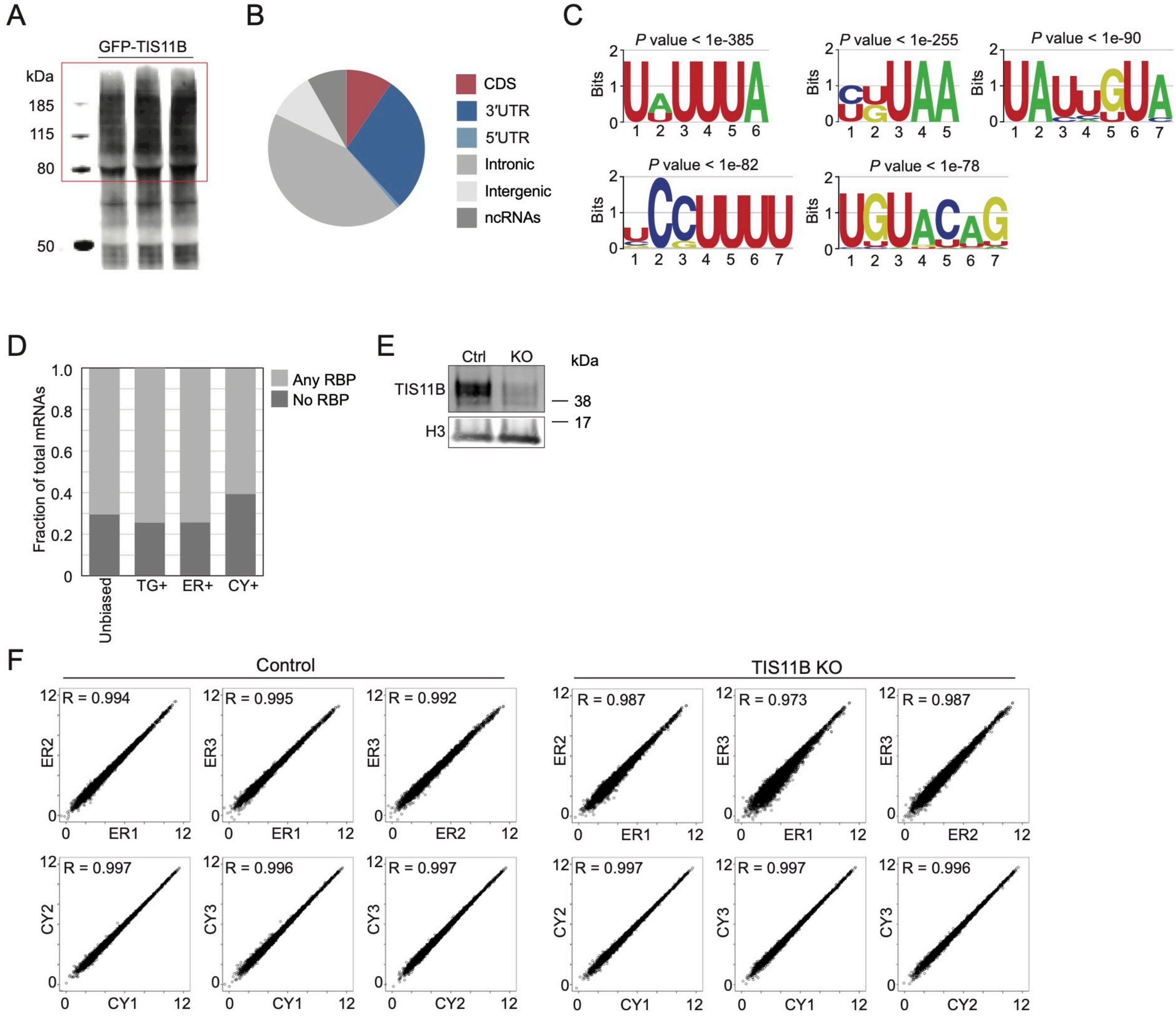

Horste et al., Figure S6

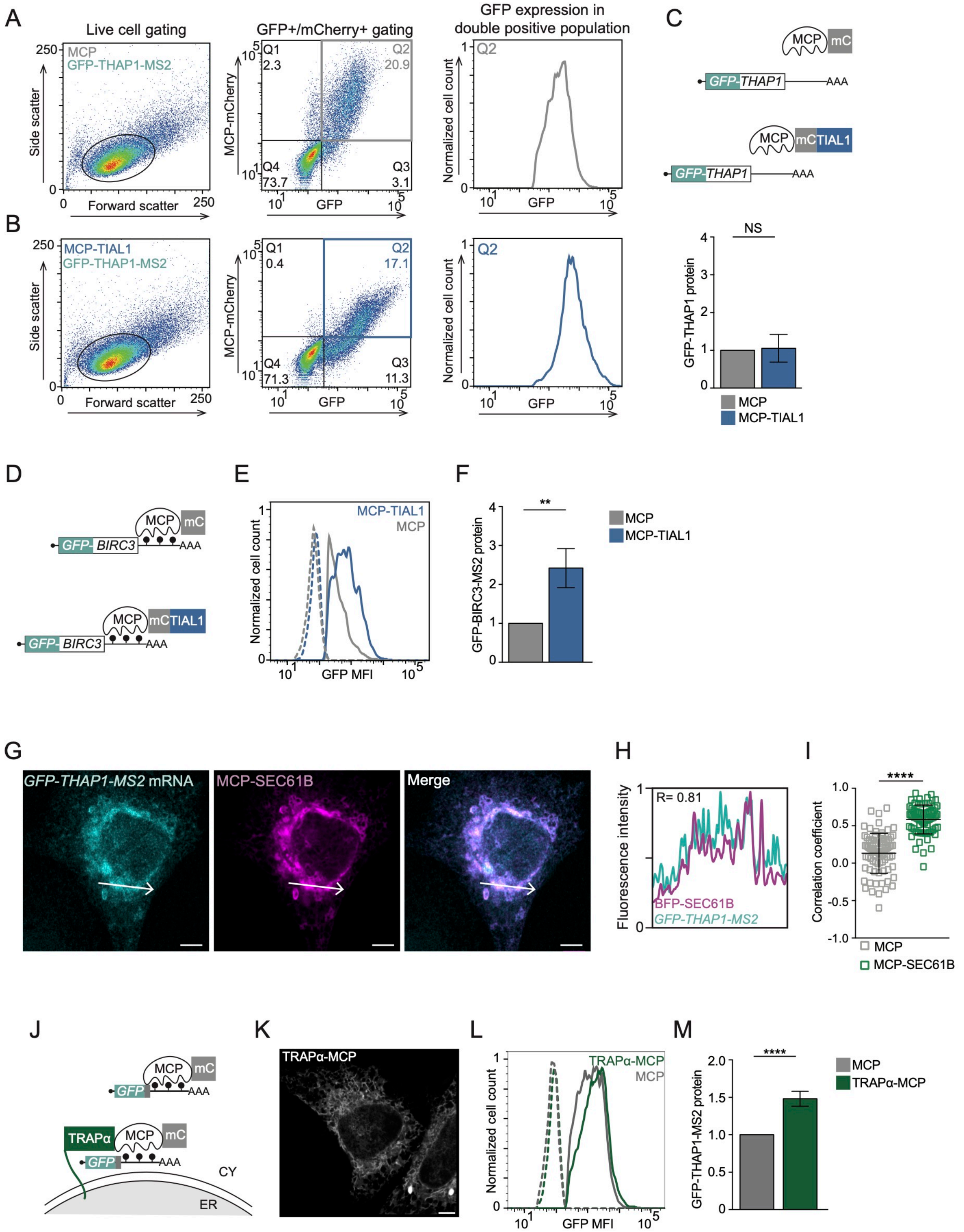

Horste et al., Figure S7

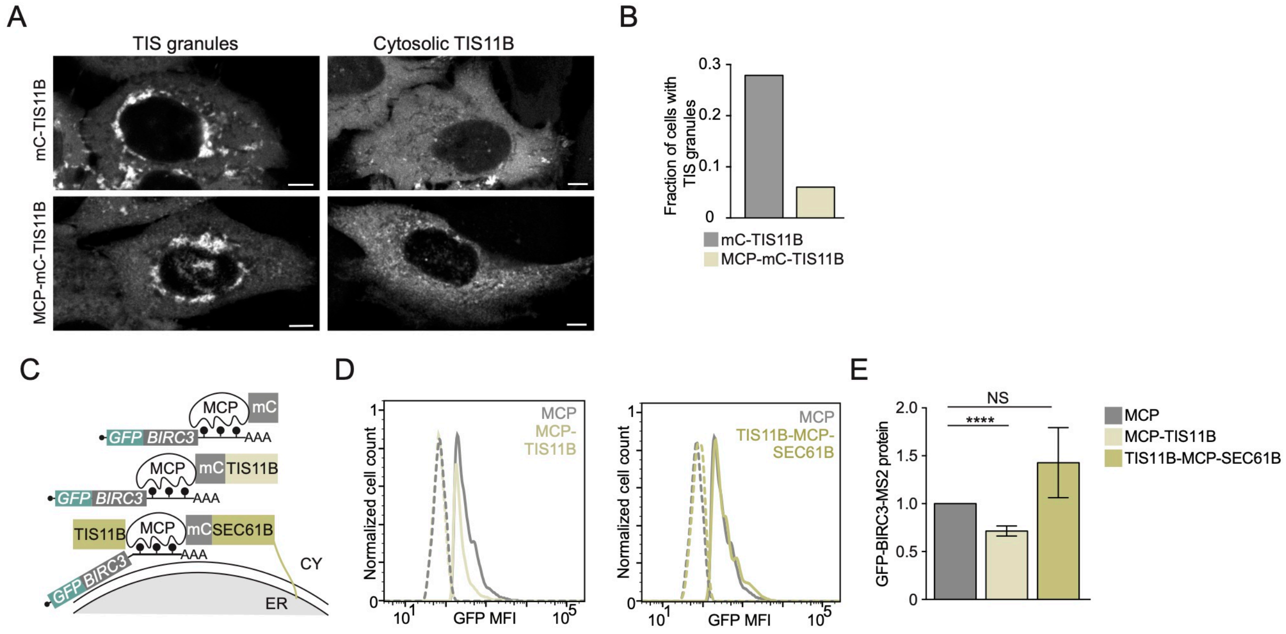
