## Supplementary material for "Subcytoplasmic location of translation controls protein output": Table S3

Table S3. Mann-Whitney statistical test values

| **Related to Figure** | **Comparison** | ***Z* Score** | ***P* Value** |
| --- | --- | --- | --- |
| 1D | Unbiased vs TG | -51.9 | 0 |
| 1E | Unbiased vs ER | -46.1 | 1.0E-123 |
| 1F | Unbiased vs CY | -55.5 | 0 |
| 1I | TG area proportion vs TG foci proportion | -6.6 | 5.1E-11 |
| 1J | ER area proportion vs ER foci proportion | -7.4 | 1.1E-6 |
| 2A | Unbiased vs TG+ | -20.6 | 5.5E-94 |
| 2A | Unbiased vs ER+ | -0.87 | 0.39 |
| 2A | Unbiased vs CY+ | -5.0 | 8.9E-40 |
| 2B | Unbiased vs TG+ | -11.6 | 3.5E-31 |
| 2B | Unbiased vs ER+ | -7.4 | 1.5E-13 |
| 2B | Unbiased vs CY+ | -5.0 | 5.4E-7 |
| 2C | Unbiased vs TG+ | -10.6 | 2.5E-26 |
| 2C | Unbiased vs ER+ | -4.3 | 1.8E-5 |
| 2C | Unbiased vs CY+ | -18.0 | 1.0E-72 |
| 2D | Unbiased vs TG+ | -3.7 | 2.4E-4 |
| 2D | Unbiased vs ER+ | -5.3 | 8.5E-8 |
| 2D | Unbiased vs CY+ | -8.6 | 6.1E-18 |
| 2E | Unbiased vs TG+ | -2.3 | 0.024 |
| 2E | Unbiased vs ER+ | -20.5 | 1.5E-93 |
| 2E | Unbiased vs CY+ | -22.3 | 1.5E-110 |
| 2F | Unbiased vs TG+ | -2.3 | 0.020 |
| 2F | Unbiased vs ER+ | -13.1 | 3.9E-39 |
| 2F | Unbiased vs CY+ | -22.9 | 1.2E-115 |
| 2G | Unbiased vs TG+ | -2.7 | 0.007 |
| 2G | Unbiased vs ER+ | -0.193 | 0.85 |
| 2G | Unbiased vs CY+ | -15.2 | 2.6E-52 |
| 2H | Unbiased vs TG+ | -17.4 | 1.7E-67 |
| 2H | Unbiased vs ER+ | -3.7 | 2.5E-4 |
| 2H | Unbiased vs CY+ | -10.0 | 1.9E-23 |
| 3D | No RBP vs TIS11B | -12.3 | 8.7E-35 |
| 3D | No RBP vs LARP4B/METAP2 | -0.481 | 0.630 |
| 3D | No RBP vs TIS11B | -10.2 | 2.4E-24 |
| 3D | No RBP vs LARP4B/METAP2 | -3.5 | 4.5E-4 |
| 3D | No RBP vs TIS11B | -9.1 | 5.9E-20 |
| 3D | No RBP vs LARP4B/METAP2 | -5.2 | 52.5E-7 |
| 3D | No RBP vs TIS11B | -5.9 | 4.2E-9 |
| 3D | No RBP vs LARP4B/METAP2 | -6.2 | 7.0E-10 |
| 3F | No RBP vs TIA1/L1 | -3.9 | 1.1E-4 |
| 3F | No RBP vs LARP4B/METAP2 | -3.4 | 0.001 |
| 3F | No RBP vs TIA1/L1 | -4.1 | 3.7E-5 |
| 3F | No RBP vs LARP4B/METAP2 | -3.1 | 0.002 |
| 3F | No RBP vs TIA1/L1 | -3.3 | 0.001 |
| 3F | No RBP vs LARP4B/METAP2 | -3.3 | 0.001 |
| 3F | No RBP vs TIA1/L1 | -3.4 | 0.001 |
| 3F | No RBP vs LARP4B/METAP2 | -2.2 | 0.027 |
| 3H | No RBP vs LARP4B/METAP2 | -6.3 | 3.2E-10 |
| 3H | No RBP vs TIS11B | -5.7 | 1.4E-8 |
| 3H | No RBP vs LARP4B/METAP2 | -7.3 | 3.8E-13 |
| 3H | No RBP vs TIS11B | -6.9 | 5.9E-12 |
| 3H | No RBP vs LARP4B/METAP2 | -6.1 | 1.5E-9 |
| 3H | No RBP vs TIS11B | -7.8 | 6.1E-15 |
| 3H | No RBP vs LARP4B/METAP2 | -3.1 | 0.002 |
| 3H | No RBP vs TIS11B | -4.5 | 5.6E-6 |
| 4B | No change vs Up in ER | -0.98 | 0.922 |
| 4B | No change vs Up in CY | -2.8 | 0.005 |
| 4B | Up in ER vs Up in CY | -2.4 | 0.018 |
| 4C | No change vs Up in ER | -2.7 | 0.007 |
| 4C | No change vs Up in CY | -3.9 | 1.2E-4 |
| 4C | Up in ER vs Up in CY | -5.2 | 1.9E-7 |
| 4D | No change vs Up in ER | -4.4 | 9.9E-6 |
| 4D | No change vs Up in CY | -1.4 | 0.164 |
| 4D | Up in ER vs Up in CY | -4.9 | 1.1E-6 |
| 5A | No RBP vs TIS11B | -1.0 | 0.296 |
| 5A | No RBP vs TIA1/L1 | -7.9 | 2.4E-15 |
| 5A | TIS11B vs TIA1/L1 | -7.9 | 2.3E-15 |
| 5N | No RBP vs TIA1/L1, unbiased | -3.8 | 1.7E-4 |
| 5N | No RBP vs TIA1/L1, TG+ | -2.0 | 0.044 |
| 5N | No RBP vs TIA1/L1, ER+ | -6.3 | 2.8E-10 |
| 5N | No RBP vs TIA1/L1, CY+ | -2.7 | 0.008 |
| S1J | ER+ vs CY+, ERM | -5.6 | 1.6E-8 |
| S1J | ER+ vs CY+, Cytosol | -4.1 | 3.5E-5 |
| S1M | TG+ vs ER+, Granule proportion in LoRNA | -3.1 | 0.002 |
| S1M | TG+ vs CY+, Granule proportion in LoRNA | -18.4 | 2.7E-75 |
| S1M | ER+ vs CY+, Granule proportion in LoRNA | -14.7 | 5.9E-49 |
| S1M | TG+ vs ER+, Membrane proportion in LoRNA | -20.0 | 1.5E-88 |
| S1M | TG+ vs CY+, Membrane proportion in LoRNA | -20.3 | 1.4E-91 |
| S1M | ER+ vs CY+, Membrane proportion in LoRNA | -31.1 | 5.8E-213 |
| S1M | TG+ vs ER+, Cytosol proportion in LoRNA | -7.4 | 1.4E-13 |
| S1M | TG+ vs CY+, Cytosol proportion in LoRNA | -25.1 | 3.1E-139 |
| S1M | ER+ vs CY+, Cytosol proportion in LoRNA | -27.0 | 3.8E-160 |
| S2H | TG area proportion vs TG foci proportion, *BAG3* | -5.1 | 2.5E-7 |
| S2H | TG area proportion vs TG foci proportion, *DUSP1* | --4.4 | 7.9E-6 |
| S2H | TG area proportion vs TG foci proportion, *DNAJB1* | -3.2 | 0.001 |
| S2H | TG area proportion vs TG foci proportion, *PLA2GA4* | -2.6 | 0.01 |
| S2H | TG area proportion vs TG foci proportion, *IDH1* | -1.7 | 0.088 |
| S2H | TG area proportion vs TG foci proportion, *ALDH18A1* | -0.034 | 0.973 |
| S2H | TG area proportion vs TG foci proportion, *TES* | -2.7 | 0.007 |
| S2H | TG area proportion vs TG foci proportion, *ACTN4* | -1.9 | 0.059 |
| S2I | ER area proportion vs ER foci proportion, *BAG3* | -1.6 | 0.103 |
| S2I | ER area proportion vs ER foci proportion, *DUSP1* | -0.916 | 0.437 |
| S2I | ER area proportion vs ER foci proportion, *DNAJB1* | -0.377 | 0.706 |
| S2I | ER area proportion vs ER foci proportion, *PLA2GA4* | -3.8 | 1.6E-4 |
| S2I | ER area proportion vs ER foci proportion, *IDH1* | -3.3 | 0.001 |
| S2I | ER area proportion vs ER foci proportion, *ALDH18A1* | -2.5 | 0.011 |
| S2I | ER area proportion vs ER foci proportion, *TES* | -2.2 | 0.028 |
| S2I | ER area proportion vs ER foci proportion, *ACTN4* | -1.6 | 0.104 |
| S4A | Unbiased vs lowest TG+ | -9.2 | 2.9E-20 |
| S4A | Unbiased vs lowest ER+ | -1.1 | 0.252 |
| S4A | Unbiased vs lowest CY+ | -6.7 | 2.9E-11 |
| S4A | Lowest TG+ vs lowest ER+ | -6.9 | 3.8E-12 |
| S4A | Lowest TG+ vs lowest CY+ | -11.5 | 9.9E-31 |
| S4A | Lowest ER+ vs lowest CY+ | -3.3 | 0.001 |
| S4B | Unbiased vs lowest TG+ | -6.6 | 3.3E-11 |
| S4B | Unbiased vs lowest ER+ | -4.0 | 5.2E-5 |
| S4B | Unbiased vs lowest CY+ | -2.7 | 0.006 |
| S4B | Lowest TG+ vs lowest ER+ | -7.4 | 9.5E-14 |
| S4B | Lowest TG+ vs lowest CY+ | -7.2 | 4.4E-13 |
| S4B | Lowest ER+ vs lowest CY+ | -1.4 | 0.148 |
| S4C | Unbiased vs TG+ | -20.6 | 5.9E-94 |
| S4C | Unbiased vs ER+ | -2.6 | 0.008 |
| S4C | Unbiased vs CY+ | -13.2 | 1.1E-39 |
| S4D | Unbiased vs lowest TG+ | -5.1 | 4.4E-7 |
| S4D | Unbiased vs lowest ER+ | -3.0 | 0.003 |
| S4D | Unbiased vs lowest CY+ | -8.7 | 3.2E-18 |
| S4D | Lowest TG+ vs lowest ER+ | -1.1 | 0.264 |
| S4D | Lowest TG+ vs lowest CY+ | -9.8 | 6.6E-23 |
| S4D | Lowest ER+ vs lowest CY+ | -7.9 | 2.0E-15 |
| S4E | Unbiased vs lowest TG+ | -0.79 | 0.43 |
| S4E | Unbiased vs lowest ER+ | -3.9 | 9.5E-5 |
| S4E | Unbiased vs lowest CY+ | -4.5 | 7.0E-6 |
| S4E | Lowest TG+ vs lowest ER+ | -3.4 | 0.001 |
| S4E | Lowest TG+ vs lowest CY+ | -2.5 | 0.012 |
| S4E | Lowest ER+ vs lowest CY+ | -6.4 | 1.5E-10 |
| S4F | Unbiased vs lowest TG+ | -0.689 | 0.491 |
| S4F | Unbiased vs lowest ER+ | -11.3 | 1.4E-29 |
| S4F | Unbiased vs lowest CY+ | -10.6 | 1.9E-26 |
| S4F | Lowest TG+ vs lowest ER+ | -9.1 | 1.0E-19 |
| S4F | Lowest TG+ vs lowest CY+ | -8.8 | 1.3E-18 |
| S4F | Lowest ER+ vs lowest CY+ | -15.3 | 2.4E-53 |
| S4G | Unbiased vs lowest TG+ | -1.0 | 0.308 |
| S4G | Unbiased vs lowest ER+ | -6.7 | 1.6E-11 |
| S4G | Unbiased vs lowest CY+ | -10.3 | 4.6E-15 |
| S4G | Lowest TG+ vs lowest ER+ | -4.9 | 1.1E-6 |
| S4G | Lowest TG+ vs lowest CY+ | -8.7 | 5.2E-18 |
| S4G | Lowest ER+ vs lowest CY+ | -11.9 | 1.9E-32 |
| S4H | Unbiased vs lowest TG+ | -2.2 | 0.026 |
| S4H | Unbiased vs lowest ER+ | -0.782 | 0.434 |
| S4H | Unbiased vs lowest CY+ | -6.1 | 1.2E-9 |
| S4H | Lowest TG+ vs lowest ER+ | -0.926 | 0.354 |
| S4H | Lowest TG+ vs lowest CY+ | -6.3 | 2.8E-10 |
| S4H | Lowest ER+ vs lowest CY+ | -4.7 | 2.8E-6 |
| S4I | Unbiased vs lowest TG+ | -7.7 | 1.2E-14 |
| S4I | Unbiased vs lowest ER+ | -0.953 | 0.34 |
| S4I | Unbiased vs lowest CY+ | -4.0 | 6.8E-5 |
| S4I | Lowest TG+ vs lowest ER+ | -5.3 | 1.3E-7 |
| S4I | Lowest TG+ vs lowest CY+ | -8.2 | 1.9E-18 |
| S4I | Lowest ER+ vs lowest CY+ | -3.7 | 2.5E-4 |
| S4J | Unbiased vs TG+ | -10.9 | 1.1E-22 |
| S4J | Unbiased vs ER+ | -18.8 | 8.4E-79 |
| S4J | Unbiased vs CY+ | -14.8 | 1.7E-49 |
| S4K | Unbiased vs lowest TG+ | -4.6 | 3.8E-6 |
| S4K | Unbiased vs lowest ER+ | -11.0 | 5.8E-28 |
| S4K | Unbiased vs lowest CY+ | -7.0 | 3.2E-12 |
| S4K | Lowest TG+ vs lowest ER+ | -11.5 | 7.6E-31 |
| S4K | Lowest TG+ vs lowest CY+ | -1.4 | 0.62 |
| S4K | Lowest ER+ vs lowest CY+ | -13.2 | 5.1E-40 |
