## Supplementary material for "Subcytoplasmic location of translation controls protein output": Table S6

**Table S6. Sequences of oligonucleotides, related to STAR Methods**

| **Sequences of primers for PCR** |  |
| --- | --- |
| THAP1 MS2 F | 5´-ATATTGTACAAGGTGCAGTCCTGCTCCG-3´ |
| THAP1 MS2 R | 5´-CAGTACCGGTTTATGCTGGTACTTCAACTA-3´ |
| MCP F | 5´-CAGTGCTAGCATGGCTTCTAACTTTACTCA-3´ |
| MCP R | 5´-TATAGGATCCGTAGATGCCGGAGTTTGCTG-3´ |
| TIS11B MCP F | 5´-ATATGTACAAGACCACCACCCTCGTGT-3´ |
| TIS11B MCP R | 5´-CAGGTCTAGATTAGTCATCTGAGATGGAAA-3´ |
| TIAL1 MCP F | 5´-TATATGTACAAGATGGAAGACGACGGGCAG-3´ |
| TIAL1 MCP R | 5´-GCGATCTAGATCACTGTGTTTGGTAACTTG-3´ |
| TIS-SEC F | 5´-TAATGCTAGCATGACCACCACCCTCGTGTC-3´ |
| TIS-SEC R | 5´-CAGTGCTAGCGTCATCTGAGATGGAAAGTC-3´ |
| TRAPA MCP F | 5´-TAATGCTAGCATGAGACTCCTCCCCCGCTT-3´ |
| TRAPA MCP R | 5´-CAGTGCTAGCCTCATCAGATCCCACTGATC-3´ |
| TIAL1 CAAX F | 5´-AGGTCCTCCCCAAGGACAAGCTCCTCC-3´ |
| TIAL1 CAAX R | 5´-CCTAGGGCCCTTACATGATAACACACTTGG-3´ |
| CAAX F | 5´-CAGTCTGTACAAGATGGGTGGAGGTTCTGG-3´ |
| **Sequences of sgRNA synthetic DNA** | 5´homology region - 1st gRNA -sgRNA scaffold + Pol III terminator - H1 promoter - 2nd gRNA - 3' homology region |
| *TIS11B* sgRNA  1^st^ sgRNA target: Chr14:68,790,381-68,790,400  2^nd^ sgRNA target:  Chr14:68,790,262-68,790,281 (Hg38) | TGTGGAAAGGACGAAACACCGGGGTGACTGAGTGCCTCCGAGTTTTAGAGCTAGAAATAGCAAGTTAAAATAAGGCTAGTCCGTTATCAACTTGAAAAAGTGGCACCGAGTCGGTGCTTTTTTGAACGCTGACGTCATCAACCCGCTCCAAGGAATCGCGGGCCCAGTGTCACTAGGCGGGAACACCCAGCGCGCGTGCGCCCTGGCAGGAAGATGGCTGTGAGGGACAGGGGAGTGGCGCCCTGCAATATTTGCATGTCGCTATGTGTTCTGGGAAATCACCATAAACGTGAAATGTCTTTGGATTTGGGAATCTTATAAGTTCTGTATGAGACCACTCTTTCCCACCTTCTCGGAAGGGGGCGAGGTTTTAGAGCTAGAAATAGCAAGT |
| Control (intergenic) gRNA  1^st^ sgRNA target:  Chr7:10,195,223-10,195,242  2^nd^ sgRNA target:  Chr7:10,195,281-10,195,300 (Hg38) | TGTGGAAAGGACGAAACACCGTCAACAGGTGAAGATGAACTGTTTTAGAGCTAGAAATAGCAAGTTAAAATAAGGCTAGTCCGTTATCAACTTGAAAAAGTGGCACCGAGTCGGTGCTTTTTTGAACGCTGACGTCATCAACCCGCTCCAAGGAATCGCGGGCCCAGTGTCACTAGGCGGGAACACCCAGCGCGCGTGCGCCCTGGCAGGAAGATGGCTGTGAGGGACAGGGGAGTGGCGCCCTGCAATATTTGCATGTCGCTATGTGTTCTGGGAAATCACCATAAACGTGAAATGTCTTTGGATTTGGGAATCTTATAAGTTCTGTATGAGACCACTCTTTCCCATCAGGGAGACCATTAGTGGGGTTTTAGAGCTAGAAATAGCAAGT |
| **Sequences of RNA-FISH oligos** | **5´- -3´** |
| 2_BAG3 -_Seq_2 | GGGATGTATTGAAGGAGGATGTCGCGGTCACCGTTGCCGGACGCCACCTG |
| 4_BAG3 -_Seq_4 | GGGATGTATTGAAGGAGGATTGTGGTCCACGAAGAAGGGCCAGCCGGTCT |
| 5_BAG3 -_Seq_5 | GGGATGTATTGAAGGAGGATGCGGGTCGTTCCACGTAGTGGTGCGGCTGT |
| 7_BAG3 -_Seq_7 | GGGATGTATTGAAGGAGGATCCTCCCGGGAAGGGCCATTGGCAGAGGAT |
| 8_BAG3 -_Seq_8 | GGGATGTATTGAAGGAGGATGGCCTTCCCTAGCAGGCGGCAGCCTAGAGC |
| 10_BAG3 -_Seq_10 | GGGATGTATTGAAGGAGGATCGCCTTCATGGAGCACAGGAATGGGAATGT |
| 11_BAG3 -_Seq_11 | GGGATGTATTGAAGGAGGATCATGGAAAGGGTGCACCTGCCGGTTCTCAG |
| 13_BAG3 -_Seq_13 | GGGATGTATTGAAGGAGGATTCTGAGGAGCCGCTGCTGCCGCCTCAGTTC |
| 14_BAG3 -_Seq_14 | GGGATGTATTGAAGGAGGATTCTGGCATGCCCCGCAGAGGTGACTGGGAC |
| 16_BAG3 -_Seq_16 | GGGATGTATTGAAGGAGGATGGCTGGGGGCTGGGCTGCCGCCGCCGCTGC |
| 17_BAG3 -_Seq_17 | GGGATGTATTGAAGGAGGATTGGAGACTGGGACCGCTCAGGTCCGTGGGA |
| 19_BAG3 -_Seq_19 | GGGATGTATTGAAGGAGGATGCTCCTGCCGGAGGAAGGCAGGCTGGCCGA |
| 20_BAG3 -_Seq_20 | GGGATGTATTGAAGGAGGATCCCCCGCGGGAGCTGGTGACTGCCCAGGCT |
| 22_BAG3 -_Seq_22 | GGGATGTATTGAAGGAGGATTGGTGGAAGGAGGGCTGGGCTGCTGGCCGG |
| 23_BAG3 -_Seq_23 | GGGATGTATTGAAGGAGGATTGCTGCGCTGGGTAGTGCGTCTTCTGGGCT |
| 25_BAG3 -_Seq_25 | GGGATGTATTGAAGGAGGATGGCTCCCAGTCATCCCCCTGGATCTTGTGG |
| 26_BAG3 -_Seq_26 | GGGATGTATTGAAGGAGGATGACCTGAACGGGGATGCCGCCCGCAGGGGC |
| 28_BAG3 -_Seq_28 | GGGATGTATTGAAGGAGGATTGGAGTGGCGTGCTGCTCCTGGCTGGTGAG |
| 29_BAG3 -_Seq_29 | GGGATGTATTGAAGGAGGATACGGTGTGCACACGGATGGGCGAGGGGGAG |
| 31_BAG3 -_Seq_31 | GGGATGTATTGAAGGAGGATTCAGGCTGGGAAACAGGTGCAGTTTCTCGA |
| 32_BAG3 -_Seq_32 | GGGATGTATTGAAGGAGGATACTGGGCCTGGCTTACTTTCTGGTTTGTTT |
| 34_BAG3 -_Seq_34 | GGGATGTATTGAAGGAGGATGAATCCACCTCTTTGCGGATCACTTGAATT |
| 35_BAG3 -_Seq_35 | GGGATGTATTGAAGGAGGATGGAGGTGGGGGCTTCTGGGAAACAGGTTTA |
| 37_BAG3 -_Seq_37 | GGGATGTATTGAAGGAGGATGGGCTGGGAGGAGGACAAGGAACTGGAGCA |
| 39_BAG3 -_Seq_39 | GGGATGTATTGAAGGAGGATGGGGCTGCCCTCTCTTCTGTAGCCACACTC |
| 40_BAG3 -_Seq_40 | GGGATGTATTGAAGGAGGATGGAGGTGTAGCTTCTGCAGGGGCAGTGCTG |
| 42_BAG3 -_Seq_42 | GGGATGTATTGAAGGAGGATATGGCTTCCACTTTCAGCACTCCTGGATGT |
| 43_BAG3 -_Seq_43 | GGGATGTATTGAAGGAGGATGCCTGCTCCAGCCCCTGTACCTTCTCCAGG |
| 45_BAG3 -_Seq_45 | GGGATGTATTGAAGGAGGATAAATACTCTTCGATCATCAGGTACTTTTTG |
| 46_BAG3 -_Seq_46 | GGGATGTATTGAAGGAGGATACTGAATCCAGGGCCAGCAGCTCTTTGGTC |
| 48_BAG3 -_Seq_48 | GGGATGTATTGAAGGAGGATTGAACCTTCCTGACACCGTCTCTCCTGGCC |
| 49_BAG3 -_Seq_49 | GGGATGTATTGAAGGAGGATGCTTTCTGTTCAAGTTTTTCCAAGATGGTC |
| 51_BAG3 -_Seq_51 | GGGATGTATTGAAGGAGGATTCTGCTTCAAGGTTGCTGGGCTGGAGTTCA |
| 52_BAG3 -_Seq_52 | GGGATGTATTGAAGGAGGATCCCATCTCCATGATTGCCTGCAGTGGCTGA |
| 54_BAG3 -_Seq_54 | GGGATGTATTGAAGGAGGATTCTGTGTGGGGATCTTCTGCATTTCCAGCA |
| 55_BAG3 -_Seq_55 | GGGATGTATTGAAGGAGGATGCTGCTGCTGTGGCTTCTGGCTGCTGGGTT |
| 57_BAG3 -_Seq_57 | GGGATGTATTGAAGGAGGATAGAGGCTACGGTGCTGCTGGGTTACCAGGG |
| 58_BAG3 -_Seq_58 | GGGATGTATTGAAGGAGGATCATCGGTTCCGAGTCTGATTTTTACAGGGC |
| 60_BAG3 -_Seq_60 | GGGATGTATTGAAGGAGGATTAAAAACCAACTGACTTAAAGTCTCTGAAA |
| 61_BAG3 -_Seq_61 | GGGATGTATTGAAGGAGGATCACCCAAGTTACTGCATACCAAGCAGCTAA |
| 63_BAG3 -_Seq_63 | GGGATGTATTGAAGGAGGATCAGAGTAAGAATATAGAAGAAAAGCATCAT |
| 64_BAG3 -_Seq_64 | GGGATGTATTGAAGGAGGATCTCAAACAACAAGCAACTTCTTTATTTGTA |
| 66_BAG3 -_Seq_66 | GGGATGTATTGAAGGAGGATAGGTGGTGGGGGTGCCCAAGTAGACAGGGC |
| 67_BAG3 -_Seq_67 | GGGATGTATTGAAGGAGGATTACAAAAGACAGTGCACAACCACAGCTAAC |
| 69_BAG3 -_Seq_69 | GGGATGTATTGAAGGAGGATCTGATAAATGTTTCATATTTATGTGATGGG |
| 70_BAG3 -_Seq_70 | GGGATGTATTGAAGGAGGATGAAGAAAATCATCTCATTAAAATGGCAACA |
| 72_BAG3 -_Seq_72 | GGGATGTATTGAAGGAGGATATATTCCTATGGCTCCTGGCACATTTTACT |
| 73_BAG3 -_Seq_73 | GGGATGTATTGAAGGAGGATAATGTAGCATTAAAGTCATCCAACATACAG |
| 2_DUSP1 -_Seq_2 | GGGATGTATTGAAGGAGGATTCGCTCCCCCAGCAGCGCCCGCAGGCCTCC |
| 3_DUSP1 -_Seq_3 | GGGATGTATTGAAGGAGGATGCGGCAGTCCAGCAGCAGGCATTGCGCCGC |
| 4_DUSP1 -_Seq_4 | GGGATGTATTGAAGGAGGATGATGTGGCCGGCGTTGAAAGCGAAGAAGGA |
| 5_DUSP1 -_Seq_5 | GGGATGTATTGAAGGAGGATGGTGCTGAAGCGCACGTTGACAGAGCCGGC |
| 6_DUSP1 -_Seq_6 | GGGATGTATTGAAGGAGGATCTCCAGGCCCATGGCGCCCTTGGCCCGGCG |
| 7_DUSP1 -_Seq_7 | GGGATGTATTGAAGGAGGATCAGCAACACCACGGCGTGGTAGGCGCCGGC |
| 8_DUSP1 -_Seq_8 | GGGATGTATTGAAGGAGGATCCAGGGTGCCGTCGCGCTTGGCGCCGTCCA |
| 9_DUSP1 -_Seq_9 | GGGATGTATTGAAGGAGGATTTGAGGAAGAAGACTTGCGCGGCGCGCGCC |
| 10_DUSP1 -_Seq_10 | GGGATGTATTGAAGGAGGATCAGGAAGCCGAAAACGCTTCGTATCCTCCT |
| 11_DUSP1 -_Seq_11 | GGGATGTATTGAAGGAGGATGGGGGTCGACTGTTTGCTGCACAGCTCCGG |
| 12_DUSP1 -_Seq_12 | GGGATGTATTGAAGGAGGATGCTAGTACTCAGGGGAAGGCTGAGCCCCAT |
| 13_DUSP1 -_Seq_13 | GGGATGTATTGAAGGAGGATACTGCACCCAGATTCCGCGCTGTCAGGGAC |
| 15_DUSP1 -_Seq_15 | GGGATGTATTGAAGGAGGATGTACAGAAAGGGCAGGATTTCCACCGGGCC |
| 16_DUSP1 -_Seq_16 | GGGATGTATTGAAGGAGGATCTTGCGGGAAGCGTGATACGCACTGCCCAG |
| 17_DUSP1 -_Seq_17 | GGGATGTATTGAAGGAGGATGGCAGTGATGCCCAAGGCATCCAGCATGTC |
| 18_DUSP1 -_Seq_18 | GGGATGTATTGAAGGAGGATGTTGGGACAATTGGCTGAGACGTTGATCAA |
| 19_DUSP1 -_Seq_19 | GGGATGTATTGAAGGAGGATGCTCTTGTACTGGTAGTGACCCTCAAAATG |
| 20_DUSP1 -_Seq_20 | GGGATGTATTGAAGGAGGATGTCTGCCTTGTGGTTGTCCTCCACAGGGAT |
| 21_DUSP1 -_Seq_21 | GGGATGTATTGAAGGAGGATGTCAATGGCCTCGTTGAACCAGGAGCTGAT |
| 22_DUSP1 -_Seq_22 | GGGATGTATTGAAGGAGGATTCCTCCAGCATTCTTGATGGAGTCTATGAA |
| 23_DUSP1 -_Seq_23 | GGGATGTATTGAAGGAGGATAATGCCTGCCTGGCAGTGGACAAACACCCT |
| 24_DUSP1 -_Seq_24 | GGGATGTATTGAAGGAGGATGTAAGCAAGGCAGATGGTGGCTGACCGGGA |
| 25_DUSP1 -_Seq_25 | GGGATGTATTGAAGGAGGATGTCCAGCTTGACTCGATTAGTCCTCATAAG |
| 26_DUSP1 -_Seq_26 | GGGATGTATTGAAGGAGGATTCGCCTCTGCTTCACAAACTCAAAGGCCTC |
| 28_DUSP1 -_Seq_28 | GGGATGTATTGAAGGAGGATCACCTGGGACTCAAACTGCAGCAGCTGGCC |
| 29_DUSP1 -_Seq_29 | GGGATGTATTGAAGGAGGATCCCAGCCTCTGCCGAACAGTGCGGAGCCAG |
| 30_DUSP1 -_Seq_30 | GGGATGTATTGAAGGAGGATGAGGTGCCTCGGTCGAGCACAGCCATGGCG |
| 31_DUSP1 -_Seq_31 | GGGATGTATTGAAGGAGGATGAGACGGGGAAGTTGAACACGGTGGTGGTG |
| 32_DUSP1 -_Seq_32 | GGGATGTATTGAAGGAGGATAGCGCACTGTTCGTGGAGTGGACAGGGATG |
| 33_DUSP1 -_Seq_33 | GGGATGTATTGAAGGAGGATGAGGTCGTAATGGGGCTCTGAAGGTAGCTC |
| 34_DUSP1 -_Seq_34 | GGGATGTATTGAAGGAGGATTCACCTCCCGTGGCCTTTCAGCAGCTGGGA |
| 35_DUSP1 -_Seq_35 | GGGATGTATTGAAGGAGGATGCATGGAGTCCCAATGGGATGTGAAGAGCC |
| 36_DUSP1 -_Seq_36 | GGGATGTATTGAAGGAGGATCCAGAGTTATTGCATTTCTCCTCTCAAGGA |
| 37_DUSP1 -_Seq_37 | GGGATGTATTGAAGGAGGATTAAATAAGGACCAGCCCTCTCGAGCCCCTC |
| 38_DUSP1 -_Seq_38 | GGGATGTATTGAAGGAGGATGAAACCCAGAGGAACTCGGGTGAAGTTAAA |
| 39_DUSP1 -_Seq_39 | GGGATGTATTGAAGGAGGATTTGACGCTAAGTCATCACCATAACTGCTTA |
| 41_DUSP1 -_Seq_41 | GGGATGTATTGAAGGAGGATAGATGGACTTGATGTACCCACTATATATTG |
| 42_DUSP1 -_Seq_42 | GGGATGTATTGAAGGAGGATTGAGTCCTTTCTCTTCTGCCCCATTTTGTC |
| 43_DUSP1 -_Seq_43 | GGGATGTATTGAAGGAGGATGGGCGAGCAAAAAGAAACCGGATCACACAC |
| 44_DUSP1 -_Seq_44 | GGGATGTATTGAAGGAGGATTATGTCAAGCATGAAGAGATTCTACAAAAA |
| 45_DUSP1 -_Seq_45 | GGGATGTATTGAAGGAGGATATATGTGTCGTCGGGAATAATACTGGTAGG |
| 46_DUSP1 -_Seq_46 | GGGATGTATTGAAGGAGGATCAGACACCTACACAAAAATAAATAAGGTAT |
| 47_DUSP1 -_Seq_47 | GGGATGTATTGAAGGAGGATTCTAGGAGTAGACAATGACATTTGTGAAGG |
| 48_DUSP1 -_Seq_48 | GGGATGTATTGAAGGAGGATCTCAAAAACAAAAATTGAGGTATTTGGTTC |
| 49_DUSP1 -_Seq_49 | GGGATGTATTGAAGGAGGATGCTTAAGATATATTTACAGGATAGTACAGT |
| 50_DUSP1 -_Seq_50 | GGGATGTATTGAAGGAGGATGTATTTTCCATCAGTGCTGAAAACAAACCT |
| 51_DUSP1 -_Seq_51 | GGGATGTATTGAAGGAGGATAGCAAACATACAACTGTTGGCAACTAAAAA |
| 52_DUSP1 -_Seq_52 | GGGATGTATTGAAGGAGGATCCTTATGTAACAAAATGTCTTCTTAGAAGA |
| 1_SF3A2 -_Seq_1 | GGGATGTATTGAAGGAGGATCTTGCCCCCGGGGCGATGCTGGAAGTCCAT |
| 2_SF3A2 -_Seq_2 | GGGATGTATTGAAGGAGGATGGAGGAGGAGGCCACGCCCCCGCTCCCGGT |
| 3_SF3A2 -_Seq_3 | GGGATGTATTGAAGGAGGATGAGGCGCTCCCTGCGGTCACGGTTGCTCTC |
| 4_SF3A2 -_Seq_5 | GGGATGTATTGAAGGAGGATGTGGTTCTTCATGAAGTACGGGTCCTTGTT |
| 5_SF3A2 -_Seq_6 | GGGATGTATTGAAGGAGGATCAGGCAGAGTTTGCATTCATAGGAGCCCAG |
| 6_SF3A2 -_Seq_7 | GGGATGTATTGAAGGAGGATCAGGTAGCTCCCCTCATTGTTGTGAAGTGT |
| 7_SF3A2 -_Seq_8 | GGGATGTATTGAAGGAGGATGGTCTGGTGCTTCTTCCCCTGCGTATGTGC |
| 8_SF3A2 -_Seq_9 | GGGATGTATTGAAGGAGGATGGCCTCCTTGGCTGCTCGCCGGGCCAGGTT |
| 9_SF3A2 -_Seq_10 | GGGATGTATTGAAGGAGGATTGACCTTCTCAGGCGCGGGCTGGGCAGGGG |
| 10_SF3A2 -_Seq_11 | GGGATGTATTGAAGGAGGATCGATCTTCACAAACTTCTTCACCTCCACCT |
| 11_SF3A2 -_Seq_12 | GGGATGTATTGAAGGAGGATTCTGCTTGGTCACTTTGTAGCCCGGGCGGC |
| 12_SF3A2 -_Seq_13 | GGGATGTATTGAAGGAGGATGGAGGCTCTGCTGGCCCATCTCCGAGTCTC |
| 13_SF3A2 -_Seq_14 | GGGATGTATTGAAGGAGGATCGGCGATCTCAGGGTAGTCAATCTGGAAGA |
| 14_SF3A2 -_Seq_15 | GGGATGTATTGAAGGAGGATACATGAAGCGGTGACGTGGCATGATGCCCT |
| 15_SF3A2 -_Seq_16 | GGGATGTATTGAAGGAGGATCCGGAGGCTCGATCCTCTGCTCGTACGCAG |
| 16_SF3A2 -_Seq_17 | GGGATGTATTGAAGGAGGATTCGGCGGCCATGAGCAGGTACTGCCAGCGC |
| 17_SF3A2 -_Seq_18 | GGGATGTATTGAAGGAGGATGGCACCTTGAAGGCAATGGTCTCGTAGGGC |
| 18_SF3A2 -_Seq_19 | GGGATGTATTGAAGGAGGATTTGCCCTCCGCCTTGTCGATCTCTCTGCTC |
| 19_SF3A2 -_Seq_20 | GGGATGTATTGAAGGAGGATTTGGTCTCCCGGTTCCAGTGTGTCCAGAAC |
| 20_SF3A2 -_Seq_22 | GGGATGTATTGAAGGAGGATGGGAGGCTGGGTGGAGCCGGGGGCTTCTCC |
| 21_SF3A2 -_Seq_23 | GGGATGTATTGAAGGAGGATCGGGGGTGGAGGCCGCTTCACCCCAGGGGG |
| 22_SF3A2 -_Seq_24 | GGGATGTATTGAAGGAGGATCGGTGGCCGAGGGGGCAGACCGTTCATCAG |
| 23_SF3A2 -_Seq_25 | GGGATGTATTGAAGGAGGATTGGCGGGGGCGGTGGCAAAGACTCAGGCAG |
| 24_SF3A2 -_Seq_26 | GGGATGTATTGAAGGAGGATTAGCTGGGGTGGTCCCGGGGGCCCTGAGGG |
| 25_SF3A2 -_Seq_27 | GGGATGTATTGAAGGAGGATTGCAGGGGGATGCACCACTGGGGCCGGGGG |
| 26_SF3A2 -_Seq_28 | GGGATGTATTGAAGGAGGATGACGCCAGGAGCTGGGGGATGGACCCCAGA |
| 27_SF3A2 -_Seq_29 | GGGATGTATTGAAGGAGGATGGCTGGGGGATGGACGCCAGGAGCTGGGGG |
| 28_SF3A2 -_Seq_30 | GGGATGTATTGAAGGAGGATGACCCCAGAGGTTGGTGGGTGGACCCCAGG |
| 29_SF3A2 -_Seq_31 | GGGATGTATTGAAGGAGGATGGCTGGAGGGTGGACTCCAGGAGCTGGGGG |
| 30_SF3A2 -_Seq_32 | GGGATGTATTGAAGGAGGATGACTCCGGGGGCTGGTGGGTGAACCCCGGG |
| 31_SF3A2 -_Seq_33 | GGGATGTATTGAAGGAGGATTGGTGGGTGAACCCCAGGGGCTGGTGGGTG |
| 32_SF3A2 -_Seq_34 | GGGATGTATTGAAGGAGGATCGCTGATGGGGGAGGATGGACCCCTGGGGC |
| 33_SF3A2 -_Seq_35 | GGGATGTATTGAAGGAGGATTGGGTGCACCCCCGGGGCCTGGGGGTGAAC |
| 34_SF3A2 -_Seq_36 | GGGATGTATTGAAGGAGGATTGGGGCCTGAGGGTGAACGGCGGGGGCTGC |
| 35_SF3A2 -_Seq_37 | GGGATGTATTGAAGGAGGATAGGGTGCATCCCTGGGGCTGGTGGGTGCAC |
| 36_SF3A2 -_Seq_38 | GGGATGTATTGAAGGAGGATGGGAGGTTGGGGGTGGACCCCCGGGGCCTG |
| 37_SF3A2 -_Seq_39 | GGGATGTATTGAAGGAGGATGGTGGACCCCAGGAGCCGACGGATGGACCC |
| 38_SF3A2 -_Seq_40 | GGGATGTATTGAAGGAGGATGATTTGAGGGGTGAACTCCCGGAGGCTGAG |
| 39_SF3A2 -_Seq_41 | GGGATGTATTGAAGGAGGATCCTCAGCATTGGGGGCATGGGAGTTGGGGG |
| 40_SF3A2 -_Seq_42 | GGGATGTATTGAAGGAGGATGTTCCCTGGGCCTTCGGAGGGAAGTGGGGG |
| 41_SF3A2 -_Seq_43 | GGGATGTATTGAAGGAGGATTTCTCAGTTGGTTGGGGGAGGGGGAGGTAT |
| 42_SF3A2 -_Seq_44 | GGGATGTATTGAAGGAGGATTGGCGCTGGGCTTGCTGGGGGAGGGAGCAG |
| 43_SF3A2 -_Seq_45 | GGGATGTATTGAAGGAGGATCTTCTCTCAGTGGGAAAAGGCAAGAGCACC |
| 1_MLST8 ._Seq_1 | GGGATGTATTGAAGGAGGATACTGCCCACCGTGCCTGGGGAGGTGTTCAT |
| 2_MLST8 ._Seq_2 | GGGATGTATTGAAGGAGGATGTAGCCTGCAGTGGCCAGGATGACCGGGTC |
| 3_MLST8 ._Seq_3 | GGGATGTATTGAAGGAGGATGTGGGCCTGCCAGAAGCGCACGGTGTGGTC |
| 4_MLST8 ._Seq_4 | GGGATGTATTGAAGGAGGATGTGCTGCACCGTCCGGGTGCAGATGCCGCT |
| 5_MLST8 ._Seq_5 | GGGATGTATTGAAGGAGGATGACCTCCAAGGCATTCACCTGGGAGTCCTG |
| 6_MLST8 ._Seq_6 | GGGATGTATTGAAGGAGGATTGCAGCAGCAATCATGCTGCGGTCCGGTGT |
| 7_MLST8 ._Seq_7 | GGGATGTATTGAAGGAGGATGAGATCATACATGCGGATGTGCTGGTAACC |
| 8_MLST8 ._Seq_8 | GGGATGTATTGAAGGAGGATGCTGATGATGGGGTTAGGGTTATTGGAGTT |
| 9_MLST8 ._Seq_9 | GGGATGTATTGAAGGAGGATAGACGCGATGTTCTTGTTGACGCCGTCGTA |
| 10_MLST8 ._Seq_10 | GGGATGTATTGAAGGAGGATCATCCAGCGGCCGTCTTCGTGGAAGCCCAC |
| 11_MLST8 ._Seq_11 | GGGATGTATTGAAGGAGGATCCTGGCTGTGCAGTCCTCGCCGCCCGTGTA |
| 12_MLST8 ._Seq_12 | GGGATGTATTGAAGGAGGATCTGCAGGTTCCGGGACCTGAGGTCCCAGAT |
| 13_MLST8 ._Seq_13 | GGGATGTATTGAAGGAGGATGGGTGCGTTCACCTGGAAGATCCGCTGGCA |
| 14_MLST8 ._Seq_14 | GGGATGTATTGAAGGAGGATCTGGTTGGGGTGCAGGCACACGCAGTTAAT |
| 15_MLST8 ._Seq_15 | GGGATGTATTGAAGGAGGATCCCGCTCTGGTCACCCACGATGAGCTCTGC |
| 16_MLST8 ._Seq_16 | GGGATGTATTGAAGGAGGATGTCTGTTTTCAAGTCCCAGATGTGGATAGC |
| 17_MLST8 ._Seq_17 | GGGATGTATTGAAGGAGGATCTCGGGCTCAGGGATCAGCTGCTCGTTGTG |
| 18_MLST8 ._Seq_18 | GGGATGTATTGAAGGAGGATGGGATCGATGTGGGCGGACGTGATGGAGAC |
| 19_MLST8 ._Seq_19 | GGGATGTATTGAAGGAGGATGCTATTGACAGCTGCCATGTAGCTGGCGTC |
| 20_MLST8 ._Seq_20 | GGGATGTATTGAAGGAGGATCGTCAGATTCCAGACATAGCAGTTTCCGGT |
| 21_MLST8 ._Seq_21 | GGGATGTATTGAAGGAGGATGAGCTGGGTCACCTCGTCACCAATGCCCCC |
| 22_MLST8 ._Seq_22 | GGGATGTATTGAAGGAGGATCGTGTGGGCAGGGATCTTAGTCTTGGGGAT |
| 23_MLST8 ._Seq_23 | GGGATGTATTGAAGGAGGATGGGGCTGAAGCGACACTGCAGGGCGTAGCG |
| 24_MLST8 ._Seq_24 | GGGATGTATTGAAGGAGGATAGCCGAGCAGGTGGCGAGGAGCGTGGAGTC |
| 25_MLST8 ._Seq_25 | GGGATGTATTGAAGGAGGATGGACGTCCTCCAGATCTTGCACGTCTGATC |
| 26_MLST8 ._Seq_26 | GGGATGTATTGAAGGAGGATGATGCTCAGCTCCGTCATCAGGGAGAAGTT |
| 27_MLST8 ._Seq_27 | GGGATGTATTGAAGGAGGATGCGGGAGGACTCCCCGGGGTTGCCGCTCTT |
| 28_MLST8 ._Seq_28 | GGGATGTATTGAAGGAGGATCCCCGAGAAGGCGCAGCCCCACATCCAGCC |
| 29_MLST8 ._Seq_29 | GGGATGTATTGAAGGAGGATCGAGGAAGCAGTGACGATGTACTGGGAGTC |
| 30_MLST8 ._Seq_30 | GGGATGTATTGAAGGAGGATCTCCACACACCAGAGCCGGGCCAGGTTGTC |
| 31_MLST8 ._Seq_31 | GGGATGTATTGAAGGAGGATGCCGCCATACTCTCTCTTGATCTCTCCAGT |
| 32_MLST8 ._Seq_32 | GGGATGTATTGAAGGAGGATGAAGGCCAGGCAGACAACAGCCTTCTGGTG |
| 33_MLST8 ._Seq_33 | GGGATGTATTGAAGGAGGATGGTCACAGGCTAGCCCAGCACACTGTCATT |
| 34_MLST8 ._Seq_34 | GGGATGTATTGAAGGAGGATTGCCACCACCTGCACCAGGCAGTCCCGAGG |
| 35_MLST8 ._Seq_35 | GGGATGTATTGAAGGAGGATTGACCTGGGTGCTGCATGGGTCCCTCCAGC |
| 36_MLST8 ._Seq_36 | GGGATGTATTGAAGGAGGATTGGCGCAGGCCGGCAGGGGAGGGTCTGCTC |
| 37_MLST8 ._Seq_37 | GGGATGTATTGAAGGAGGATAAGGCGCCACAGGGGGCCATCAGGTCCAGC |
| 38_MLST8 ._Seq_38 | GGGATGTATTGAAGGAGGATGGCTGAGAGTCCCAGGGCAGCCTGGCCCAG |
| 39_MLST8 ._Seq_39 | GGGATGTATTGAAGGAGGATAGCTCTGTCACATCTGGATAAGCAACTGGG |
| 40_MLST8 ._Seq_40 | GGGATGTATTGAAGGAGGATGTCCAGGAGTGTGCAGCCTGGCTTGGGTCG |
| 41_MLST8 ._Seq_41 | GGGATGTATTGAAGGAGGATCGACTTTCCCAGGCAGTGCAGGCTAGCCCA |
| 42_MLST8 ._Seq_42 | GGGATGTATTGAAGGAGGATCAGACCCCTCAGCAGCTTTGGGCCCTCGGC |
| 43_MLST8 ._Seq_43 | GGGATGTATTGAAGGAGGATACACACTAGCTTGGGGGTGGGCACCAGCCT |
| 44_MLST8 ._Seq_44 | GGGATGTATTGAAGGAGGATCCCTGAAACGCGGGCAGGGAGGGGCAGAGA |
| 45_MLST8 ._Seq_45 | GGGATGTATTGAAGGAGGATCATGGTGGTGGTGTTCTCTATGGACCGAGG |
| 1_MAP2K2 -_Seq_1 | GGGATGTATTGAAGGAGGATCGGCAGCACCGGCTTCCTCCGGGCCAGCAT |
| 2_MAP2K2 -_Seq_3 | GGGATGTATTGAAGGAGGATGGAGGCGCCCTCGCTGGTAGGGGATGGGCC |
| 3_MAP2K2 -_Seq_8 | GGGATGTATTGAAGGAGGATCGCGCCCAGCTCTGAGATCCTTTCGAAGTC |
| 4_MAP2K2 -_Seq_12 | GGGATGTATTGAAGGAGGATCTGCAGCTCGCGGATGATCTGGTTCCGGAT |
| 5_MAP2K2 -_Seq_13 | GGGATGTATTGAAGGAGGATGATGTACGGCGAGTTGCATTCGTGCAGGAC |
| 6_MAP2K2 -_Seq_14 | GGGATGTATTGAAGGAGGATGTCACTGTAGAAGGCCCCGTAGAAGCCCAC |
| 7_MAP2K2 -_Seq_15 | GGGATGTATTGAAGGAGGATCATGTGTTCCATGCAAATGCTGATCTCCCC |
| 8_MAP2K2 -_Seq_16 | GGGATGTATTGAAGGAGGATTTTCAGCACCTGGTCCAGGGAGCCGCCGTC |
| 9_MAP2K2 -_Seq_17 | GGGATGTATTGAAGGAGGATCAGGATCTCCTCGGGAATCCTCTTGGCCTC |
| 10_MAP2K2 -_Seq_18 | GGGATGTATTGAAGGAGGATGCCCCGGAGAACCGCGATGCTGACTTTCCC |
| 11_MAP2K2 -_Seq_20 | GGGATGTATTGAAGGAGGATGATGTTGGAGGGCTTCACATCTCGGTGCAT |
| 12_MAP2K2 -_Seq_25 | GGGATGTATTGAAGGAGGATGATGTCCGACTGCACCGAGTAATGTGTGCC |
| 13_MAP2K2 -_Seq_26 | GGGATGTATTGAAGGAGGATCAGCTCCACCAGGGACAGGCCCATGCTCCA |
| 14_MAP2K2 -_Seq_27 | GGGATGTATTGAAGGAGGATGGCCTCCAGCTCTTTGGCGTCGGGCGGGGG |
| 15_MAP2K2 -_Seq_28 | GGGATGTATTGAAGGAGGATTTCCCCGTCGACCACGGGCCGGCCAAAGAT |
| 16_MAP2K2 -_Seq_29 | GGGATGTATTGAAGGAGGATCCGAGGCGAGATGCTGTGAGGCTCTCCTTC |
| 17_MAP2K2 -_Seq_30 | GGGATGTATTGAAGGAGGATCCATCCCGTGACCGCTGACGGGGCGCCCGG |
| 18_MAP2K2 -_Seq_31 | GGGATGTATTGAAGGAGGATGTTCAAAGATGGCCATGGCAGGCCGGCTAT |
| 19_MAP2K2 -_Seq_32 | GGGATGTATTGAAGGAGGATGAGGTGGCTCGTTCACAATATAGTCCAGGA |
| 20_MAP2K2 -_Seq_35 | GGGATGTATTGAAGGAGGATATCTTCAGGTCCGCCCGCTCCGCTGGGTTC |
| 21_MAP2K2 -_Seq_36 | GGGATGTATTGAAGGAGGATGACCGCTTGATGAAGGTGTGGTTTGTGAGC |
| 22_MAP2K2 -_Seq_37 | GGGATGTATTGAAGGAGGATCAGCCGGCAAAATCCACTTCTTCCACCTCG |
| 23_MAP2K2 -_Seq_38 | GGGATGTATTGAAGGAGGATGGCTGGTTCAGCCGCAGGGTTTTACACAAC |
| 24_MAP2K2 -_Seq_39 | GGGATGTATTGAAGGAGGATCACTGTCACACGGCGGTGCGCGTGGGTGTG |
| 25_MAP2K2 -_Seq_40 | GGGATGTATTGAAGGAGGATGACGGTGGGCAGGTCACCAGCGGGACGCAG |
| 26_MAP2K2 -_Seq_41 | GGGATGTATTGAAGGAGGATCCTCAGCTGGAAGGGCGGGGCATGGACAGG |
| 27_MAP2K2 -_Seq_42 | GGGATGTATTGAAGGAGGATGGTGAGGCAGGAGGGTGGGTGGAGGCGCCA |
| 28_MAP2K2 -_Seq_43 | GGGATGTATTGAAGGAGGATCATGCGCTGTCGCCCCGCCACGGTGCTCTC |
| 29_MAP2K2 -_Seq_44 | GGGATGTATTGAAGGAGGATGACGGGCAGGAGAGGAGACCCCCGTTCCTG |
| 30_MAP2K2 -_Seq_45 | GGGATGTATTGAAGGAGGATGTCGCCCGTCCCCAGAGGCACCCCGGCCAG |
| 31_MAP2K2 -_Seq_46 | GGGATGTATTGAAGGAGGATAGCAGAGCCTCTGAGACCACACACAGCAGC |
| 32_MAP2K2 -_Seq_47 | GGGATGTATTGAAGGAGGATCTCTCCCTGTTTTGTTTTGTAACCTAAGGA |
| 3_ALDH18A1 -_Seq_3 | GGGATGTATTGAAGGAGGATATGAGATCTGAAGACGGTTGTACACTTGAC |
| 5_ALDH18A1 -_Seq_5 | GGGATGTATTGAAGGAGGATAGTGATAAACGGGATGTTGCTCCAAGAACG |
| 7_ALDH18A1 -_Seq_7 | GGGATGTATTGAAGGAGGATGGCATGCTTCAGCTCACTGCGGTGGGCGAA |
| 9_ALDH18A1 -_Seq_9 | GGGATGTATTGAAGGAGGATCAGGCCACATTCATCCCCTCGGGTCACCAC |
| 11_ALDH18A1 -_Seq_11 | GGGATGTATTGAAGGAGGATTCTGCCCTGATTCTGCAGCACTGATACCTG |
| 13_ALDH18A1 -_Seq_13 | GGGATGTATTGAAGGAGGATCTCATGGCGCAAGCGTTGTTTGCCAAAGGC |
| 15_ALDH18A1 -_Seq_15 | GGGATGTATTGAAGGAGGATTTCTTTCAGCTGGTTCTGCCCCGAGTGGAG |
| 17_ALDH18A1 -_Seq_17 | GGGATGTATTGAAGGAGGATCATCAGCCCACTCTGTCCGGCAGCTGCACA |
| 20_ALDH18A1 -_Seq_20 | GGGATGTATTGAAGGAGGATGCGCTTCTGCTCATCATGGAAATCCAAATT |
| 22_ALDH18A1 -_Seq_22 | GGGATGTATTGAAGGAGGATGTTGACAATGGGGACAATGTTCATTCTAAG |
| 24_ALDH18A1 -_Seq_24 | GGGATGTATTGAAGGAGGATGCTATCATTATCTTTAACACTAATAACCCC |
| 26_ALDH18A1 -_Seq_26 | GGGATGTATTGAAGGAGGATTACATCTGAAAGAACAATCAAGAGATCAGT |
| 28_ALDH18A1 -_Seq_28 | GGGATGTATTGAAGGAGGATAGACTGCTGATCTCCGGGATAAAATATATC |
| 30_ALDH18A1 -_Seq_30 | GGGATGTATTGAAGGAGGATTGCTTTCACCTTGGCTTCCATGCCACCCAT |
| 32_ALDH18A1 -_Seq_32 | GGGATGTATTGAAGGAGGATCTTTGGGTGGGTTCCATTGGCAATAACAAC |
| 34_ALDH18A1 -_Seq_34 | GGGATGTATTGAAGGAGGATTGAAAAGAAGGTACCAACTTTCTTCCCCTC |
| 37_ALDH18A1 -_Seq_37 | GGGATGTATTGAAGGAGGATCTGCTCAGGTTCCAAGGTGGCCAACATCCT |
| 39_ALDH18A1 -_Seq_39 | GGGATGTATTGAAGGAGGATCAGGATCTCATCACGCTGGTCCGTCAACAG |
| 41_ALDH18A1 -_Seq_41 | GGGATGTATTGAAGGAGGATGGCTTAAACGTTTCAGCAGAGGAGCTGCAA |
| 43_ALDH18A1 -_Seq_43 | GGGATGTATTGAAGGAGGATGGGAGGAGGCTGCGATCTGTCGCAGACCGA |
| 45_ALDH18A1 -_Seq_45 | GGGATGTATTGAAGGAGGATGTTCCAGTTCCAAGTTTTTGGCGATTCGGG |
| 47_ALDH18A1 -_Seq_47 | GGGATGTATTGAAGGAGGATGTAGACAGTCAGGACGAGATTCAAAGATCA |
| 49_ALDH18A1 -_Seq_49 | GGGATGTATTGAAGGAGGATCCTTCCCTCCTTTGAGTAACAAGCCATTGC |
| 51_ALDH18A1 -_Seq_51 | GGGATGTATTGAAGGAGGATCATGGATTGAGAGAGCCTCCTGGGTCAGGA |
| 54_ALDH18A1 -_Seq_54 | GGGATGTATTGAAGGAGGATGTGGAATGATCAGATCTATCATTTTGTCTA |
| 56_ALDH18A1 -_Seq_56 | GGGATGTATTGAAGGAGGATCCATCACTGGAATCCCCTTAGCAGCTTTCT |
| 58_ALDH18A1 -_Seq_58 | GGGATGTATTGAAGGAGGATTGACCTTATCAACACTGGCCTCGGAATCCA |
| 60_ALDH18A1 -_Seq_60 | GGGATGTATTGAAGGAGGATAAGTCTCCAAAGCATTACAGGCAGCTGGAT |
| 62_ALDH18A1 -_Seq_62 | GGGATGTATTGAAGGAGGATTCAGCATATCAATGATCTGGTCAAATAATG |
| 64_ALDH18A1 -_Seq_64 | GGGATGTATTGAAGGAGGATGGCTGAAGGTCAGATAGGAGGCAAATTTGG |
| 66_ALDH18A1 -_Seq_66 | GGGATGTATTGAAGGAGGATCTACTTCAATGCATAATTCCAGGTCCCCAT |
| 68_ALDH18A1 -_Seq_68 | GGGATGTATTGAAGGAGGATCATCCGTGTGGGAGCTGCCATACTTGTGGA |
| 71_ALDH18A1 -_Seq_71 | GGGATGTATTGAAGGAGGATAAAAGCGAGTGCTGGCATTCCAGAACACAC |
| 73_ALDH18A1 -_Seq_73 | GGGATGTATTGAAGGAGGATCGTGGATTCTCGATGTACTGATTCCCACTT |
| 75_ALDH18A1 -_Seq_75 | GGGATGTATTGAAGGAGGATCCTTCCCTCGCAGCAGCCACTTAGTAGTAA |
| 77_ALDH18A1 -_Seq_77 | GGGATGTATTGAAGGAGGATGGAGGTTCTCATGAAGATATTTTAAACTTC |
| 79_ALDH18A1 -_Seq_79 | GGGATGTATTGAAGGAGGATCTTTTGGAAAATTCCCGGGTTTTCCTGGCT |
| 81_ALDH18A1 -_Seq_81 | GGGATGTATTGAAGGAGGATCAGGAACTGGGAGACAAGAGCGGGCTCTCT |
| 83_ALDH18A1 -_Seq_83 | GGGATGTATTGAAGGAGGATTGCTATTGCCAAACGGAGCCCAGAAGCATC |
| 85_ALDH18A1 -_Seq_85 | GGGATGTATTGAAGGAGGATCTATCGGTAGCAACTATTTTCTTACTTTAA |
| 88_ALDH18A1 -_Seq_88 | GGGATGTATTGAAGGAGGATAAGGGGGACTGTAAGTCACTGAGGTGACAC |
| 90_ALDH18A1 -_Seq_90 | GGGATGTATTGAAGGAGGATGATGACTACATGTGAAAGAAATAAAATGTG |
| 92_ALDH18A1 -_Seq_92 | GGGATGTATTGAAGGAGGATTCTGATGAGAATGCTAAAAAGAAGAGTTGC |
| 94_ALDH18A1 -_Seq_94 | GGGATGTATTGAAGGAGGATCAAATAAATACAAAAGGTCAATCTTCCCAG |
| 96_ALDH18A1 -_Seq_96 | GGGATGTATTGAAGGAGGATCACAGCATCTCCTGGATGCAGGAAGCTGCA |
| 98_ALDH18A1 -_Seq_98 | GGGATGTATTGAAGGAGGATGGAACATCCTGAAACTTGCATCTCCTGCTG |
| 100_ALDH18A1 -_Seq_100 | GGGATGTATTGAAGGAGGATGGTAGGACGACAAAATAATTCATACAAAAA |
| 102_ALDH18A1 -_Seq_102 | GGGATGTATTGAAGGAGGATAAAAGGGGGAAGAATCCAAAGTATTAAAAT |
| 2_TES -_Seq_2 | GGGATGTATTGAAGGAGGATGGCTCCAAATCCTTGCTCGTGACCTAAGCC |
| 4_TES -_Seq_4 | GGGATGTATTGAAGGAGGATTACGACATATTTTTCTCCAGAAGTGCAGTT |
| 5_TES -_Seq_5 | GGGATGTATTGAAGGAGGATCATCATGCTCTTCTTGGCCACACTTGCAGT |
| 7_TES -_Seq_7 | GGGATGTATTGAAGGAGGATTATACTTGGTGTCTTCAAAAAGTTTTCCCA |
| 8_TES -_Seq_8 | GGGATGTATTGAAGGAGGATCATCTGACTTTAGTTTTGCAATCAGAGTGG |
| 10_TES -_Seq_10 | GGGATGTATTGAAGGAGGATTCTTCTTGGCAGCAACTGGATTCGTCAATA |
| 11_TES -_Seq_11 | GGGATGTATTGAAGGAGGATACTCATAGGTAACTGTATTGATGGAGACAT |
| 13_TES -_Seq_13 | GGGATGTATTGAAGGAGGATCCTTGGGTAGCATCTGCATGTACTGCCTGG |
| 14_TES -_Seq_14 | GGGATGTATTGAAGGAGGATCCCCCTCTGAGCCTGCTACTGGCTGCTTTT |
| 16_TES -_Seq_16 | GGGATGTATTGAAGGAGGATTTGAAGGGTCCTGGTCATGTGCAGGGAGCT |
| 17_TES -_Seq_17 | GGGATGTATTGAAGGAGGATTCACCTCTCTGGGAGACAACTCATGGCACT |
| 19_TES -_Seq_19 | GGGATGTATTGAAGGAGGATTGACATCTCCTACTCCCAGAGCTTCGCTCT |
| 20_TES -_Seq_20 | GGGATGTATTGAAGGAGGATGGCCTTGGGCATCCATCTCACAGGGAAGTT |
| 22_TES -_Seq_22 | GGGATGTATTGAAGGAGGATCCATGGCCCCCACTGCTGCTGGGGTGCTTC |
| 23_TES -_Seq_23 | GGGATGTATTGAAGGAGGATGAGTTCTTTTGTGCTCAGCAGATTTGTCCT |
| 25_TES -_Seq_25 | GGGATGTATTGAAGGAGGATCGGCATAGATGGCTGGGTCACCTTCTTTCA |
| 26_TES -_Seq_26 | GGGATGTATTGAAGGAGGATGGTGCCACAGTTTATCATAGCCAGCCCTTT |
| 28_TES -_Seq_28 | GGGATGTATTGAAGGAGGATTCCAAAAATAAATCATGTCAACCAGGAGTT |
| 29_TES -_Seq_29 | GGGATGTATTGAAGGAGGATAATGTCTGCCACAGTATAGCTTCTCATTCT |
| 31_TES -_Seq_31 | GGGATGTATTGAAGGAGGATACTCATTGCTGAATATCAGCTCGTCACAGC |
| 32_TES -_Seq_32 | GGGATGTATTGAAGGAGGATGGTGCCAATTCTGGTTTTCTGCCTGGGTAT |
| 34_TES -_Seq_34 | GGGATGTATTGAAGGAGGATCCATCACGTATATCTCCCCAGCTAGAATGC |
| 35_TES -_Seq_35 | GGGATGTATTGAAGGAGGATAGCAGGGCTTGCACACGGGCTTGTCATTGA |
| 37_TES -_Seq_37 | GGGATGTATTGAAGGAGGATGCACTTCTGGGTCGATGGCATTGTGGCATC |
| 39_TES -_Seq_39 | GGGATGTATTGAAGGAGGATAAGAGCACAGAAAGCACTCTGTGGATGCAT |
| 40_TES -_Seq_40 | GGGATGTATTGAAGGAGGATACTTCTGCCCAATGAGGCATTTGCTGCAGC |
| 42_TES -_Seq_42 | GGGATGTATTGAAGGAGGATCCTAAGACATCCTCTTCTTACATTCCACTG |
| 43_TES -_Seq_43 | GGGATGTATTGAAGGAGGATCTATGGCTCGATACTTCTGGGTGCCCTCCT |
| 45_TES -_Seq_45 | GGGATGTATTGAAGGAGGATATCTCTTGGTTTCCTTTACAGTTTCTTTTT |
| 46_TES -_Seq_46 | GGGATGTATTGAAGGAGGATAAACCAAGTTTAAAAGTTGAGACCGAAAAA |
| 48_TES -_Seq_48 | GGGATGTATTGAAGGAGGATTAATGAGATCCAAGCTTCCAAAAATGATTT |
| 49_TES -_Seq_49 | GGGATGTATTGAAGGAGGATGTGGCACAAATGGAATAGAGACATGAAGTT |
| 51_TES -_Seq_51 | GGGATGTATTGAAGGAGGATAAAAGAAGTGTAGAACTAGGAATGCTCATC |
| 52_TES -_Seq_52 | GGGATGTATTGAAGGAGGATTTTTCTTTTCATTTTACACATGAGGGGGAA |
| 54_TES -_Seq_54 | GGGATGTATTGAAGGAGGATAAAGGTCAGTGAATAAGACGAGGCAATAAA |
| 55_TES -_Seq_55 | GGGATGTATTGAAGGAGGATTGTCAAAAAGAATTCACTGTGTATCATTAC |
| 57_TES -_Seq_57 | GGGATGTATTGAAGGAGGATGAAAGAGTAAATGCATAGGCATTAAAACAG |
| 58_TES -_Seq_58 | GGGATGTATTGAAGGAGGATTAAAGAAAATGCCACCTCTGCCTAAATCAG |
| 60_TES -_Seq_60 | GGGATGTATTGAAGGAGGATGGCCACAGATGCAAGTCACCAACTTACAAA |
| 61_TES -_Seq_61 | GGGATGTATTGAAGGAGGATGATACTGTTATGCCACATGTGATAATAAAA |
| 63_TES -_Seq_65 | GGGATGTATTGAAGGAGGATGGCTAACATGGTGAAAACCTGTCTCTACTA |
| 64_TES -_Seq_68 | GGGATGTATTGAAGGAGGATCAGGCATGGTGGCTCCCACACCTGTAATCC |
| 66_TES -_Seq_70 | GGGATGTATTGAAGGAGGATGGTGGCTTGGCTTTTATGTCCAAGCCAAAA |
| 67_TES -_Seq_71 | GGGATGTATTGAAGGAGGATGGTACAAAAACTATACATGTTCTTTGGATT |
| 69_TES -_Seq_73 | GGGATGTATTGAAGGAGGATCCATTAGGAAAGAATAAAAACCATGCAAAA |
| 70_TES -_Seq_74 | GGGATGTATTGAAGGAGGATGACAGGCTTGGAAGTATTATAAAACAGGAT |
| 72_TES -_Seq_76 | GGGATGTATTGAAGGAGGATGAATAAACATTAGAAATACCTAGCCATGAA |
| 73_TES -_Seq_77 | GGGATGTATTGAAGGAGGATGCATAGCTATGTGTAGAAGTACACAGGGAA |
| 2_IDH1 ._Seq_3 | GGGATGTATTGAAGGAGGATGGAAAAATGAGTTTCTCTTTAATCAATTCC |
| 3_IDH1 ._Seq_4 | GGGATGTATTGAAGGAGGATTCATAGCTATGTAGATCCAATTCCACGTAG |
| 4_IDH1 ._Seq_5 | GGGATGTATTGAAGGAGGATTTGGTGGCATCACGATTCTCTATGCCTAAA |
| 5_IDH1 ._Seq_6 | GGGATGTATTGAAGGAGGATGCTTCTGCAGCATCCTTGGTGACTTGGTCG |
| 7_IDH1 ._Seq_8 | GGGATGTATTGAAGGAGGATACCCTCTTCTCATCAGGAGTGATAGTGGCA |
| 8_IDH1 ._Seq_10 | GGGATGTATTGAAGGAGGATAGAATATTTCGTATGGTGCCATTTGGTGAT |
| 9_IDH1 ._Seq_11 | GGGATGTATTGAAGGAGGATATAATGGCTTCTCTGAAGACCGTGCCACCC |
| 10_IDH1 ._Seq_12 | GGGATGTATTGAAGGAGGATCCACTCACAAGCCGGGGGATATTTTTGCAG |
| 12_IDH1 ._Seq_14 | GGGATGTATTGAAGGAGGATTCAGTTGCTCTGTATTGATCCCCATAAGCA |
| 13_IDH1 ._Seq_16 | GGGATGTATTGAAGGAGGATTGGGTTCCGTCACTTGGTGTGTAGGTTATC |
| 14_IDH1 ._Seq_17 | GGGATGTATTGAAGGAGGATTCAAAGTTATGTACCAGGTATGTCACCTTT |
| 15_IDH1 ._Seq_18 | GGGATGTATTGAAGGAGGATTACATCCCCATGGCAACACCACCACCTTCT |
| 16_IDH1 ._Seq_19 | GGGATGTATTGAAGGAGGATGTGCAAAATCTTCAATTGACTTATCTTGAT |
| 18_IDH1 ._Seq_21 | GGGATGTATTGAAGGAGGATTGTTTTTGGTGCTCAGATACAAAGGCCAAC |
| 19_IDH1 ._Seq_22 | GGGATGTATTGAAGGAGGATTAAAACGCCCATCATATTTCTTCAGAATAG |
| 20_IDH1 ._Seq_24 | GGGATGTATTGAAGGAGGATTCTTTTGAGCTTCAAACTGGGACTTGTACT |
| 21_IDH1 ._Seq_25 | GGGATGTATTGAAGGAGGATTGTCGTCGATGAGCCTATGCTCATACCAGA |
| 23_IDH1 ._Seq_28 | GGGATGTATTGAAGGAGGATCTTGGGCCACAGAGTCCGACTGCACGTCAC |
| 24_IDH1 ._Seq_29 | GGGATGTATTGAAGGAGGATCGCTGGTCATCATGCCGAGAGAGCCATACC |
| 25_IDH1 ._Seq_30 | GGGATGTATTGAAGGAGGATCTACTGTCTTGCCATCTGGACAAACCAGCA |
| 26_IDH1 ._Seq_31 | GGGATGTATTGAAGGAGGATGGGTTACAGTCCCGTGGGCAGCCTCTGCTT |
| 28_IDH1 ._Seq_33 | GGGATGTATTGAAGGAGGATAAATGGAAGCAATGGGATTGGTGGACGTCT |
| 29_IDH1 ._Seq_34 | GGGATGTATTGAAGGAGGATCTCTGTGGGCTAACCCTCTGGTCCAGGCAA |
| 30_IDH1 ._Seq_35 | GGGATGTATTGAAGGAGGATAGGCAAGCTCTTTATTGTTATCAAGCTTTG |
| 31_IDH1 ._Seq_36 | GGGATGTATTGAAGGAGGATTAGAGACTTCTTCCAAAGCATTTGCAAAGA |
| 32_IDH1 ._Seq_37 | GGGATGTATTGAAGGAGGATTGGTCATGAAGCCAGCCTCAATTGTCTCAA |
| 34_IDH1 ._Seq_40 | GGGATGTATTGAAGGAGGATTTTCTCCAAGTTTATCCATGAACTCAAATG |
| 35_IDH1 ._Seq_41 | GGGATGTATTGAAGGAGGATGTTTGGCCTGAGCTAGTTTGATCTTCAAGT |
| 36_IDH1 ._Seq_42 | GGGATGTATTGAAGGAGGATTATCCTTCTTAGCTCAGGTATGAACTTAAA |
| 37_IDH1 ._Seq_43 | GGGATGTATTGAAGGAGGATAACCTGTAGACCTAGTTACCAAAAGACAAT |
| 39_IDH1 ._Seq_45 | GGGATGTATTGAAGGAGGATCTCTGGCTTCTAAACAAATTACAAAATTGA |
| 40_IDH1 ._Seq_46 | GGGATGTATTGAAGGAGGATGCAATAACTGTATATATAAGAAAAAGGCTG |
| 41_IDH1 ._Seq_47 | GGGATGTATTGAAGGAGGATAAAAAGTCCCTTGCCATGTTCACAAAGGTG |
| 42_IDH1 ._Seq_48 | GGGATGTATTGAAGGAGGATCCTAGGCTGGTACTAGAAAATAAAATAAAA |
| 44_IDH1 ._Seq_50 | GGGATGTATTGAAGGAGGATCTTTTGAGCAATCTTGATTTTGGTCATTTA |
| 45_IDH1 ._Seq_51 | GGGATGTATTGAAGGAGGATTAGGGAGCAATACTGTGGCTATCATTTACC |
| 46_IDH1 ._Seq_52 | GGGATGTATTGAAGGAGGATAGGCAGTGAATTTCTACTTTATGCATATTT |
| 47_IDH1 ._Seq_53 | GGGATGTATTGAAGGAGGATCCTGTGCCCAAGGTCATGGACAGGAGGGGA |
| 48_IDH1 ._Seq_54 | GGGATGTATTGAAGGAGGATAAAACGGGATATCTATGACACCAGAACTTC |
| 50_IDH1 ._Seq_56 | GGGATGTATTGAAGGAGGATTGAGTTGCACTCCTACTTCGTTCCAGTCAT |
| 51_IDH1 ._Seq_57 | GGGATGTATTGAAGGAGGATCAAAAATGACTGCAGTATCTTCAACACATT |
| 52_IDH1 ._Seq_58 | GGGATGTATTGAAGGAGGATGTCTATTGGAAACATTCAGCAAGGTCTTTA |
| 53_IDH1 ._Seq_59 | GGGATGTATTGAAGGAGGATCAAACTCTCCTGCGGCCTAAACAGTATTTA |
| 55_IDH1 ._Seq_61 | GGGATGTATTGAAGGAGGATCAGGAGGAGAAAGAGAAAAATGGAGAGGAC |
| 56_IDH1 ._Seq_62 | GGGATGTATTGAAGGAGGATATTTAGAGTAGTATAATATTCAGGCCAGGC |
| 57_IDH1 ._Seq_63 | GGGATGTATTGAAGGAGGATTTACATTATTGCACTTGGATGAAATATGCT |
| 58_IDH1 ._Seq_64 | GGGATGTATTGAAGGAGGATAAATAAAACAGGCCAGCAGAAGTCCAAAAA |
| 59_IDH1 ._Seq_65 | GGGATGTATTGAAGGAGGATCAGGAGAAGATAAGATAGTGTTTAATATCA |
| 2_PLA2G4A ._Seq_2 | GGGATGTATTGAAGGAGGATAAACTTGTGGGAATACTGGTGCTCCACTAT |
| 4_PLA2G4A ._Seq_4 | GGGATGTATTGAAGGAGGATATCAAGCATGTCACCAAAGGCCCCCTTTGT |
| 6_PLA2G4A ._Seq_6 | GGGATGTATTGAAGGAGGATTGTTCTCTTCCTGCTGTCAGGGGTTGTAGA |
| 8_PLA2G4A ._Seq_8 | GGGATGTATTGAAGGAGGATATCCAAAATAAATTCAAAGGTCTCATTCCA |
| 9_PLA2G4A ._Seq_9 | GGGATGTATTGAAGGAGGATCGTAATCTCCAAAACATTTTCCTGATTAGG |
| 11_PLA2G4A ._Seq_11 | GGGATGTATTGAAGGAGGATAGATACAGTAAATGTTGCTGTCCCTAGAGT |
| 13_PLA2G4A ._Seq_13 | GGGATGTATTGAAGGAGGATCATTTCAGTGACTTGGTTGAAAATAAAAGG |
| 15_PLA2G4A ._Seq_15 | GGGATGTATTGAAGGAGGATCAGAGCCATACTAAATCGTAGGTCTGGGCA |
| 17_PLA2G4A ._Seq_17 | GGGATGTATTGAAGGAGGATCTTCATGCTCTCCCTTATGTGTTCTTTTCT |
| 18_PLA2G4A ._Seq_18 | GGGATGTATTGAAGGAGGATTCCTTCACTATTCTTTGGACCCAAGAGTTT |
| 20_PLA2G4A ._Seq_20 | GGGATGTATTGAAGGAGGATTCGGAAACCCCCACCTGAACCCAATATGGC |
| 22_PLA2G4A ._Seq_22 | GGGATGTATTGAAGGAGGATACAATCCAGAATTCCTGATTCGTATAATGC |
| 24_PLA2G4A ._Seq_24 | GGGATGTATTGAAGGAGGATGTGAGAATACAAGGTTGACATATACCAGGT |
| 26_PLA2G4A ._Seq_26 | GGGATGTATTGAAGGAGGATGTGGCTAACATTTTTCATTAGTTCTTCATT |
| 27_PLA2G4A ._Seq_27 | GGGATGTATTGAAGGAGGATTTTCTGTGGTGTGAGAAGTAAAAGGGGATT |
| 29_PLA2G4A ._Seq_29 | GGGATGTATTGAAGGAGGATAAAGGTGACAGGTTGTCCAGAGCTTTTCTT |
| 31_PLA2G4A ._Seq_31 | GGGATGTATTGAAGGAGGATGCTCAGAGTAGTATTCATTCTATTATGAAT |
| 33_PLA2G4A ._Seq_33 | GGGATGTATTGAAGGAGGATATGAAGACAGGTGAAAAGAGGTAAAGGGCA |
| 35_PLA2G4A ._Seq_35 | GGGATGTATTGAAGGAGGATTTCGTATGGACTAAATTCAACCCAATCTGC |
| 36_PLA2G4A ._Seq_36 | GGGATGTATTGAAGGAGGATCATAAAAGTACCATATTTAGCCATGCCAAT |
| 38_PLA2G4A ._Seq_38 | GGGATGTATTGAAGGAGGATCACCCATTAAGAAATGCAAGGGGTTTTCTT |
| 40_PLA2G4A ._Seq_40 | GGGATGTATTGAAGGAGGATTTTGTGAACCAGAAACGCCCAAAACTCTGT |
| 42_PLA2G4A ._Seq_42 | GGGATGTATTGAAGGAGGATCATTACTCACAATATGCTTTGTGGTAATAT |
| 43_PLA2G4A ._Seq_43 | GGGATGTATTGAAGGAGGATCGTGTGATTCATCATCACTGTCCGAGCTAT |
| 45_PLA2G4A ._Seq_45 | GGGATGTATTGAAGGAGGATTTGCTTGATTATCACTTTGATAGTCACTTC |
| 47_PLA2G4A ._Seq_47 | GGGATGTATTGAAGGAGGATCTCTGGTATTGAATAAAGCTGAATCACTCA |
| 49_PLA2G4A ._Seq_49 | GGGATGTATTGAAGGAGGATGATAAGATGTATTGAGATTCAAGCCCAGCA |
| 51_PLA2G4A ._Seq_51 | GGGATGTATTGAAGGAGGATCATCCAGTTCATCATCATCAAAGGAGTCCT |
| 52_PLA2G4A ._Seq_52 | GGGATGTATTGAAGGAGGATGCTCAAATTCATCAGGATCTGCTACAGCTG |
| 54_PLA2G4A ._Seq_54 | GGGATGTATTGAAGGAGGATTGAGCCCACTGTCCACTACATGAATCTTTT |
| 56_PLA2G4A ._Seq_56 | GGGATGTATTGAAGGAGGATAGATTATGAGATCAACCCCTCTCTGAGGTC |
| 58_PLA2G4A ._Seq_58 | GGGATGTATTGAAGGAGGATCAAGTAGAAGTTCCTTGAACGGAGGACTAG |
| 60_PLA2G4A ._Seq_60 | GGGATGTATTGAAGGAGGATCAAACACATAAGGATCAATCTTTGGAAAGG |
| 61_PLA2G4A ._Seq_61 | GGGATGTATTGAAGGAGGATAGACATAGCACTCCTTCAGCCCTTCCCGAT |
| 63_PLA2G4A ._Seq_63 | GGGATGTATTGAAGGAGGATCCAGAACAAAGTGGATGATGGTTGGGCAAT |
| 65_PLA2G4A ._Seq_65 | GGGATGTATTGAAGGAGGATTCTCTTCCTCAGTTTCCCTTGGAACACCTG |
| 67_PLA2G4A ._Seq_67 | GGGATGTATTGAAGGAGGATAATTGAAGGTTGAAAATGGTGATTCTGGGT |
| 69_PLA2G4A ._Seq_69 | GGGATGTATTGAAGGAGGATTCAGAGTATTGAAGTGCATAAGATCATGTA |
| 70_PLA2G4A ._Seq_70 | GGGATGTATTGAAGGAGGATCCATGGCTTCTTTTATCACATCAATGTTGT |
| 72_PLA2G4A ._Seq_72 | GGGATGTATTGAAGGAGGATCATTACTAAGGGAAACAGAGCAACGAGATG |
| 74_PLA2G4A ._Seq_74 | GGGATGTATTGAAGGAGGATACATGAACTATGCTTTGGGTTTACTTAGAA |
| 76_PLA2G4A ._Seq_76 | GGGATGTATTGAAGGAGGATAATCCAGTTGTCATGGGATTGCAAACTGCC |
| 78_PLA2G4A ._Seq_78 | GGGATGTATTGAAGGAGGATGCAACTTTGAGTATCAGCCAGTCTCTCATG |
| 79_PLA2G4A ._Seq_79 | GGGATGTATTGAAGGAGGATATAGTATTATTCTCATGCAGCTAAGTAACT |
| 81_PLA2G4A ._Seq_81 | GGGATGTATTGAAGGAGGATTGACTGAAAATGTAGCTAAGTATATCCTTA |
| 83_PLA2G4A ._Seq_83 | GGGATGTATTGAAGGAGGATCACATAAGAAAGTTGGTGAGAAATGTTTAA |
| 85_PLA2G4A ._Seq_85 | GGGATGTATTGAAGGAGGATTAGTCAGTGAATAACATTCATACATAATGC |
| 86_PLA2G4A ._Seq_86 | GGGATGTATTGAAGGAGGATAATAGTGTTGTCTCATGGTATGAATAAATC |
| 3_ACTN4 ._Seq_3 | GGGATGTATTGAAGGAGGATGTCGCCCATGCTGCCCCCGCCGCCAGCGCC |
| 5_ACTN4 ._Seq_6 | GGGATGTATTGAAGGAGGATGCACCATGCCGTGAAGGTCTTGCGCTGCTG |
| 8_ACTN4 ._Seq_9 | GGGATGTATTGAAGGAGGATGACCTCCAGGAGCAGCATGAGCTTGAGCCC |
| 10_ACTN4 ._Seq_11 | GGGATGTATTGAAGGAGGATGTTGATTTTGTGCACTCTCATCTTCCCCCG |
| 12_ACTN4 ._Seq_13 | GGGATGTATTGAAGGAGGATCCCGATGGAGACCAGCTTGACGCCTTTGCT |
| 15_ACTN4 ._Seq_16 | GGGATGTATTGAAGGAGGATCACGGAGATGTCCTGGATGGCGAACCTAAG |
| 17_ACTN4 ._Seq_18 | GGGATGTATTGAAGGAGGATATACGGGGCTGTCTTTCTCTGGCACCAGAG |
| 19_ACTN4 ._Seq_20 | GGGATGTATTGAAGGAGGATGGCATTGAAGGCAAGACCATCCTTCCAGCT |
| 22_ACTN4 ._Seq_23 | GGGATGTATTGAAGGAGGATCACTTCGAAGGCATTGTTCAGGTTGGTGAC |
| 24_ACTN4 ._Seq_25 | GGGATGTATTGAAGGAGGATGGCCGTGTTCACGATGTCCTCTGCATCCAG |
| 26_ACTN4 ._Seq_27 | GGGATGTATTGAAGGAGGATTCCTGAAAAGGCATGGTAGAAGCTGGACAC |
| 29_ACTN4 ._Seq_30 | GGGATGTATTGAAGGAGGATCTTCTCGTAGTCCTCCATCAGGTGCTCGTT |
| 31_ACTN4 ._Seq_32 | GGGATGTATTGAAGGAGGATCACACGGTCCTCCAGCCAGGGGATGGTGCG |
| 33_ACTN4 ._Seq_34 | GGGATGTATTGAAGGAGGATACGCCGGTAGTCGCGGAAGTCCTCCAGCTT |
| 36_ACTN4 ._Seq_39 | GGGATGTATTGAAGGAGGATTTCTCAGCCTGCTCCAAGTGCTGCCAGCCA |
| 38_ACTN4 ._Seq_41 | GGGATGTATTGAAGGAGGATTCTGCCAGGTGGTCGAGCCGCTCCAGCCTG |
| 41_ACTN4 ._Seq_45 | GGGATGTATTGAAGGAGGATTGCTTGCGAATGAGGGCTTTGATGTCCGAT |
| 43_ACTN4 ._Seq_47 | GGGATGTATTGAAGGAGGATGGGCAATGGCGGCGATCTGCTCCACGCGGT |
| 45_ACTN4 ._Seq_49 | GGGATGTATTGAAGGAGGATTCTTCTGGCACCGGGTGTTGACATTGTGGG |
| 48_ACTN4 ._Seq_52 | GGGATGTATTGAAGGAGGATGGTCGATGGCCTCCAGCTGCTTCTCTGTTT |
| 50_ACTN4 ._Seq_55 | GGGATGTATTGAAGGAGGATGGACGATGAACATGTCCTGGAGGTCCTCCA |
| 52_ACTN4 ._Seq_57 | GGGATGTATTGAAGGAGGATGCAGGGTGGACTTGAACTGGTCATGGGCTG |
| 55_ACTN4 ._Seq_61 | GGGATGTATTGAAGGAGGATTTTGCGGGGTGACGGTGGTGTAGGGGTTGC |
| 57_ACTN4 ._Seq_63 | GGGATGTATTGAAGGAGGATGGAGGGCATGGTCCCGTTTTGGCACCAGCT |
| 59_ACTN4 ._Seq_67 | GGGATGTATTGAAGGAGGATCCGTTCATCTCAATGGAGATGCGCCCGATC |
| 62_ACTN4 ._Seq_72 | GGGATGTATTGAAGGAGGATTGCTCCATGGTATAGTTGGTGTGCTTGTTG |
| 64_ACTN4 ._Seq_75 | GGGATGTATTGAAGGAGGATCCCTTGGCGTCGCGGGTGAGGATCTGGTTC |
| 66_ACTN4 ._Seq_78 | GGGATGTATTGAAGGAGGATCTTGAACTCCTCGGGCCCCAGCGCCCCGCC |
| 69_ACTN4 ._Seq_84 | GGGATGTATTGAAGGAGGATAGGAAGCGATGACCTGGTCAGCCGTGTCCG |
| 71_ACTN4 ._Seq_86 | GGGATGTATTGAAGGAGGATGCAGCTCTCTCCGCAGCTCCTCAGCTGTGA |
| 73_ACTN4 ._Seq_88 | GGGATGTATTGAAGGAGGATACTTGTAGTCGAGGGCACCGGGCACGGCGT |
| 76_ACTN4 ._Seq_91 | GGGATGTATTGAAGGAGGATGGCTGCCCAGGCCCCTCCTGGAGGCCGTCG |
| 78_ACTN4 ._Seq_93 | GGGATGTATTGAAGGAGGATACCCCGGAGGACTGCAGAGAGTGCTTTGCA |
| 81_ACTN4 ._Seq_96 | GGGATGTATTGAAGGAGGATCCCCAAGTCTCCCCAACCTGGTCGGGGGAA |
| 83_ACTN4 ._Seq_98 | GGGATGTATTGAAGGAGGATGGAATCCACTGGCCCCTCCTTGGTTAAAAA |
| 85_ACTN4 ._Seq_100 | GGGATGTATTGAAGGAGGATAGGAAGAGACCTGGGTGTGGTGAGGCATCC |
| 88_ACTN4 ._Seq_103 | GGGATGTATTGAAGGAGGATGCCCTCTGCTGGCCCCTCGCATGGCCCAGG |
| 90_ACTN4 ._Seq_105 | GGGATGTATTGAAGGAGGATGAGTCTGGAGCAGAGAGGGGAGAGGGGCTG |
| 92_ACTN4 ._Seq_107 | GGGATGTATTGAAGGAGGATTCAGCTCCTCTGCTGCAAAAGCGGGGTGCT |
| 95_ACTN4 ._Seq_129 | GGGATGTATTGAAGGAGGATCTTAGTGGATGGAGGTGGCCACCTTCTCCT |
| 97_ACTN4 ._Seq_131 | GGGATGTATTGAAGGAGGATCAGGGTGGCTCAGGCTCTGTCCTGCGAGCT |
| 99_ACTN4 ._Seq_134 | GGGATGTATTGAAGGAGGATCCTCTCTGCACAGGCCACTGCAGCCAGCAC |
| 102_ACTN4 ._Seq_137 | GGGATGTATTGAAGGAGGATTGTGGCCCACCTCACCTGCATCTTCTGTGA |
| 104_ACTN4 ._Seq_142 | GGGATGTATTGAAGGAGGATATGCAATGCCAAGGTGGGTTCCCTGTGGGC |
| 106_ACTN4 ._Seq_144 | GGGATGTATTGAAGGAGGATGCAGAGTTGGGGTGTGACACCCTGTGGCAT |
| 109_ACTN4 ._Seq_147 | GGGATGTATTGAAGGAGGATGGGTCGGGGCTGTCTTAGTCCGTTTGATGG |
| 111_ACTN4 ._Seq_149 | GGGATGTATTGAAGGAGGATCCACACATCGAGTCACGTGTCAGACGCTGT |
| 113_ACTN4 ._Seq_151 | GGGATGTATTGAAGGAGGATATATCCTCCAGTGCCCACTGGCAAGGAGAA |
